# Histone methyltransferase PRDM9 promotes survival of drug-tolerant persister cells in glioblastoma

**DOI:** 10.1101/2025.01.06.631591

**Authors:** George L. Joun, Emma G. Kempe, Brianna Chen, Jayden R. Sterling, Ramzi H. Abbassi, W. Daniel du Preez, Ariadna Recasens, Teleri Clark, Tian Y. Du, Jason K.K. Low, Hani Kim, Pengyi Yang, Jasmine Khor, Monira Hoque, Dinesh C. Indurthi, Mani Kuchibhotla, Ranjith Palanisamy, William T. Jorgensen, Andrew P. Montgomery, Jennifer R. Baker, Sarah L. Higginbottom, Eva Tomaskovic-Crook, Jeremy M. Crook, Lipin Loo, Bryan W. Day, G. Gregory Neely, Ernesto Guccione, Terrance G. Johns, Michael Kassiou, Anthony S. Don, Lenka Munoz

## Abstract

Chemotherapy often kills a large fraction of cancer cells but leaves behind a small population of drug- tolerant persister cells. These persister cells survive drug treatments through reversible, non-genetic mechanisms and cause tumour recurrence upon cessation of therapy. Here, we report a drug tolerance mechanism regulated by the germ-cell-specific H3K4 methyltransferase PRDM9. Through histone proteomic, transcriptomic, lipidomic, and ChIP-sequencing studies combined with CRISPR knockout and phenotypic drug screen, we identified that chemotherapy-induced PRDM9 upregulation promotes metabolic rewiring in glioblastoma stem cells, leading to chemotherapy tolerance. Mechanistically, PRDM9-dependent H3K4me3 at cholesterol biosynthesis genes enhances cholesterol biosynthesis, which persister cells rely on to maintain homeostasis under chemotherapy- induced oxidative stress and lipid peroxidation. PRDM9 inhibition, combined with chemotherapy, resulted in strong anti-cancer efficacy in preclinical glioblastoma models, significantly enhancing the magnitude and duration of the antitumor response by eliminating persisters. These findings demonstrate a previously unknown role of PRDM9 in promoting metabolic reprogramming that enables the survival of drug-tolerant persister cells.

## INTRODUCTION

Cancer therapy failure and tumour recurrence is, in part, attributed to a cancer cell subpopulation known as drug-tolerant persister cells.^1^ Unlike resistant cells, persister cells do not possess permanent resistance-conferring genetic mutations but instead enter a reversible drug-tolerant state.^2^ Upon cessation of therapy or in periods between therapy cycles (‘drug holiday’), persister cells give rise to a new population of cells that are often as sensitive to the drug as the original drug naïve population. Alternatively, overtime persister cells may acquire genetic mutations that confer irreversible drug resistance.^3^ The non-genetic mechanisms reported to underlie the drug-tolerant persister phenotype include epigenetic, transcriptomic, translational and metabolic reprogramming.^4^

Glioblastoma is the most common and lethal primary brain tumour. The DNA-alkylating agent temozolomide, albeit largely ineffective, remains the only approved drug for this disease.^5^ Several lines of pre-clinical evidence indicate that glioblastoma cells can be effectively targeted with microtubule-targeting agents (MTAs) which disrupt fundamental cell functions such as mitosis, migration, and vesicle transport.^6, 7, 8, 9^ Moreover, glioblastoma cells are connected via ultralong thin membrane protrusions known as tumour microtubes. Microtubes provide the anatomical basis for transfer of self-renewal factors between cells and their functioning depends on high content of microtubules.^10, 11^ However, clinical MTAs like taxanes and vinca alkaloids have limited therapeutic value in neuro-oncology due to their large molecular size and polarity which are incompatible with diffusion across the blood-brain barrier.^12^ Optical blood-brain-tumour barrier modulation to enhance paclitaxel delivery and the small-molecule MTA lisavanbulin have shown promise in glioblastoma models and trials.^13, 14,15^ These findings underscore the potential of brain-permeable MTA for glioblastoma therapy. Nonetheless, MTAs give rise to drug-tolerant persister cells.^16,8^ Thus, the identification of strategies to combat drug tolerance to MTAs holds the potential to enhance their efficacy, which is also critical for the clinical translation of brain-permeable MTAs.

PRDM9 (PR/SET domain 9) is a zinc finger protein that binds DNA and trimethylates histone H3 on lysine 4 and 36 to form H3K4me3 and H3K36me3. PRDM9 is normally expressed only in germ cells, where it regulates DNA recombination in meiosis.^17^ Following fertilisation, meiotic genes are silenced. Yet, the failure of meiotic gene-silencing is common in cancer, with over 200 cancer/testis genes, including PRDM9, found in tumours.^18, 19^ Pan-cancer analysis of 32 different cancer types (TCGA) revealed PRDM9 upregulation in 20% of tumours compared to healthy matching tissue, with the highest PRDM9 activity detected in glioblastoma.^19, 20^ PRDM9 in cancer has been linked to genomic instability, likely stemming from its known recombination function in meiosis.^19, 20^ However, despite PRDM9’s potential as an ideal and safe therapeutic target due to its germ-cell restricted expression, the functional role of PRDM9 in cancer is unknown.

Here, using a small-molecule microtubule-targeting agent CMPD1^21^ as the lead, we developed a novel brain-permeable microtubule-targeting agent WJA88. In pursuit of optimising the outcomes of MTA chemotherapy, we show that PRDM9’s methyltransferase activity is critical for the survival of glioblastoma persister cells. Mechanistically, PRDM9-mediated H3K4me3 maintains cholesterol homeostasis under chemotherapy stress. Moreover, inhibiting PRDM9 in combination with brain-permeable MTAs resulted in strong anti-glioblastoma efficacy and significantly reduced the population of persister cells, the main reason for tumour recurrence.

## RESULTS

### Glioblastoma persisters display transcriptomic and lipidomic reprogramming

MTAs induce apoptosis in glioblastoma cells^8, 21^, however do not kill all cells. Using the growth-rate (GR) metrics^22^, we have previously shown that small-molecule MTAs do not reach GR_max_ value of - 1, which denotes complete killing efficacy. Across 15 glioblastoma stem cell lines, RKI1 and FPW1 cells were least responsive to MTAs.^8^ Indeed, viable RKI1 and FPW1 cells can still be detected after 14 days of treatment with high concentrations (25x GR_50_) of CMPD1 or tivantinib (Fig. 1a). We have also shown that the size of RKI1 subpopulation surviving MTAs did not decrease when cells were co-treated with inhibitors of drug efflux pumps.^8^ In this study, we observed that surviving cells were morphologically different from the parent cells. As microtubule-targeting agents induce prominent changes in cell morphology, the observed shrinkage caused by disrupted microtubule cytoskeleton can be used as a marker of continuous target engagement in cells.^23, 24^

**Figure 1.**
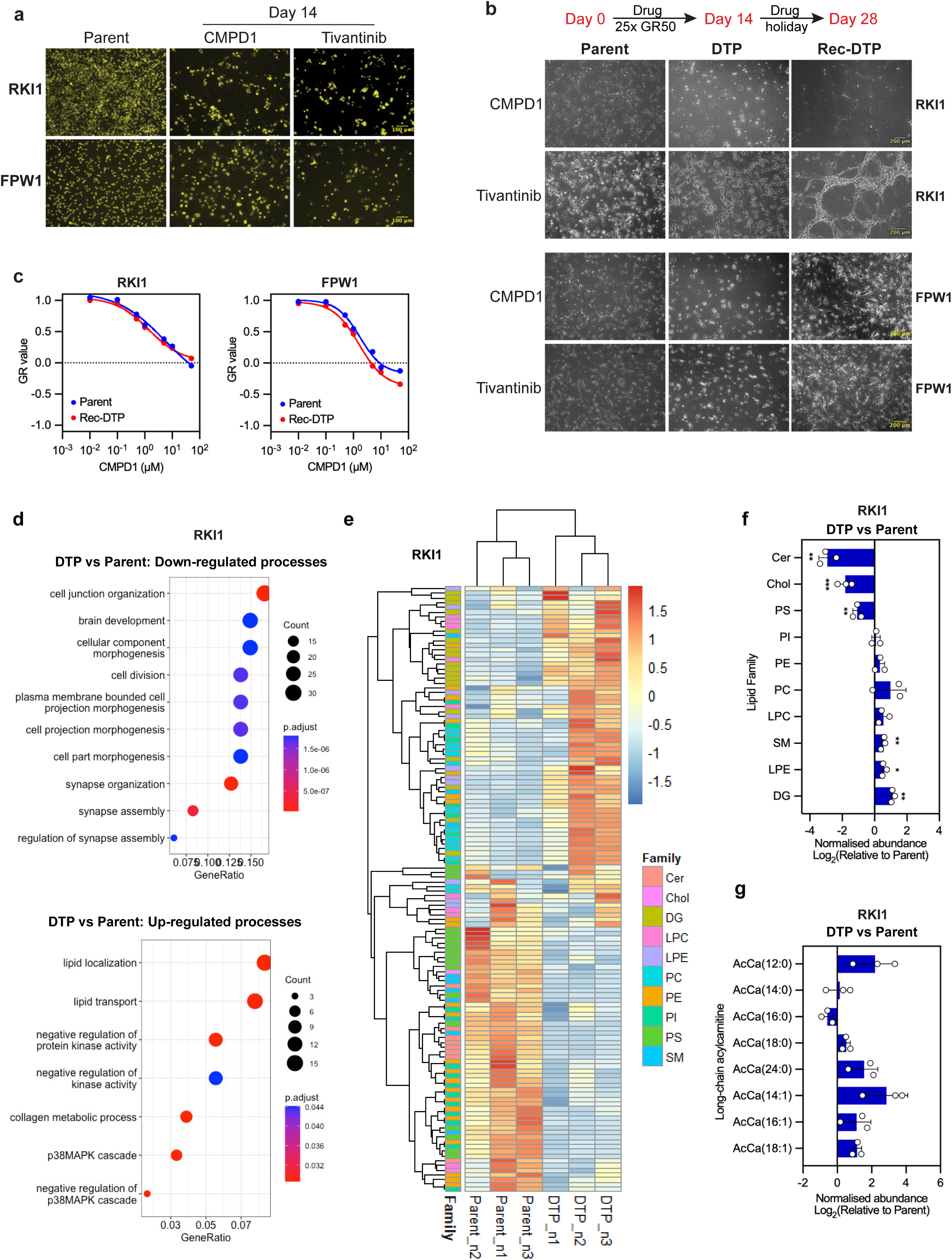
Characterisation of drug-tolerant persister cells in glioblastoma. a) Nuclear-ID red stain images (pseudo-coloured yellow) of RKI1 and FPW1 glioblastoma stem cells treated with CMPD1 or tivantinib (25 µM). Scale bar 100 µm. b) Schematic and brightfield images of parent (Day 0), drug-tolerant persister (DTP) and recovered DTP (Rec-DTP) cells generated with CMPD1 or tivantinib (25 µM), followed by recovery in drug- free media (drug holiday). Scale bar 200 µm. c) CMPD1 dose response curves in parent and recovered DTP cells. Data are mean of n = 3, corresponding GR metrics are in Supplementary Figure 1d. Rec-DTP cells were generated with CMPD1 (25 µM, 14 days), followed by drug holiday. d) Gene Ontology of top 200 down- and upregulated genes in CMPD1 (25 μM, 14 days) derived drug-tolerant persister cells compared to parent cells (RNA sequencing of n = 3). e) Heatmap of normalised lipids abundance in parent and CMPD1 (25 µM, 14 days) derived drug- tolerant persister cells (n = 3). f) Fold-change of lipid families in CMPD1 (25 μM, 14 days) derived drug-tolerant persister cells compared to parent cells. Data are mean ± SD (n = 3; multiple unpaired t-test). g) Fold-change of long-chain carnitines in CMPD1 (25 µM, 14 days) derived drug-tolerant persister cells compared to parent cells. Data are mean ± SD (n = 3; multiple unpaired t-test).

We used RT-qPCR to confirm that the CMPD1-surviving cells remained in a non-proliferative state throughout the treatment period, evidenced by a decrease in replication genes and upregulation of senescence and quiescence genes, particularly *CDKN1A* and p21 (Supplementary Fig. 1a-b). However, in drug holiday (*i.e.,* MTA withdrawal) surviving cells recovered their morphology, began to proliferate and re-colonized the wells within 14 days (Fig. 1b). Once a substantial population of recovered cells had been regenerated, we rechallenged these cells with CMPD1 or tivantinib. Consistent with a persister cell classification^2^, RKI1 and FPW1 persisters yielded MTA-sensitive progeny (Fig. 1c; Supplementary Fig. 1c-d). These observations suggest that we did not select out a subclone with a pre-existing resistance mutation and the fractional killing efficacy of CMPD1 and tivantinib leads to drug-tolerant persisters.

To investigate persisters in an unbiased manner, we compared the transcriptomes of parent and CMPD1-derived RKI1 persister cells by bulk RNA sequencing (RNA-seq). We identified 1,573 differentially expressed genes (DEGs, *P_adj_* < 0.01; Supplementary Fig. 1e). Gene Ontology of the top 200 DEGs identified that the down-regulated processes were related to cell division and morphogenesis, whereas lipid localisation and transport were the most up-regulated processes (Fig. 1d). Following up on these findings, we found a distinctive lipidomic signature in CMPD1-derived persisters compared to parent cells (Fig. 1e). Significant treatment-related decreases occurred with ceramide (Cer), cholesterol (Chol) and phosphatidylserine (PS), whereas sphingomyelin (SM), lysophosphatidylethanolamine (LPE), and diacylglycerol (DG) increased in persisters (Fig. 1f). Furthermore, carnitine-linked fatty acids, which are substrates for mitochondrial β-oxidation, were more abundant in persisters (Fig. 1g), in line with data showing the dependence of melanoma persisters on fatty acid oxidation.^25^

### Glioblastoma persisters exhibit increased histone lysine methylation

Epigenetic reprogramming is considered as a critical mechanism driving the reversible drug-tolerant state^4^ that we observed in glioblastoma persisters (Fig. 1b-c). We therefore profiled histone extracts of parent and CMPD1-derived persister cells by mass-spectrometry (Supplementary Fig. 2a) and constructed H3-centric peptide heatmaps (Fig. 2a, Supplementary Fig. 2b). This comprehensive analysis revealed deposition of replication-independent histone variant H3.3 compared to canonical H3.1/H3.2 in persisters (Fig. 2b), confirmed with immunoblotting (Fig. 2c). H3.3 preferentially occurs at active gene bodies and has been associated with aggressive metastasis of carcinomas.^26^ Further, we observed significant deacetylation occurring on K27 and K36 (Fig. 2d). In line, H3K27me3 and H3K36me3 were the most prominent modifications in persisters (Fig. 2e-f). There was no significant modification of the H3K4 and H3K9 methylation marks, except for H3K4me3 levels being consistently low in RKI1 persisters (Fig. 2f).

**Figure 2.**
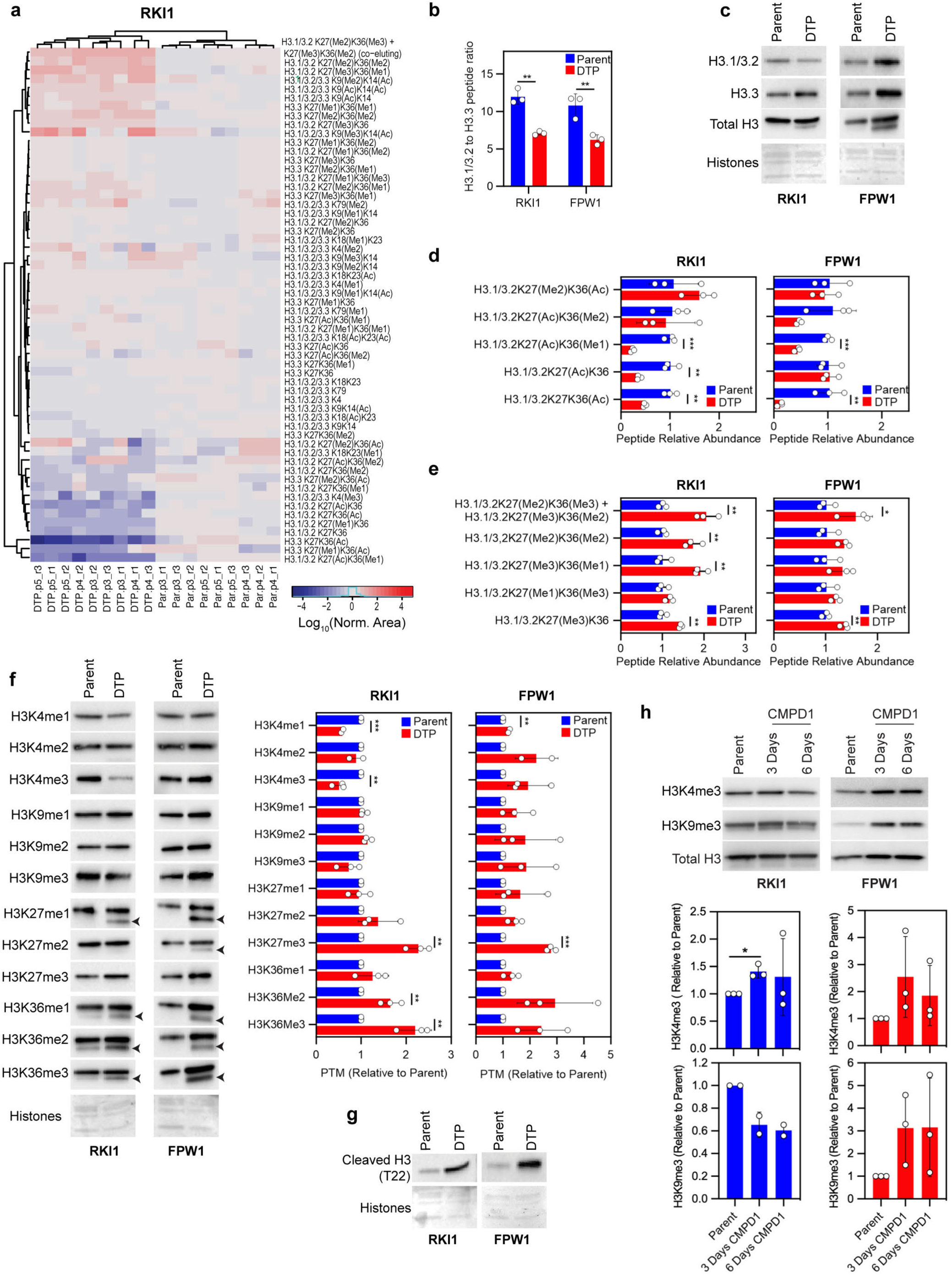
Increased H3 methylation in drug-tolerant persister cells. a) Heatmap of peak areas for H3 peptides in parent (Par) and CMPD1 (25 μM, 14 days) derived drug- tolerant persister (DTP) cells. Data are parent-DTP pairs of 3 independent experiments performed in triplicate. b) The sum of H3.1/3.2K27K36 peptides peak area was divided by the sum of H3.3K27K36 peptides peak area within each sample. Data are mean ± SD (n = 3; multiple unpaired t-tests) c) Immunoblots of total H3, H3.1/H3.2 and H3.3 variants in parent and CMPD1 (25 μM, 14 days) derived drug-tolerant persister cells. Equal loading was monitored with amido black B membrane stain (bottom panel). d - e) Relative abundance of H3.1/3.2K27K36 peptides in parent and CMPD1 (25 μM, 14 days) derived drug-tolerant persister cells. Data are mean ± SD (n = 3; multiple unpaired t-test). f - g) Immunoblots and quantification of H3 lysine methylation and cleavage in parent and CMPD1 (25 µM, 14 days) derived drug-tolerant persister cells. Data are mean ± SD (n = 3; one sample t- tests). Equal loading was monitored with amido black B membrane stain (bottom panel). h) Immunoblots and quantification of H3K4me3, H3K9me3 and total H3 in cells treated with CMPD1 (25 µM) for 3 and 6 days. Data are mean ± SD (n = 3; one sample t-tests).

Interestingly, immunoblotting of more distal H3K27 and H3K36 methylation marks revealed an additional faster migrating species of H3 in persisters that was absent in parent cells (Fig. 2f, arrows). This was not detected with antibodies specific for H3K4 and H3K9. Given that H3.3 variant is cleaved during senescence^27^, generating a new H3.3 tail beginning with the amino acid T22; and that persisters had elevated H3.3 deposition (Fig. 2b-c) as well as expression of senescence associated markers (Supplementary Fig. 1a-b), we used H3 cleavage-specific antibody to confirm H3 cleavage in persisters (Fig. 2g). Further, we demonstrate that CMPD1 treatment of RKI1 cells increased H3K4me3 at Day 3; and the signal dropped by Day 6, while H3K9me3 levels were reduced at Day 3 and 6 (Fig. 2h). In FPW1 cells, we observed increased H3K4me3, H3K9me3 and total H3 levels at Day 3 and Day 6. These findings establish that global H3 methylation occurs during the onset of drug tolerance, with proteolytic cleavage of histone tails removing H3K4 and H3K9 methyl marks later during the treatment. The deposition of H3.3, coupled with methylation of gene-activating (H3K4, H3K36) as well as gene-repressing (H3K9, H3K27) residues correlates with the extensive transcriptional changes observed in persisters (Fig. 1d; Supplementary Fig. 1e).

### PRDM9 inhibition eliminates glioblastoma persisters

Given the significant chromatin remodelling in persister cells, we assessed the sensitivity of MTA- tolerant cells to 37 epigenetic chemical probes (*i.e.,* well-characterised inhibitors).^28^ Structurally related KMT6 (aka EZH2) inhibitors UNC-1999 and GSK-343 attenuated viability of persisters without changing the viability of the parent cells (Supplementary Fig. 3a). However, GR curves for orthogonal KMT6 inhibitors UNC-1999, GSK-126, CPI-169, CPI-1205 and tazemetostat revealed minimally increased sensitivity of persisters to KMT6 inhibitors compared to parent cells (Supplementary Fig. 3b). GSK-J4 and KDOBA-67a (targeting KDM6) and JQ1 (targeting BRD4) reduced viability of both parent and persister cells (Supplementary Fig. 3a), consistent with previous findings^29^ and suggesting that they do not target a mechanism specific to MTA tolerance.

In the viability screen (Supplementary Fig. 3a), we introduced epigenetic probes to persisters after a 14-days treatment with CMPD1 or tivantinib. Considering that H3K4 and H3K9 methylation occurs immediately after CMPD1 treatment but is then removed by proteolytic cleavage (Fig. 2h), we co-treated cells with CMPD1 and UNC1999, KDOBA-67a, GSK-J4, or MRK-740 for 14 days and quantified persisters (Fig. 3a). Notably, MRK-740, an inhibitor of H3K4 methyltransferase PRDM9^30^, significantly reduced the number of RKI1 persisters (Fig. 3a-b). To further investigate the efficacy of MRK-740, four glioblastoma stem cell lines, each with distinct combinations of genetic mutations^31^ were co-treated with CMPD1 and either MRK-740 or matched inactive compound MRK- 740-NC which does not inhibit PRDM9^30^, followed by drug holiday (Fig. 3c-d). In addition to its efficacy in RKI1 cells, MRK-740 reduced the colony area of recovered persisters in HW1, FPW1 and SB2B cells (Fig. 3d). Of note, single MRK-740 inhibited cell proliferation while present in the cell culture medium (Supplementary Fig. 3c), but cells fully recovered during the drug holiday (Fig. 3c). MRK-740-NC had no effect on cell viability and did not reduce the number of persister-derived colonies (Fig. 3c-d). To confirm PRDM9 inhibition in cells, we demonstrate that MRK-740 (3 µM), but not MRK-740-NC, decreased bulk H3K4me3 in RKI1 cells treated with CMPD1 without reducing H3K4me3 when used as a single agent (Fig. 3e).

**Figure 3.**
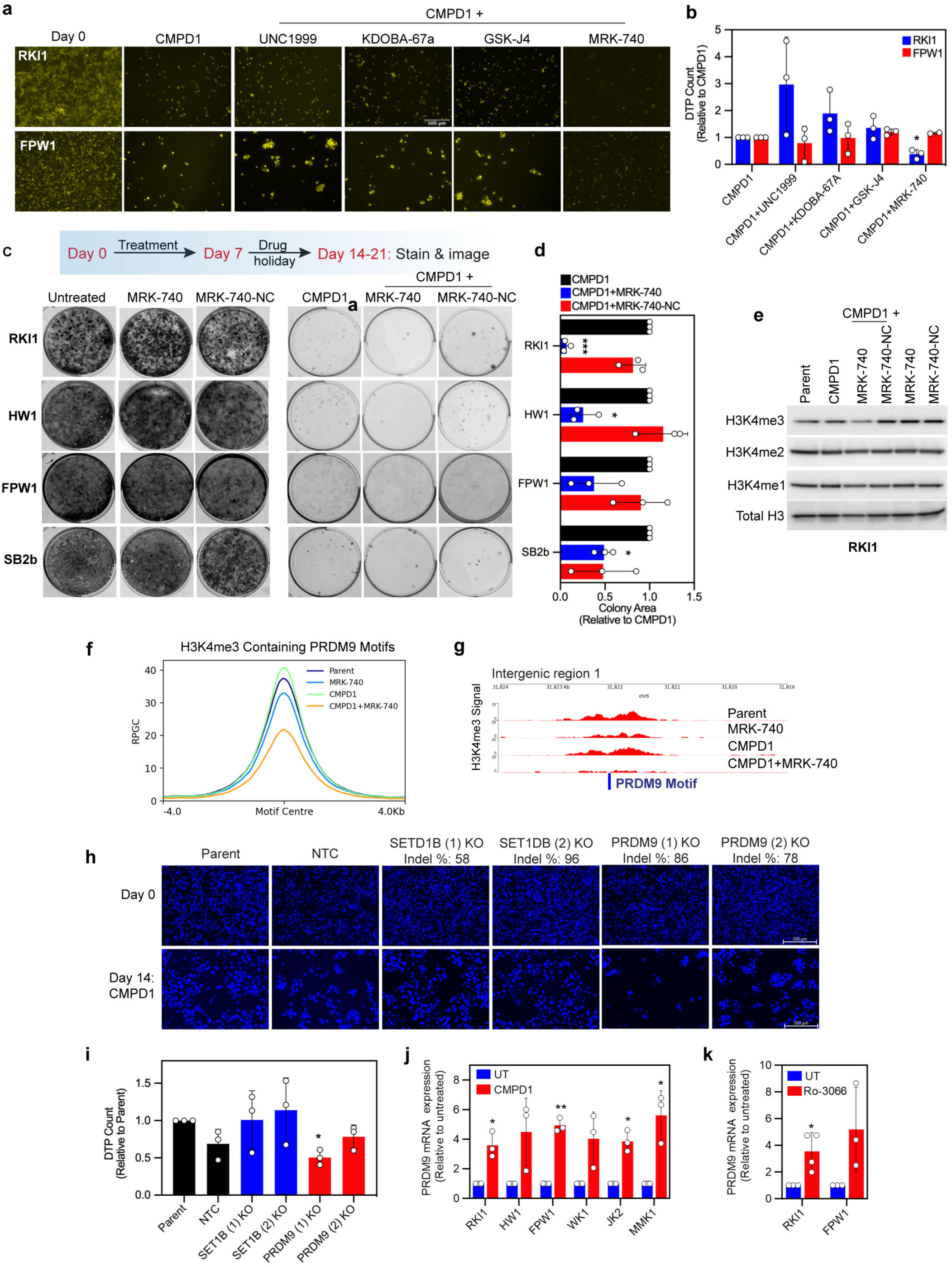
PRDM9 activity is required for survival of drug-tolerant persisters. a - b) Nuclear-ID red stain images (pseudo-coloured yellow) and quantification of drug-tolerant persister (DTP) cells surviving CMPD1 (25 µM, 14 days) treatment combined with UNC1999 (3 μM), KDOBA-67a (5 µM), GSK-J4 (5 µM) and MRK-740 (3 µM). Data are mean ± SD (n = 3; one sample t-test) c - d) Representative images and quantification of colony formation assay with glioblastoma stem cell lines treated with MRK-740 (3 µM), MRK-740-NC (3 µM) alone and in combination with CMPD1 (25 µM) for 7 days, followed by drug holiday until visible colonies formed in CMPD1-only treatments. Data are mean ± SD (n = 3; one sample t-test) e) Immunoblots of H3K4 methylation in RKI1 cells treated with CMPD1 (10 µM) ± MRK-740 (3 µM) ± MRK-740-NC (3 µM) for 3 days. f) Combined intensity (RPGC) of H3K4me3 peaks aligning with a PRDM9 motif in RKI1 cells treated with CMPD1 (10 µM) ± MRK-740 (3 µM) for 3 days. g) Individual genome track of H3K4me3 intensity (RPGC) at an intergenic region aligning with PRDM9 motif (blue bar) in RKI1 cells treated with CMPD1 (10 µM) ± MRK-740 (3 µM) for 3 days. h - i) DAPI-stained images and quantification of RKI1 cells following SETD1B and PRDM9 knock- out (KO) and treated with CMPD1 (25 µM) for 14 days. Data are mean ± SD (n = 3; one sample t-test). NTC: non-targeted control. j) RT-qPCR analysis of PRDM9 mRNA in glioblastoma stem cell lines treated with CMPD1 (25 μM) for 72 h. Data are mean ± SD (n = 3; one sample t-test). k) RT-qPCR analysis of PRDM9 mRNA in RKI1 and FPW1 cells treated with Ro-3306 (10 μM) for 72 h. Data are mean ± SD (n = 3; one sample t-test).

To further validate PRDM9 inhibition by MRK-740 in cells and exclude inhibition of other H3K4 methyltransferases, we performed H3K4me3 ChIP-Seq (chromatin immunoprecipitation followed by next-generation sequencing) analyses of RKI1 cells treated with CMPD1 ± MRK-740. The intensity of 27,749 H3K4me3 peaks (q-value < 0.05, Supplementary Table 1) increased in cells treated with CMPD1, while MRK-740, alone or combined with CMPD1, reduced the signal (Supplementary Fig. 3d). Genomic regions with the most significant PRDM9 motif matches (P < 2.67x10^-7^, MA1723.2, JASPAR database, Supplementary Table 2) identified that 3,872 out of 27,749 H3K4me3 peaks overlap with at least one PRDM9 motif (Supplementary Table 3). Plotting these 3,872 peaks confirmed alignment with the PRDM9 motif center and reduced H3K4me3 signal in cells treated with MRK-740 (Fig. 3f). Since over 80% of these peaks are located at promoters (Supplementary Fig. 3e) where H3K4me3 are also formed by the COMPASS complex^32^, H3K4me3 peaks at distal intergenic sites can be attributed solely to PRDM9 activity.^30^ We found near complete eradication of H3K4me3 at intergenic regions in cells treated with MRK-740 (Fig. 3g, Supplementary Fig. 3f). These data unequivocally confirm inhibition of PRDM9’s methyltransferase activity by MRK-740 in RKI1 cells.

Next, we knocked out PRDM9 in RKI1 cells to test whether loss of PRDM9 phenocopies the anti-persister efficacy of MRK-740. As controls, we included H3K4 methyltransferases SETD1A and SETD1B, members of the COMPASS complex.^32^ DepMap database revealed that SETD1A is a common essential gene (1,070/1,095 cell lines with Chronos score ≤ -0.5), and 154 cell lines showed probability of SETD1B dependency. However, no cell lines had a probability of PRDM9 dependency (Supplementary Fig. 3g), aligning with cytostatic efficacy of MRK-740 in RKI1 cells (Supplementary Fig. 3c). As expected, SETD1A knockout was detrimental to RKI1 cell viability (data not shown). Knocking out SETD1B and PRDM9 had no impact on RKI1 cell fitness, suggesting that these genes are not essential in unchallenged cells (Fig. 3h). However, when these cells were treated with CMPD1 for 14 days, PRDM9 knockout decreased the number of persister cells compared to parent cells, whereas SETD1B knock-out had no effect (Fig. 3h-i).

PRDM9 protein levels are very low, even in germ cells, with its expression being transient and restricted to the prophase, the first phase of meiosis.^33^ Using commercially available antibodies we were unable to assess PRDM9 protein levels in glioblastoma cells, but detected increased PRDM9 mRNA in several glioblastoma cells treated with CMPD1 for 72 hrs (Fig. 3j). Given that microtubule- targeting agents, including CMPD1 induce mitotic arrest^21^, we reasoned that PRDM9 upregulation could results from the mitotic arrest induced by CMPD1. To test this hypothesis, we used CDK1 inhibitor Ro-3306 to synchronise cells to mitosis^34^ and found a significant Ro-3306 induced PRDM9 mRNA upregulation (Fig. 3k). Together, these findings suggest that while PRDM9 is not an essential gene in glioblastoma, its expression increases in mitotically arrested cells and the development of a drug-tolerant state during chemotherapy is contingent on this methyltransferase.

### PRDM9-dependent H3K4me3 regulates cholesterol biosynthesis genes

To unravel the mechanism by which PRDM9 promotes survival of drug-tolerant persisters, we performed RNA sequencing (RNAseq) of our pharmacological and genetic PRDM9 inhibition models (MRK-740 and PRDM9 knockout) head-to-head. RKI1 cells were treated with CMPD1 ± MRK-740 for 3 days and harvested for analysis. In parallel, RKI1 cells transduced with NTC sgRNA (referred to as NTC cells) and PRDM9(1) sgRNA (referred to as PRDM9 KO cells) were analysed after 3 days treatment with CMPD1. Given that H3K4me3 is a transcription-activating chromatin mark^35^ and aiming to identify a process exclusive to CMPD1+MRK-740 treatment, which is absent in CMPD1 monotherapy and thus likely to underscore MRK-740-induced persisters’ death, we focused on top 50 down-regulated genes (down-DEGs, P_adj_ < 0.05) in cells treated with CMPD1+MRK-740 versus CMPD1-only cells (Fig. 4a). Gene Ontology (GO) mapped these down-DEGs to cholesterol biosynthesis (Fig. 4b). In line, transcripts for numerous enzymes involved in cholesterol biosynthesis, including HMGCR, HMGCS1, MVD, EBP, SQLE, DHCR7 and DHCR24, were down-regulated upon MRK-740 treatment (Fig. 4a). Mapping the 50 down-DEGs from Fig. 4a on the waterfall plot ranking differentially expressed genes in PRDM9 KO cells identified DHCR24, DHCR7, EBP and SEMA7A transcripts as most significantly impacted by PRDM9 loss (Fig. 4c).

**Figure 4.**
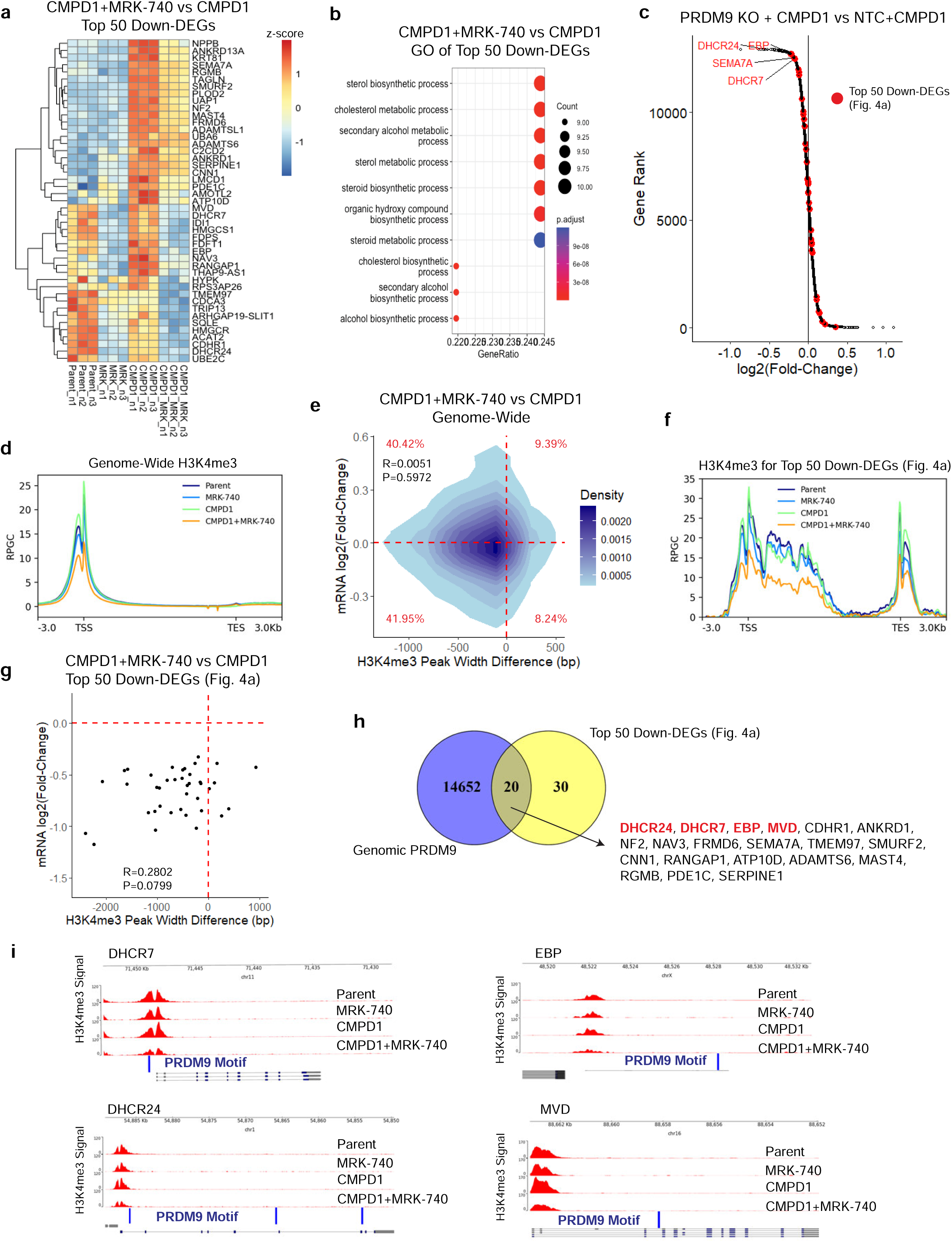
PRDM9-dependent H3K4me3 marks cholesterol biosynthesis genes. a - b) Heatmap and Gene Ontology of top 50 down-regulated DEGs (RNA sequencing of n = 3) in RKI1 cells treated with CMPD1 (10 µM) ± MRK-740 (3 µM) for 3 days. c) Waterfall plot of differentially expressed genes (RNA sequencing of n = 3) in CMPD1 (10 µM, 3 days) treated RKI1 cells transduced with NTC sgRNA (NTC) or PRDM9 sgRNA (PRDM9 KO). d) Genome wide H3K4me3 signal intensity (RPGC) (± 3kb from transcription start/end site) in RKI1 cells treated with CMPD1 (10 µM) ± MRK-740 (3 µM) for 3 days. e) 2D kernel density plot showing the relationship between changes in H3K4me3 peak width (x-axis) and mRNA expression in RKI1 cells treated with CMPD1 (10 µM) ± MRK-740 (3µM) for 3 days. Each data point corresponds to a gene that was detected by RNA sequencing and contained a H3K4me3 peak with > 1000 bp in CMPD1-treated cells. The colour bar reflects the density. Pearson’s product-moment correlation was used to calculate the correlation coefficient (R) and statistical significance (P). f) H3K4me3 signal intensity (RPGC) for down-regulated genes (listed in Figure 4a; ± 3kb from transcription start/end site) in RKI1 cells treated with CMPD1 (10 µM) ± MRK-740 (3 µM) for 3 days. g) Scatter plot of top down-DEGs listed in Figure 4a and containing a H3K4me3 peak > 1000 bp in CMPD1-treated cells. The plot shows correlation between changes in H3K4me3 peak width (x-axis) and mRNA expression in RKI1 cells treated with CMPD1 (10 µM) ± MRK-740 (3 µM) for 3 days. Pearson’s product-moment correlation was used to calculate the correlation coefficient (R) and statistical significance (P). h) Venn diagram of functional genomic PRDM9 annotated regions (Homo Sapiens PRDM9 matrix; Supplementary Table 2) and down-regulated genes listed in Figure 4a. i) H3K4me3 genome tracks for cholesterol biosynthesis genes *DHCR7*, *DHCR24*, *EBP* and *MVD* in RKI1 cells treated with CMPD1 (10 µM) ± MRK-740 (3 µM) for 3 days.

Next, we reanalysed our H3K4me3 ChIP-seq datasets obtained from RKI1 cells treated with CMPD1 ± MRK-740 to determine whether PRDM9-dependent H3K4me3 peaks localise to cholesterol biosynthesis genes and regulate their transcription. First, we confirmed that the positive correlation between H3K4me3 peak width, which has been found instructive for gene transcription^35^, and transcripts levels remained constant during all treatments (Supplementary Fig. 4a). Using the K-means clustering of H3K4me3 peak intensities, we divided genes in untreated RKI1 cells into Cluster 1 (genes with H3K4me3) and Cluster 2 (genes without H3K4me3; Supplementary Fig. 4b). Cluster 1 genes had significantly higher mRNA expression compared to Cluster 2 genes (Supplementary Fig. 4c). Additionally, when treated with CMPD1, Cluster 1 genes were kept to a tighter expression range compared to Cluster 2 genes (Supplementary Fig. 4d). This confirms that H3K4me3 maintains transcriptional consistency in RKI1 cells, as reported previously.^35^

Genome-wide analysis revealed that H3K4me3 peak intensity was highest at the promoters in CMPD1-treated cells and lowest in cells co-treated with CMPD1 and MRK-740 (Fig. 4d, Supplementary Fig. 4e-f). By plotting changes in H3K4me3 peak width for individual genes against changes in mRNA expression, we found no correlation (Fig. 4e). This suggests that not all genes’ transcription is sensitive to MRK-740 induced changes in H3K4me3 peak width. However, decreased H3K4me3 intensity (Fig. 4f) and peak width (Fig. 4g) for the top 50 down-DEGs more strongly correlated with decreased mRNA levels (Fig. 4g), indicating that the transcription of cholesterol biosynthesis genes is particularly sensitive to MRK-740 treatment. To identify PRDM9-dependent H3K4me3 marks, a Venn diagram of 14,652 genes annotations with a PRDM9 motif and the 50 down-DEGs revealed that 20 genes, including cholesterol biosynthesis genes *DHCR7*, *DHCR24*, *EBP*, and *MVD*, contain the PRDM9 binding motif (Fig. 4h). The remaining 30 genes, despite lacking the PRDM9 motif, could be methylated by PRDM9 via its KRAB domain interactions with CXXC1, a COMPASS complex subunit.^36, 37^ Individual genome tracks confirmed that *DHCR7* and *DHCR24* have PRDM9 motifs at their promoters, while *EBP* and *MVD* contain an intronic PRDM9 motif (Fig. 4i). Importantly, reduced H3K4me3 intensity at these genes’ promoters was found in cells co-treated with CMPD1 and MRK-740.

To functionally support that the observed effects of MRK-740 are truly PRDM9-dependent, we performed H3K4me3 ChIP-Seq of NTC and PRDM9 KO cells treated with CMPD1 for 3 days. We found that CMPD1 reduced total H3K4me3 in NTC cells (Supplementary Fig. 4g), an effect not seen in un-transduced RKI1 cells treated with CMPD1 (Supplementary Fig. 3d). Notably, long-term CMPD1 treatment (14 days) of NTC cells resulted in fewer persisters compared to un-transduced RKI1 cells (Fig. 3i). Although the reason for the decreased H3K4me3 in CRISPR-Cas9 modified NTC cells remains unclear, this finding supports a link between H3K4me3 intensity and persister cell survival. Nevertheless, the intensity of 23,073 H3K4me3 peaks (q-value < 0.05, Supplementary Table 4, Supplementary Fig. 4g); genome-wide H3K4me3 peaks (Figure 5a); 1,905 H3K4me3 peaks overlapping with at least one PRDM9 motif (Fig. 5b, Supplementary Table 5); and individual H3K4me3 peaks at intergenic regions overlapping with a PRDM9 motif (Fig. 5c, Supplementary Fig. 4h) was reduced in PRDM9 knockout (KO) cells, especially when treated with CMPD1. Genome tracks of cholesterol genes *DHCR7*, *DHCR24*, *EBP*, and *MVD* confirmed most significantly reduced H3K4me3 in PRDM9 KO cells treated with CMPD1 (Fig. 5d). Overall, CMPD1 treatment combined with PRDM9 loss phenocopied the results obtained with CMPD1+MRK-740.

**Figure 5.**
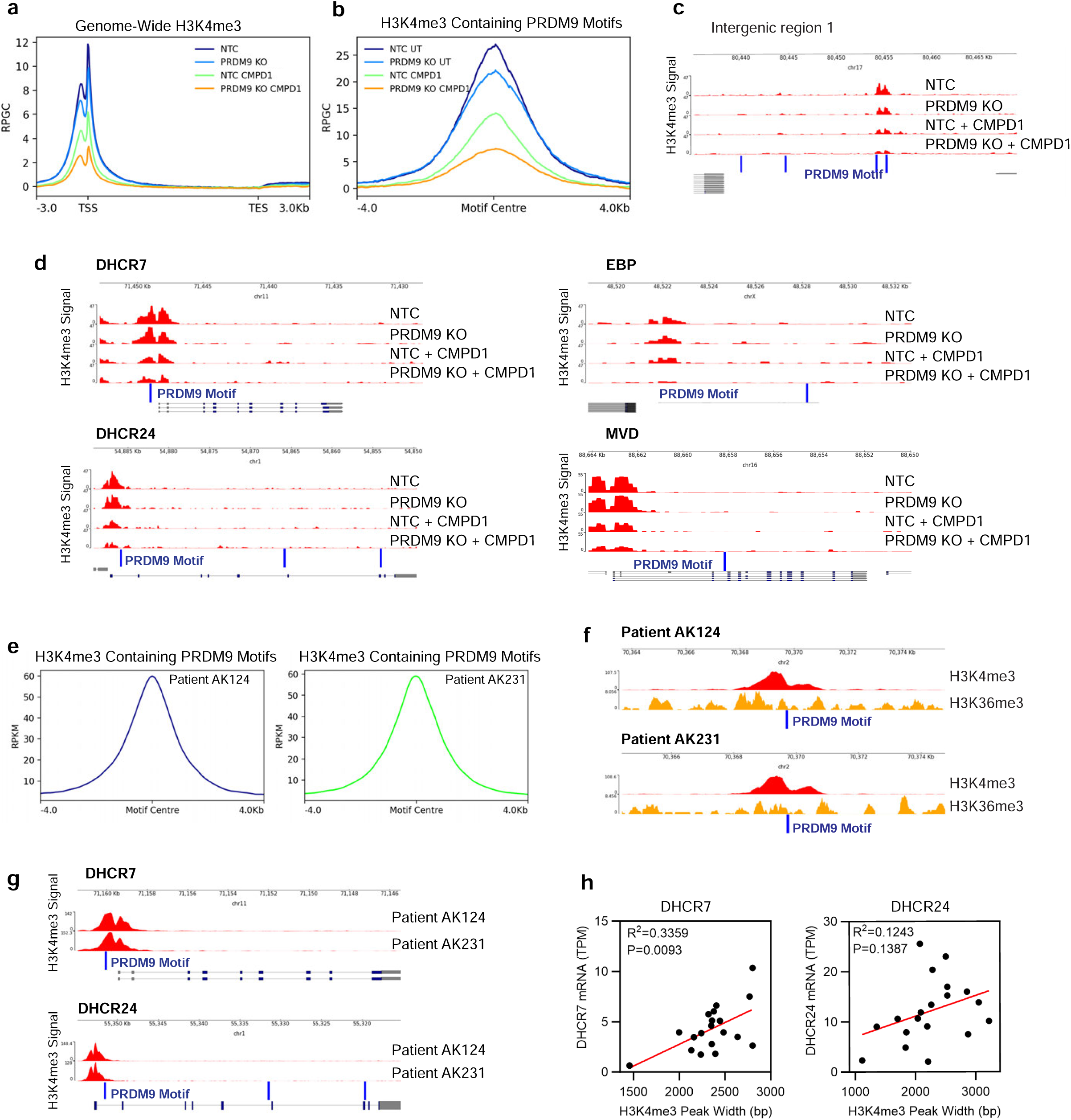
H3K4me3 regulates transcription of cholesterol biosynthesis genes in glioblastoma. a) Genome wide H3K4me3 ChIPseq signal intensity (RPGC) (± 3kb from transcription start/end site) in CMPD1 (10 µM, 3 days) treated RKI1 cells transduced with NTC sgRNA (NTC) or PRDM9 sgRNA (PRDM9 KO). b) Combined intensity (RPGC) of H3K4me3 peaks aligning with a PRDM9 motif in CMPD1 (10 µM, 3 days) treated RKI1 cells transduced with NTC sgRNA (NTC) or PRDM9 sgRNA (PRDM9 KO). c) Individual genome track of H3K4me3 intensity (RPGC) at an intergenic region aligning with PRDM9 motif (blue bar) in CMPD1 (10 µM, 3 days) treated RKI1 cells transduced with NTC sgRNA (NTC) or PRDM9 sgRNA (PRDM9 KO). d) H3K4me3 genome tracks for cholesterol biosynthesis genes *DHCR7*, *DHCR24*, *EBP* and *MVD* in CMPD1 (10 µM, 3 days) treated RKI1 cells transduced with NTC sgRNA (NTC) or PRDM9 sgRNA (PRDM9 KO). e) Combined intensity (RPKM) of H3K4me3 peaks aligning with a PRDM9 motif in glioblastoma patient AK124 and AK231 (GSE121723). f) H3K4me3 and H3K36me3 genome tracks at intergenic regions aligning with a PRDM9 motif in glioblastoma patients AK124 and AK231 (GSE121723). g) H3K4me3 genome tracks for cholesterol biosynthesis genes *DHCR7* and *DHCR24* in glioblastoma patients AK124 and AK231 (GSE121723). h) Pearson’s correlation showing relationship between H3K4me3 peak widths and mRNA expression for DHCR7 and DHCR24 in 19 glioblastoma patients (GSE121723).

Our results indicate that PRDM9 regulates H3K4me3 and transcription at cholesterol biosynthesis genes. To investigate the clinical relevance of these findings, we accessed H3K4me3 ChIP-Seq data of 19 glioblastoma patients (GSE121723).^38^ Analysing two patients’ datasets, we found 3,577 H3K4me3 peaks in patient AK124 and 3,565 peaks in patient AK231 to align with the PRDM9 motif centre (Fig. 5e, Supplementary Table 6-7), similar to our results with RKI1 glioblastoma cells. We leveraged parallel H3K36me3 ChIP-seq data, and overlapped intergenic H3K4me3 and H3K36me3 peaks with the PRDM9 motif (Fig. 5f) to confirm PRDM9 activity in glioblastoma, as the co-occurrence of H3K4me3, H3K36me3 and PRDM9 motif in intergenic regions is uniquely associated with PRDM9.^39^ Genome wide H3K4me3 signals for each patient (Supplementary Fig. 5a) were consistent with our RKI1 cell data (Fig. 4d), and we found significantly higher H3K4me3 intensities for the top 50 down-DEGs (Supplementary Fig. 5b) compared to the top 50 upregulated genes (Supplementary Fig. 5c-d). *DHCR7* and *DHCR24* genome tracks revealed strong H3K4me3 peaks at their promoters, aligning with the PRDM9 motif (Fig. 5g), as observed in RKI1 cells (Fig. 4i, 5d). Finally, the widths of H3K4me3 peaks at the *DHCR7* and *DHCR24* promoters in 19 patients showed a positive correlation with their corresponding mRNA levels (Fig. 5h), supporting our findings obtained with RKI1 glioblastoma cells.

### PRDM9 inhibitor MRK-740 depletes cholesterol in persisters

Our data thus far suggest that PRDM9 is a vulnerability in drug-tolerant persister cells in glioblastoma. Mechanistically, we established that PRDM9-dependent H3K4me3 regulates transcripts of enzymes which catalyse early (MVD) and late (EBP, DHCR7, DHCR24) steps in cholesterol biosynthesis (Fig. 6a). In further support, protein levels of these enzymes were reduced following PRDM9 inhibition with MRK-740 (Fig. 6b) or PRDM9 knockout (Fig. 6c), particularly in cells treated with CMPD1. To assess how PRDM9 inhibition impacts cholesterol levels in cells, we quantified intermediates lathosterol, zymosterol, desmosterol and cholesterol in RKI1 cells treated with CMPD1 combined with MRK-740 or the inactive MRK-740-NC. Single MRK-740 reduced lathosterol, zymosterol and desmosterol, leading to ∼60% reduction in cholesterol (Fig. 6d). CMPD1 nearly depleted desmosterol (Fig. 6d), the final cholesterol precursor in the Bloch pathway, indicating that persisters use the Kandutsch-Russell pathway. Indeed, lathosterol and cholesterol were reduced by 75%, but not depleted in CMPD1-treated cells. Importantly, cholesterol levels were further reduced in cells co-treated with CMPD1+MRK-740 (Fig. 5d). The compound MRK-740-NC, which does not inhibit PRDM9^30^, H3K4me3 (Fig. 3e) and failed to eliminate persister cells (Fig. 3c-d), did not affect cholesterol levels (Fig. 6d), further supporting the notion that cholesterol biosynthesis is crucial for glioblastoma persisters.

**Figure 6.**
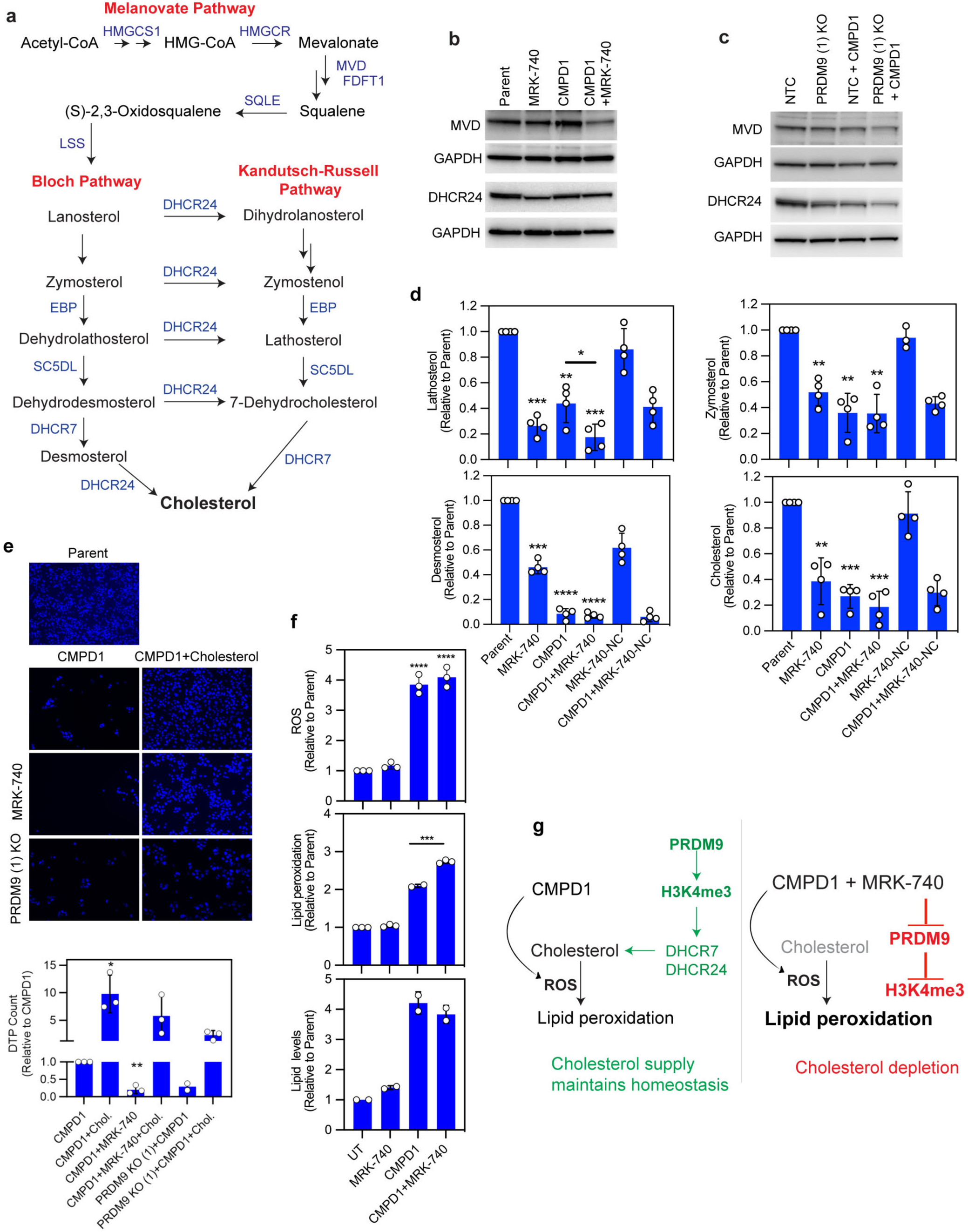
PRDM9 maintains cholesterol homeostasis under chemotherapy stress. a) Schematic of cholesterol biosynthesis pathways. b) Representative immunoblots of MVD and DHCR24 in RKI1 cells with CMPD1 (10 µM) ± MRK- 740 (3 µM) for 3 days. Each membrane was re-probed for GAPDH. c) Representative immunoblots of MVD and DHCR24 in in CMPD1 (10 µM, 3 days) treated RKI1 cells transduced with NTC sgRNA (NTC) or PRDM9 sgRNA (PRDM9 KO). Each membrane was re-probed for GAPDH. d) GC-MS quantification of cholesterol intermediates and cholesterol in RKI1 cells treated with CMPD1 (10 µM) ± MRK-740 (3 µM)/MRK-740-NC (3µM) for 3 days. Data are mean ± SD (n = 4; one sample t-test between sample vs parent; unpaired t-test between samples). e) DAPI-stained images and quantification of drug-tolerant persisters (DTP) in RKI1 cells treated with CMPD1 (25 µM) ± MRK-740 (3 µM) ± cholesterol complexed to methyl-beta-cyclodextrin (100 μg/mL, cholesterol component is 5% w/w) for 14 days. Bottom two images are of RKI1 cells transduced with PRDM9 sgRNA (PRDM9 KO). Data are mean ± SD (n = 3; one sample t-test). f) Flow cytometry quantification of reactive oxygen species (ROS), lipid peroxidation and total lipids in RKI1 cells treated with CMPD1 (10 μM) ± MRK-740 (3 μM) for 3 days. Data are mean ± SD (n = 3; one sample t-test). g) Working model of PRDM9-dependent cholesterol biosynthesis and persister survival.

If true, persisters should be sensitive to any agent targeting cholesterol. Statins inhibiting HMGCR in the mevalonate pathway would be the most obvious agents to test this hypothesis. However, the essentiality of mevalonate pathway^40^, coupled with the fact that statins have pleiotropic cholesterol-independent activity^41^, would not allow us to specifically validate cholesterol as critical to persister survival when using statins. We therefore targeted DHCR7 and DHCR24, as both are non-essential cancer genes (Supplementary Fig. 6a) and directly regulated by PRDM9 (Figure 4-5). The accumulation of the DHCR7 substrate zymosterol by the DHCR7 inhibitor AY-9944, along with the accumulation of the DHCR24 substrate desmosterol by the DHCR24 inhibitor SH-42, confirmed target engagement. (Supplementary Fig. 6b, 6c). As single agents, both AY-9944 and SH-42 reduced cholesterol levels to 20% compared to untreated cells (Supplementary Fig. 6c). However, this reduction had no effect on cell viability (Supplementary Fig. 6d), indicating that in the absence of chemotherapy, cells tolerate disruptions to cholesterol biosynthesis, as observed with MRK-740 treatments (Fig. 3c). Nevertheless, when combined with CMPD1, both DHCR7 and DHCR24 inhibitors significantly reduced persister-derived colonies, phenocopying CMPD1+MRK-740 treatment (Supplementary Fig. 6d).

Of note, glioblastoma stem cells are cultured in serum-free media without an external cholesterol supply and their survival depends on their ability to synthesise cholesterol. These culturing conditions accurately replicate the human brain environment, where the only source of cholesterol is *de novo* biosynthesis as dietary/peripheral cholesterol cannot penetrate the blood-brain barrier.^42^ To further confirm cholesterol biosynthesis as critical to the survival of drug-tolerant persister cells, we performed rescue experiments. Adding exogenous cholesterol to the cell culture media increased the number of CMPD1-tolerant persister cells by 10-fold (Fig. 6e), fully rescued MRK-740-induced cytostatic effect (Supplementary Fig. 6e) and persister cell death upon PRDM9 inhibition with MRK-740 or PRDM9 knockout (Fig. 6e).

Thus, our findings confirm PRDM9-H3K4me3-dependent cholesterol biosynthesis as a vulnerability in persister cells, yet they also raise an important question: why do persister cells rely on cholesterol biosynthesis, despite their non-proliferative state generally being linked to reduced cholesterol.^43^ In line, we observed a reduction in cholesterol precursors and final cholesterol levels in cells treated with CMPD1 for 3 days (Fig. 6d) and 14 days (Fig. 1f), even though the expression of cholesterol biosynthesis genes either remained unchanged or was upregulated in CMPD1-treated cells (Fig. 4). Given that chemotherapy, including microtubule-targeting agents induces oxidative stress which in turn oxidises lipids^9^, we questioned whether cholesterol could be sequestered through oxidation during CMPD1 treatment. Indeed, reactive-oxygen species (ROS) and lipid peroxidation dramatically increased in CMPD1-treated cells. While MRK-740 did not generate ROS or exacerbate the oxidative stress caused by CMPD1, it increased the proportion of peroxidised lipids in CMPD1- treated cells, without changing the total lipid levels (Fig. 6f). All in all, these data suggest CMPD1- triggered oxidative stress leads to cholesterol oxidation. To maintain cholesterol homeostasis and survive, persister cells depend onPRDM9-H3K4me3 regulated *de novo* cholesterol biosynthesis. However, when PRDM9 is inhibited, cholesterol supply is disrupted. Combined effects of CMPD1 and MRK-740 have a more pronounced impact on cholesterol homeostasis compared to each agent alone, leading to persister cell death (Fig. 6g).

### Brain permeable microtubule-targeting agent in combination with cholesterol depletion increases survival in glioblastoma xenografts

The mechanistic studies were completed with CMPD1, which is an MTA with physicochemical properties compatible with blood-brain barrier permeability.^21^ However, CMPD1 exhibited extremely short (t_1/2_ = 4.6 min) half-life in the stability assay using human liver microsomes, preventing its use *in vivo* (Fig. 7a). To improve its metabolic stability, we performed medicinal chemistry optimisation of CMPD1 (Supplementary Table 8) and developed analogue WJA88 with improved cellular efficacy (EC_50_ = 150 nM, A172 cell viability) and metabolic stability (t_1/2_ = 34.2 min) compared to CMPD1 (Fig. 7a-b). We confirmed that both CMPD1 and WJA88 inhibit tubulin polymerisation and bind into the colchicine binding site (Supplementary Fig. 7a-b). Pharmacokinetic analysis of mice administered a single 50 mg/kg WJA88 dose intraperitoneally revealed that WJA88 had an *in vivo* half-life (t_1/2_) of 71.4 min and peak serum concentration (c_max_) of 7,907 ng/mL (Supplementary Table 9). Distribution across the blood-brain barrier was evaluated in mice dosed with 50 mg/kg WJA88 once intravenously, which showed a brain/plasma ratio of 0.8 and 0.76 at 30 min and 60 min, respectively (Fig. 7c). Therefore, we reasoned that WJA88 with improved stability and sufficient brain uptake could be tested in intracranial models. In functional assays, WJA88 inhibited proliferation of glioblastoma stem cell lines with nanomolar potency (Supplementary Table 10) and attenuated the growth of MMK1 spheroids (Fig. 7d). Similarly to CMPD1, WJA88, even at a high concentration of 25 μM, led to the emergence of persister cells, and MRK-740 significantly reduced the number of WJA88-tolerant persisters (Fig. 7e). However, MRK-740 lacks pharmacokinetics suitable for *in vivo* studies.^30^ As an alternative, we used the Liver X Receptor (LXR) agonist LXR-623 due to its high brain uptake in rodents.^44^ Single LXR-623 (400 mg/kg) suppressed glioblastoma growth by enhancing ABCA1-mediated cholesterol efflux.^44^ We confirmed that LXR-623 increased ABCA1 expression (Supplementary Fig. 7c) and reduced the number of WJA88-tolerant persisters in both RKI1 and FPW1 cells (Fig. 7e). Further, the anti-persister efficacy of LXR-623 was fully rescued by exogenous cholesterol (Supplementary Fig. 7d). Thus, LXR-623 showed a persister-killing efficacy comparable to MRK-740, albeit through different mechanisms: MRK-740 inhibits cholesterol biosynthesis, whereas LXR-623 promotes cholesterol efflux.

**Figure 7.**
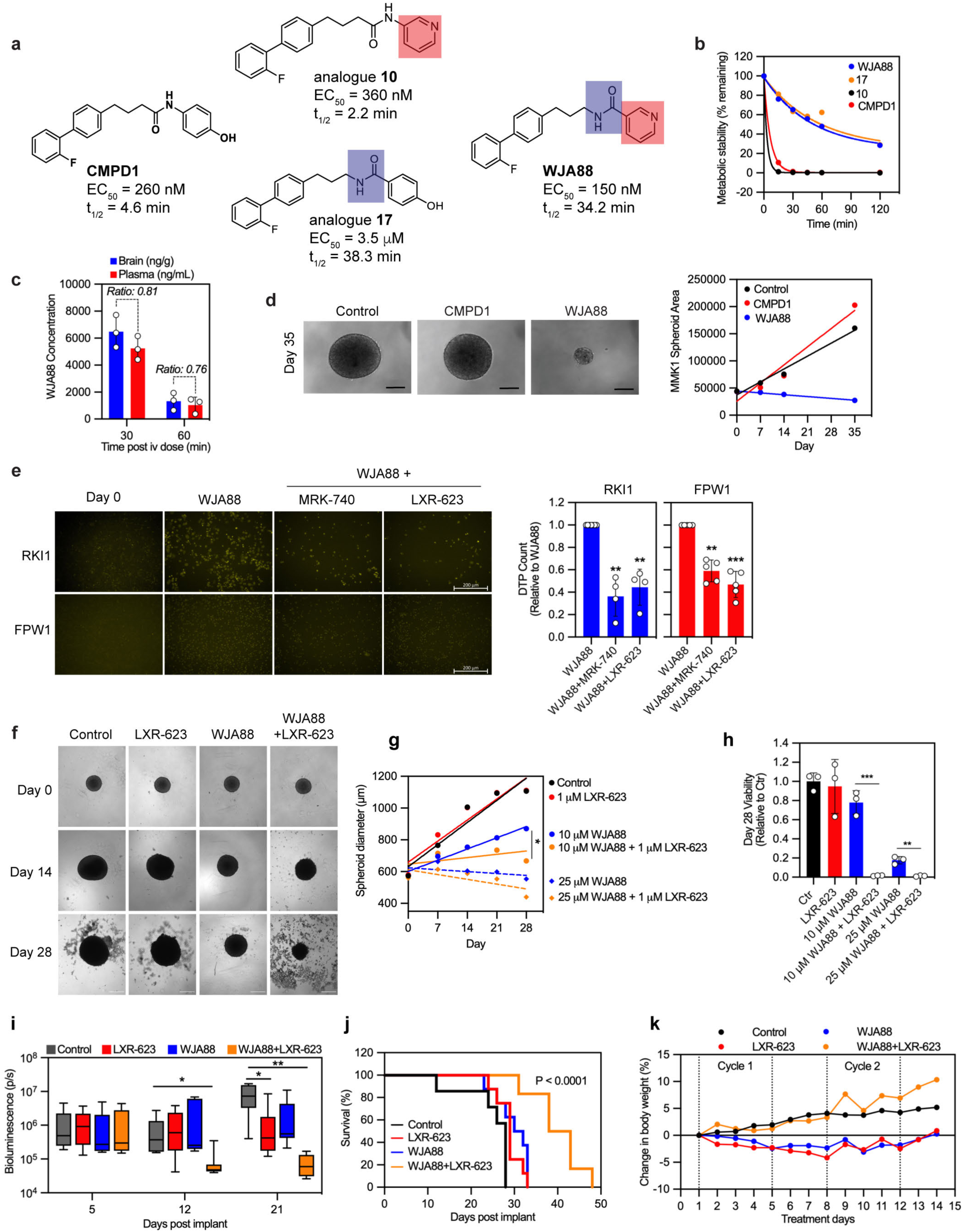
Efficacy of brain-permeable microtubule-targeting agent and LXR agonist in glioblastoma xenografts. a - b) Chemical structures, cellular efficacy (EC_50_; A172 cell viability) and metabolic stability (t_1/2_; human liver microsomes) of CMPD1, analogues **10** and **17**, and WJA88. c) Plasma and brain concentrations of WJA88 in male CD-1 mice (n = 3) of a single intravenous administration at 50 mg/kg. d) Images and quantification of MMK1 spheroids growth when treated with CMPD1 (5 μM) and WJA88 (5 μM). Data are mean of 2-3 spheroids per treatment group. Scale bar 100 nm. e) Nuclear-ID red stain images (pseudo-coloured yellow) and quantification of RKI1 and FPW1 drug- tolerant persister (DTP) cells surviving WJA88 (25 µM, 14 days) treatment combined with MRK- 740 (3 µM) or LXR-623 (1 µM). Data are mean ± SD (n = 3; one sample t-test). f - h) Images and quantification of GBM6 spheroids growth and viability when treated with WJA88 (10 or 25 μM) and LXR-623 (1 μM). Data are mean of 3 spheroids per treatment group (unpaired t- test). Scale bar 100 nm. i) Bioluminescence signal quantification of GBM6 orthotopic tumours in mice treated with WJA88 (50 mg/kg) ± LXR-623 (100 mg/kg). Boxplots show mean (middle line) ± SD, whiskers represent minimum to maximum data points (n = 8 per treatment arm; unpaired t-test). j) Kaplan-Meier regression showing survival (%) in mice with GBM6 orthotopic tumours and treated with WJA88 (50 mg/kg) ± LXR-623 (100 mg/kg). Significance was determined with log rank (Mantel-Cox) test between the survival curves for control (vehicle) vs tested agents (n = 8 per treatment arm). k) Changes in body weight of mice with GBM6 orthotopic tumours treated as in (H).

To test WJA88 and LXR-623 combination *in vivo*, we used GBM6 cells carrying EGFRvIII mutation and forming aggressive fast-growing tumours in animals. We first confirmed the efficacy of the co- treatment in GBM6 spheroids (Fig. 7f-h). Next, GBM6 cells were surgically engrafted into the cortex of Balb/c/nude mice, and mice were randomized to treatment cohorts on Day 5. As monotherapy, WJA88 (50 mg/kg) demonstrated short but significant extension in survival relative to vehicle control (median 31 days vs. 28 days: log-rank P < 0.039). LXR-623 (100 mg/kg) monotherapy had negligible effect on survival (median 29 days). Importantly, the combination of WJA88 and LXR-623 effectively inhibited tumour growth (Fig. 7i). Reduced tumour growth translated into prolonged survival of orthotopic tumour-bearing mice treated with the combination therapy compared with monotherapies or vehicle control (median 40.5 days vs. 28 days: log-rank P < 0.0001; Fig. 7j). The WJA88 and LXR-623 combination was well tolerated with no significant change in body weight after the treatments were started (Fig. 7k). In contrast, the minor reduction (up to 5%) in body weight observed with both WJA88 and LXR-623 when administered as standalone treatments was reversed when these two agents were combined. Furthermore, only mild effects on blood parameters were observed in mice receiving WJA88 and LXR-623 (Supplementary Fig. 7e). Taken together, these findings imply that the reliance on cholesterol following MTA chemotherapy was more prominent in glioblastoma cells compared with cells of other vital organs.

## DISCUSSION

Efforts to target drug-tolerant persister cells in cancer have primarily focused on enhancing the efficacy of clinical drugs. However, effective treatments for glioblastoma are lacking, which limits drug tolerance research in neuro-oncology to experimental therapies. Here, using brain-permeable microtubule-targeting agents we show that glioblastoma persister cells rely on the histone methyltransferase PRDM9 for survival. Conversely, PRDM9 inhibition increased MTA chemosensitivity, enhancing both the magnitude and duration of the antitumor response by eliminating persister cells.

PRDM9 had been exclusively studied in DNA recombination during meiosis.^17^ We found that PRDM9 is a lowly expressed non-essential enzyme in glioblastoma cells, and its inhibition leads to reversible cytostatic effects. However, chemotherapy-induced mitotic arrest leads to PRDM9 upregulation and under the oxidative stress of chemotherapy, PRDM9 promotes cholesterol homeostasis and persister cell survival. This mechanism appears to be relevant across various glioblastoma cell lines and may apply to other cancers as well. The observed increase in PRDM9 levels in mitotically arrested cells suggests that tumours treated with MTAs or drugs inducing mitotic arrest, such as Polo-like or Aurora kinase inhibitors, could upregulate PRDM9, enabling the emergence of persister cells and tumour recurrence. Further, our findings on the molecular and cellular functions of PRDM9 in glioblastoma expand the understanding of cancer-testis genes role in cancer. Notably, other cancer-testis genes, such as BORIS and MAGE-A3/6, have been also linked to drug resistance and metabolic rewiring in tumours.^45, 46^

A trait often observed in persister cells is the demethylation of H3K4me3 due to increased KDM5 activity.^47, 48^ KDM5 inhibitors, by increasing H3K4me3 abundance, reduced the number of drug-tolerant persisters in multiple cancer types.^2, 49^ Like previous studies, we observed decreased H3K4me3 after 14 days of chemotherapy. However, KDM5 inhibitors failed to eradicate glioblastoma persisters and our data suggest that the loss of H3K4me3 stems from the proteolytic cleavage of histone tails, rather than the activities of KDM5 demethylases. Histone cleavage, a well- established mechanism for epigenetic regulation, has been associated with induction of senescence^27^ which is also a hallmark of drug-tolerant persisters. Importantly, we show that PRDM9-dependent H3K4 methylation, occurring before histone cleavage, is a transient yet critical event for establishing the drug-tolerant state. Our findings align with a study revealing that the persisters survival in breast cancer is primed through the bivalent chromatin, where genes prepared for activation by H3K4me3 are concurrently repressed by H3K27me3. Maintaining H3K27me3 with a KDM6 inhibitor, thereby blocking H3K4me3 activity, eliminated persisters.^50^ Thus, in a cancer-dependent context, both reduced and elevated H3K4me3 abundance can provide a survival advantage to drug-tolerant persisters.

H3K4me3 marks transcription start sites and is widely believed to regulate transcriptional initiation, with recent study revealing a role for H3K4me3 in RNA polymerase pause-release and elongation.^51^ Either way, H3K4me3 creates a permissive environment for gene activation during cell development, differentiation, and response to environmental cues. Additionally, H3K4me3 is recognized for its role in connecting metabolic reprogramming to epigenetic alterations. Changes in methionine availability, the precursor of S-adenosylmethionine which acts as a co-factor for lysine methyltransferases, impact H3K4me3 peak width and, consequently, gene expression.^52^ In turn, H3K4me3 marks metabolic genes, contributes to a transgenerational epigenetic inheritance of obesity and regulates purine and pyrimidine nucleotide synthesis pathways.^53, 54, 55^ We extend these studies by linking PRDM9-dependent H3K4me3 to the regulation of cholesterol biosynthesis genes, further supporting the H3K4me3-dependent connection between epigenetic and metabolic plasticity.

Cholesterol has been long recognised as a key factor in cancer treatment failure.^44, 56^ Yet, cancer clinical trials with cholesterol-lowering drugs (mostly statins) have not shown consistent or significant benefits for cancer patients. While the reasons for this are diverse, one of the issues is that statins directly (by targeting ubiquitously expressed HMGCR) and indirectly (by lowering circulating cholesterol) reduce cholesterol levels in multiple cell types, adversely affecting membrane microdomains, steroidogenesis and cell viability.^57^ The cholesterol-lowering effect of PRDM9 inhibition, combined with its tumour-restricted expression, encourages the development of PRDM9- targeting drugs that could selectively target cholesterol in tumours. To generate a proof-of-concept in glioblastoma models, we developed a brain-permeable microtubule-targeting agent WJA88 and show that a combination of WJA88 with cholesterol depletion (using the brain-permeable LXR agonist LXR-623) significantly reduced tumour growth and prolongs survival in glioblastoma-bearing mice. In summary, our findings reveal a previously unrecognised role of the testis-specific histone methyltransferase PRDM9 in cancer. We uncover a molecular mechanism through which PRDM9 promotes survival of drug-tolerant persister cells, with important implications for enhancing the clinical effectiveness of chemotherapy with microtubule-targeting agents.

## METHODS

### Cell lines

Glioblastoma stem cell lines RKI1, FPW1, HW1, WK1, MMK1 and SB2b were derived from glioblastoma patient specimens. Characterisation of these cell lines including RNA sequencing, mutational profiling, subtype assignment and proteomic data are available online at https://www.qimrberghofer.edu.au/our-research/commercialisation/q-cell/ and in Ref.^31^ GBM6 glioblastoma cells (EGFR/EGFRvIII over-expression) were obtained from Paul Mischel, Ludwig Institute of Cancer Research, USA. Cells were cultured in KnockOut DMEM/F-12 basal medium kit with neural supplement, EGF (20 ng/mL) and FGF-β (10 ng/mL) (ThermoFisher Scientific, Cat# A1050901). GlutaMAX-ICTS (2 mM) (Thermofisher Scientific, Cat# A1286001) and Antibiotic- Antimycotic (Thermofisher Scientific, Cat# 15240112) were also added. Adherent cells were plated on flasks coated with 0.15% in PBS MatriGel Matrix (Corning Life Sciences, Cat# BDAA356237), incubated at 37 °C, 5% CO_2_. A172 glioblastoma cell line (Cat# 88062428) was purchased from the European Collection of Authenticated Cell Cultures (ECACC, Salisbury, UK) through Cell Bank Australia and cultured as previously described.^8^ All cell cultures were routinely tested for mycoplasma infection and the cumulative length of culturing did not exceed 10 passages.

### Chemical probes and inhibitors

CMPD1 (CAS# 41179-33-3, Cat# sc-203138) was purchased from Santa Cruz. Colchicine (CAS# 64-86-8, Cat# 1364), paclitaxel (CAS# 33069-62-4, Cat# 1097) and vinblastine (CAS# 143-67-9, Cat# 1256) were purchased from Tocris. SGC Probe Set (Supplementary Figure 3a; Cat#17748), CPI- 169 (CAS# 1450655-76-1, Cat# 18299), EPZ6438 (tazemetostat, CAS# 1403254-99-8, Cat# 16174), GSK126 (CAS# 1346574-57-9, Cat# 15415), AY-9944 (CAS# 366-93-8, Cat# 14611), SH-42 (CAS# 2143952-36-5, Cat# 34677) and Ro-3306 (CAS# 872573-93-8, Cat# 15149) were purchased from Cayman Chemicals. MRK-740 (CAS# 2387510-80-5, Cat# HY-114209) was purchased from MedChem Express. Tivantinib (CAS# 905854-02-6, Cat# S2753) was purchased from Selleckchem. CPI-1205 (CAS# 1621862-70-1, Cat# A16357) was purchased from AdooQ Bioscience. MRK-740- NC (CAS# 2421146-31-6, Cat# SML2536) was purchased from Sigma Aldrich. LXR-623 (CAS# 875787-07-8, Cat# 21117) was purchased from Sapphire Biosciences.

### Animal models

BALB/c Nude mice (Animal Resource Centre, Perth, Western Australia) were injected with 3x10^5^ GBM6 cells in the cortex, and tumour imaging was done on Day 5. On Day 6, animals were randomly grouped into the following treatment groups: *i)* Vehicle, *ii)* single WJA88, *iii)* single LXR-623 and *iv)* WJA88 + LXR-623. WJA88 was formulated using 40% Captisol and administered intraperitoneally. LXR-623 was formulated using 10% DMSO,15% PEG 300, 5% Tween 80, and 70% Milli Q water and administrated orally. Starting on Day 6 post-implantation, WJA88 and LXR- 623 were administered for two weeks (5 days on, 2 days off), and no significant weight loss was observed. WJA88 was dosed at 50 mg/kg, and LXR-623 was dosed at 100 mg/kg BID (i.e., 2 x 50mg/kg). Animals received ad-libitium food and water, with nutritional enrichment when required. Tumour growth was monitored using the bioluminescence imaging once a week during and after the treatment to evaluate the efficacy of the treatments. Toxicity was monitored twice weekly from all treatment groups, blood counts (n = 3 per group) from whole blood collected in tubes coated with heparin, and sample analysis was performed using BC 5000 hematology analyzer from Mindray. Post-treatment, 3 animals in the combination treatment group were culled for non-tumour related issues, and during necropsy, we did observe cardiomegaly or bilateral renal enlargement compared to control mice. All the studies were performed under the Telethon Kids Institute Animal Ethics 362 rules and regulations.

### Persister generation and expansion

FPW1 and RKI1 cells (1.5 × 10^4^ cells/cm^2^) were treated with CMPD1 (25 μM) or tivantinib (25 μM) for 14 days. Every 3 days, the media containing CMPD1 or tivantinib was replaced. At Day 14, drug-tolerant persister (DTP) cells were collected for further analyses. Alternatively, DTP cells were allowed to recover in drug-free media (drug holiday) until they regained their morphology and expansion started when cells resembled their parental counterparts. Images were taken using Zeiss Axio Vert.A1 microscope using the ZEN 2 – blue edition software (Zeiss).

### Persisters quantification

Cells (1.5 × 10^5^ cells/well) were seeded on black imaging 24-well plates (Eppendorf) and treated with tested compounds for 14 days. To remove dead cells, media was replaced with fresh media containing tested compounds every 3 days. Untreated (Day 0) and treated (Day 14) cells were stained with Nuclear-ID red stain (Enzo Lifesciences) at 1:1,000 dilution in StemPro media. Cells were incubated with the stain for 30 min prior to washing with PBS. Alternatively, when DAPI stain was used, on Day 14 cells were fixed in 1% PFA/PBS for 10 min, washed three times with PBS, stained and permeabilised in 1 µg/mL DAPI and 0.1% Triton X-100 in PBS for 15 min. Cells were imaged using Zeiss Axio Scope.A1 and ZEN 2 – blue edition software (Zeiss) and Fiji was used for quantification (nine images per well).

### Cell viability and GR metrics

Treatment-naïve glioblastoma cells (2 × 10^3^ cells/well) or drug-tolerant persister cells (8 × 10^3^ cells/well) were treated with DMSO or tested compounds at 8-point dilution row for 5 days. CellTitre-Blue (Promega, Cat# G808B) was added (1:10) and incubated at 37 °C for 2-4 h. Fluorescence was measured with a Tecan M200 PRO+ microplate reader (Tecan) at Ex/Em 530/590. Data were normalised to DMSO-treated controls (set as 1). Per-division GR_50_, GR_max_, h_GR_, and GR_AOC_ metrics were calculated from cell viability data on Day 0 and Day 5 using the *GRcalculator* online tool.^22^ Graphs were recreated using GR values from the *GRcalculator* and Prism v8.0 (GraphPad).

### Colony formation assay

Cells (1 × 10^4^ cells/well) were treated with the tested compounds for 7 days, then washed with PBS and allowed to recover in drug-free media for an additional 7 - 21 days (depending on the cell line). Once visible colonies were formed in CMPD1-only or WJA88-only treatments, colonies were stained with 1% toluidine blue (> 4 hr, 4 °C). Plates were imaged using Bio-Rad Gel Doc and colony area quantified using the ImageJ software.

### Spheroid growth assay

MMK1 and GBM6 cells (10 × 10^3^ cells/well) were seeded into 96-well round bottom low attachment plates and centrifuged at 190 g for 3 min. Cells were left undisturbed for 48 h to form spheroids, then treated with tested compounds or DMSO (vehicle) for 28 days, with drug-containing media replenished every 4 days. Spheroids were imaged on an AxioVert A1 microscope immediately prior to treatment and every 7 days following the initiation of treatment to determine spheroid diameter using ImageJ. After 28 days of treatment, a PrestoBlue (Thermofisher Scientific, Cat # A13261) assay was performed according to the manufacturer’s instructions with an incubation period of 3 h. Fluorescence was measured with a FLUOstar Omega microplate reader (BMG Labtech), and readings were normalised to the average of DMSO-treated vehicle controls. Graphing and statistical analyses were performed with GraphPad Prism 9.

### Incucyte Live Cell Imaging

Cells (4×10^4^ cells/well) were treated as described and placed into the Incucyte SX5 at 37 °C and 5% CO_2_. 16 images were taken per well every 2 hours. Confluency was calculated using the Incucyte SX5 complementary software.

### RNA sequencing

RNA extraction was performed with RNeasy Minikit (Qiagen, Cat# 74104), according to manufacturer’s instructions, with additional on column DNA digestion using RNase free DNase set (Qiagen, Cat# 79254). For the sequencing of DTPs (Figure 1), sequencing libraries were generated using NEBNext® UltraTM RNA Library Prep Kit for Illumina (New England Biolabs, Cat# E770) following manufacturer’s instructions. For sequencing of RKI1 cells treated with CMPD1 ± MRK- 740 or RKI1 cells transduced with NTC/PRDM9 sgRNA (Figure 4), RNA extracted from samples underwent library preparation using mRNA stranded library preparation kit (Illumina, Cat# 20040532) with associated indexes and anchors, following manufacturer’s instructions. Libraries were quantified using NEBnext Ultra II DNA library preparation kit for Illumina (New England Biolabs, Cat# E7630L). For sequencing in Figure 1, clustering of the index-coded samples was performed on a cBot Cluster Generation System using PE Cluster Kit cBot-HS (Illumina, discontinued) according to the manufacturer’s instructions. After cluster generation, the library preparations were sequenced on an Illumina Novaseq. For Figure 4, the libraries were sequenced on Illumina Nextseq2000 system. FASTQ generation, alignments and counts were processed on instrument using Dragen RNA pipeline. Heatmaps were generated using Pheatmap package, gene sets were mapped to gene ontology terms using clusterProfiler package and volcano plots were generated using ggplot2 package, all in RStudio. For data in Figure 4c, a batch correction was performed using RUV and a set of stably expressed genes.^58^ Samples were processed in three biological replicates with significance determined through adjusted P-value.

### LC-MS/MS analysis of histone peptides

All solvents, acids and bases used were HPLC-grade (Sigma-Aldrich) and all reactions were carried out in Protein-LoBind Eppendorf tubes. Chromatin-bound fractions from snap-frozen cell pellets were extracted using the Sub-Cellular Fractionation Kit (ThermoFisher Scientific, Cat # 78840) as per manufacturer’s instructions. The samples were supplemented with 50 mM sodium bicarbonate, Protease Inhibitor Cocktail, 1 mM PMSF, 1 mM sodium orthovanadate and 10 mM sodium butyrate (All Sigma-Aldrich). Protein concentration was measured using Pierce BCA Protein Assay (ThermoFisher Scientific, Cat # 23225). Proteins were precipitated with methanol:chloroform, air-dried, reconstituted to ∼ 2.5 mg/mL in 50 mM sodium bicarbonate (pH 8.0) and sonicated (SonoPlus Mini20 (Bandelin)). Histones were propionylated and trypsinised for bottom-up analysis as described by Sidoli et al. Samples were then acidified with 1% trifluoroacetic acid and desalted using Oasis HLB Sep-Pak columns (Waters) and vacuum dried.

For LC/MS-MS, 1 μg of propionylated histones in loading buffer (3% acetonitrile, 0.1% TFA) was injected onto a 30 cm × 75 μm inner diameter column packed in-house with 1.9 μm C18AQ particles (Dr Maisch GmbH, HPLC) using a Dionex Ultimate 3000 nanoflow UHPLC. Peptides were separated using a linear gradient of 5–35% buffer B over 120 min at 300 nL/min at 55 °C (buffer A: 0.1% (v/v) formic acid; buffer B: 80% (v/v) acetonitrile and 0.1% (v/v) formic acid). All MS analyses were performed using a Q-Exactive HFX mass spectrometer. For Data-Dependent Acquisition (DDA): after each full-scan MS1 (R = 120,000 at 200 *m/z*, 300–1600 *m/z*; 3 × 10^6^ AGC; 110 ms max injection time), up to 10 most abundant precursor ions were selected for MS/MS (R = 45,000 at 200 *m/z*; 2 × 10^5^ AGC; 86 ms max injection time; 30 normalised collision energy; peptide match preferred; exclude isotopes; 1.3 *m/z* isolation window; minimum charge state of +2; dynamic exclusion of 15 s). For DIA: after each full-scan MS1 (R = 60,000 at 200 *m/z* (300–1600 *m/z*; 3 × 10^6^ AGC; 100 ms max injection time), 54 × 10 *m/z* isolations windows (loop count = 27) in the 390–930 *m/z* range were sequentially isolated and subjected to MS/MS (R = 15,000 at 200 *m/z*, 5 × 10^5^ AGC; 22 ms max injection time; 30 normalised collision energy). 10 m/z isolation window placements were optimised in Skyline^59^ to result in an inclusion list starting at 395.4296 *m/z* with increments of 10.00455 *m/z*. This resulted in a duty cycle of ∼ 2.2 s.

Database searches were performed using Mascot v2.4 against the human SwissProt database (May 2019; 559,634 entries) using a precursor-ion and product-ion mass tolerance of ± 10 ppm and ± 0.02 Da, respectively. The enzyme was specified as ArgC with a maximum of 1 missed cleavage. Variable modifications were set as: acetyl(K), propionyl(K), monomethyl + propionyl(K), dimethyl(K) and trimethyl(K), propionyl(N-term), oxidation(M), carbamidomethyl(C). All DIA data were processed using Skyline (v20.1).^59^ Reference spectral libraries were built in Skyline using Mascot .dat files using the BiblioSpec algorithm.^60^ A False Discovery Rate (FDR) of 5% was set and a reverse decoy database was generated using Skyline.^61^

### Data normalisation and statistical analysis

Processed MS1 quantifications were first Log transformed (base 2) and then quantile-normalised across samples. Data from FPW1 and RKI1 were subsequently analysed independently. For data from each cell type, Combat R package^62^ was used to remove experimental batch effects.^62^ Batch corrected data were used for downstream analyses including hierarchical clustering, heatmap visualisation and bar graphs.

### ChIP sequencing

RKI1 cells treated with CMPD1 ± MRK-740 or RKI1 cells transduced with NTC/PRDM9 sgRNA ± CMPD1 were harvested, cell pellets were cross-linked with 0.75% PFA/PBS for 10 min, quenched in 1 M glycine, washed in PBS, and snap frozen in liquid nitrogen. Cells were lysed in ChIP lysis buffer (50 mM HEPES-KOH pH 7.5, 140 mM NaCl, 1 mM EDTA pH 8, 1% Triton X-100, 0.1% sodium deoxycholate, 0.1% SDS, Protease Inhibitor Cocktail) and sonicated to shear chromatin to ∼300bp DNA fragment size. DNA fragment size was checked via 1.5% agarose gel electrophoresis against a 100-1000bp DNA. Input samples were aliquoted for later DNA extraction processing. Chromatin immunoprecipitation was carried out with H3K4me3 antibody (Cell Signalling Technology, Cat # 9751) added 1:50 to chromatin lysates. The samples were agitated overnight at 4°C with bead capture of antibody bound chromatin fragments using Dynabeads Protein A (Thermofisher Scientific, Cat# 10002D). The samples were then washed in the following buffers: 1) low salt wash buffer (0.1% SDS, 1% Triton X-100, 2 mM EDTA, 20 mM Tris-HCl pH 8.0, 150 mM NaCl); 2) high salt wash buffer (0.1% SDS, 1% Triton X-100, 2 mM EDTA, 20 mM Tris-HCl pH 8.0, 500 mM NaCl); 3) LiCl wash buffer (0.25 M LiCl, 1% NP-40, 1% sodium deoxycholate, 1 mM EDTA, 10 mM Tris-HCl pH 8.0). DNA protein complexes were eluted from antibody captured beads in 1% SDS, 100 mM NaHCO_3_. Chromatin fragments were subject to overnight de-cross linking in 0.32 M NaCl and RNA digestion with 10 µg RNAse A at 65°C. Protein was digested using 20 µg Proteinase K for 1 hour at 60°C. DNA was extracted using ChIP Clean and Concentrate Kit (Zymo Research, Cat# D5205). Input and ChIP fragments were quantified using QUBIT dsDNA kit (Thermofisher Scientific, Cat# Q33230) and 5 ng DNA fragments was subjected to library preparation using NEBNext Ultra II DNA library preparation kit for Illumina (New England Biolabs, Cat# E7645S). Libraries were quantified, sequenced on Nextseq2000 instrument. Dragen BCL Convert was used to generate FASTQ sequences for further downstream analysis using Deeptools 2.0 pipeline. Briefly, FASTQ files were aligned to hg38 homo sapiens reference genome, then bigWig density files were generated from the BAM alignments for subsequent downstream analyses. pyGenomeTracks was used for generating individual gene tracks of H3K4me3 enrichments. To identify PRDM9 motif occurrences across the genome, we obtained the reference genome sequence (hg38 for RKI1, hg19 for patient data GSE121723) in FASTA format, and homo sapiens PRDM9 motif matrix MA1723.2 was scored against it using FIMO.^63^ MACS2 callpeak was used to calculate H3K4me3 ChIPseq peak widths. Samples were carried out in biological duplicates.

### LC-MS/MS analysis of lipids

RKI1 parent and CMPD1-derived drug-tolerant persister cells (Figure 1) were collected, homogenized in sample extraction buffer (50 mM Hepes pH 7.4, 25 mM KCl, Protease Inhibitor Cocktail) by sonicating for 5 min (30 s on/30 s off) at 4°C with a Qsonica Q800R2 sonicating bath. Protein concentration was determined with the BCA assay. Lipids were extracted from 200 µL lysate (∼200 µg protein) using the methyl-tert-butyl ether (MTBE)/methanol/water protocol as previously described^64^. Cell homogenate was combined with 250 µL methanol containing 0.01% 3,5-di-tert-4- butylhydroxyltoluene (BHT), internal standards and 850 µL MTBE, sonicated in a 4°C water bath for 30 min, and phase separation was induced by centrifuging at 2000g for 5 min. The upper organic phase was transferred to a 5 mL glass tube, and the aqueous phase was re-extracted with 500 µL MTBE and 150 µL methanol. The organic phase from the second extraction was combined with the first, after which the extracts were dried in a Savant SC210 SpeedVac dessicator (ThermoFisher Scientific). Lipids were reconstituted in 400 µL of 80% (v/v) methanol:20% water containing 0.01% BHT, 1 mM ammonium formate, and 0.1% formic acid, and stored at -80 °C. Lipids were detected in multiple reaction monitoring mode on a TSQ Altis triple quadrupole mass spectrometer with a Vanquish HPLC (ThermoFisher Scientific) and Waters Acquity UPLC CSH 2.1x100 mm C18 column (1.7 µm particle size), as previously described.^65^ Run time was 25 min with a flow rate of 0.28 ml/min, using mobile phases A (10 mM ammonium formate, 0.1% formic acid, 60% acetonitrile and 40% water) and B (10 mM ammonium formate, 0.1% formic acid, 10% acetonitrile and 90% isopropanol) with the following binary gradient: 0-3 min, 20% B; 3-5.5 min, ramp to 45% B; 5.5-8 min, ramp to 65% B; 8-13 min, ramp to 85% B; 13-14 min, ramp to 100% B; 14-20 min, hold at 100% B; 20-25 min, decrease to 20% B and hold to 25 min. Acylcarnitine (AcCa), ceramide, sphingomyelin (SM), phosphatidylcholine (PC), lysophosphatidylcholine (LPC), and lysophosphatidylethanolamine (LPE) species were identified as the [M+H]^+^ precursor ion, with product ion m/z values of 85.0 for AcCa, 184.1 for PC, LPC and SM, 264.3 for ceramide and SM, and neutral loss of 141.0 for LPE. Diacylglycerol (DG) was detected as the [M+NH_4_]^+^ precursor ion, with product ions corresponding to neutral loss of 35.0 and RCOOH + NH_3_. Cholesterol was detected as precursor m/z 369.4 and product ion m/z 161.1. Phosphatidylethanolamine (PE), phosphatidylinositol (PI), and phosphatidylserine (PS) species were detected as the [M-H]^-^ precursor ion, with product ions corresponding to the fatty acyl anion. TraceFinder 4.1 (ThermoFisher) was used to integrate the peaks. The molar amount of each lipid was calculated with reference to its class-specific internal standard, then normalized to protein content.

### Gas Chromatography Mass Spectrometry analysis of cholesterol

RKI1 cells treated with CMPD ± MRK-740 ± MRK-740-NC (Figure 6) were collected, cell pellets washed in PBS, lysed with 10 M NaOH containing protease inhibitors and proteins quantified by BCA assay following manufacturers’ instruction. Heavy isotope standards (100 ng per sample) and methanol supplemented with 0.02% BHT (8:1 v/v) were added to each sample and samples were heated at 60 °C for 1 hour in 1 M KOH. The reaction was neutralised with 300 mM HCl, and sterols extracted in hexane. The samples were dried, derivatised (60°C, 1 hr) in acetonitrile:BSTFA (1:3 v/v) and toluene (4:1 v/v) was added. Derivatives were analysed by gas chromatography (Agilent 8890) and mass spectrometry (Agilent 5977 System). Briefly, derivatized samples (1 μl) were injected into a fused silica capillary column (60 m x 0.25-mm internal diameter) coated with cross-linked 50% phenylmethylsiloxane (film thickness 0.25 um; Restek Rxi-5ms) using a splitless injection for sterol intermediate analysis or a 30:1 split ratio for cholesterol analysis. The helium carrier gas flow rate was 1 mL/min. Selected ion monitoring was performed by using the electron-ionization (EI) mode at 70 eV. Quantification was performed by monitoring specific ions at standard retention times and compared using the peak area ratio of the sterol vs the internal standard, on the Agilent Masshunter software. Data was then normalised to protein quantifications. The following standards were used: Cholesterol-d7 (Cat# 791645), Zymosterol-d5 (Cat# 700072), Zymostenol-d7 (Cat# 700117), Desmosterol-d6 (Cat# 700040P) (all Avanti Polar Lipids), Lathosterol-d4 (Cat# D-5546), 7-Alpha-Hydroxycholesterol-d7 (Cat# D-4064), 7-Beta-hydroxycholesterol-d7 (Cat# D-4123) (all CDN isotopes), 24-hydroxycholesterol-d7 (Cat#, D730), 27-hydroxycholesterol-d5 (Cat# D733) (all Medical Isotopes), Squalene-d6 (Cat# TRC-S683802), Lanosterol-d6 (Cat# TR-L174584), 7- dehydrocholesterol-d7 (Cat# TRC-D229457), 25-hydroxycholesterol-d6 (Cat# TRC-H918032) (all TRC).

### Western blotting

For histone immunoblots in Figure 2, histones were extracted using the Histone Extraction Kit (Abcam, Cat # ab113476) as per manufacturer’s instructions, and protein concentrations were determined. 1 μg of histone extracts were resolved (30 min, 200 V) on 12% Bolt Bis-Tris gels (Cat# NW00127BOX) and transferred onto nitrocellulose membranes (Cat# IB301002) using iBlot 2 protein transfer system (Cat# IB21001) at 15 V, 7 min (all Thermofisher Scientific). For whole cell lysates, cell pellets were lysed in RIPA buffer containing protease inhibitor PMSF, protease inhibitor cocktail and phosphatase inhibitors (sodium orthovanadate), sonicated to remove viscosity, then protein concentration was measured. Samples were prepared with Bolt LDS loading buffer (Cat# B0007) and Bolt sample reducing agent (Cat# B0009) (both Thermofisher Scientific). Samples were heated at 95 °C for 5 min. 20-30 μg of total protein were resolved (2 h, 95 V) on 4-12% Bolt Bis-Tris gels (Cat# NW04120BOX) and dry transferred onto PVDF membranes (Cat# IB24001) at 20V, 7 min (all Thermofisher Scientific). Membranes were blocked with 5% skim milk powder in TBST, washed in TBST, and incubated with primary antibody in 5% BSA in TBST overnight at 4 °C. Membranes were washed in TBST and incubated with secondary antibody for 1 h at room temperature. Detection was performed with Immobilon Western HRP Substrate Luminol-Peroxidase kit (MerckMillipore, Cat # WBKLS0500) and the ChemiDoc MP Imaging System (Bio-Rad). Densitometry quantification was done in ImageLab software (Bio-Rad). The following antibodies were used (all Cell Signalling Technologies): GAPDH (Cat# 97166), H3K4me1 (Cat# 5326), H3K4me2 (Cat# 9725), H3K4me3 (Cat# 9751S), H3K27me1 (Cat# 84932), H3K27me3 (Cat# 9733), Histone H3 (Cat# 4499), Cleaved Histone H3 (Thr22) (Cat# 12576), DHCR24 (Cat# 2033), Rabbit IgG HRP-linked (Cat# 7074) and Mouse IgG HRP-linked (Cat# 7076). ABCAM antibodies against: H3K4me3 (Cat# ab12209), H3K9me1 (Cat# ab9045), H3K9me2 (Cat# ab1220), H3K9me3 (Cat# ab8898), H3K27me2 (Cat# ab24684), H3K36me1 (Cat# ab9048), H3K36me2 (Cat# ab9049), H3K36me3 (Cat# ab9050), Histone H3.1/3.2 (Cat# ab176840) and MVD (Cat# ab129061), Histone H3.3 (MerckMillipore Cat# ABE154).

### RT-qPCR

After RNA extraction, cDNA was generated using Applied Biosystems High-Capacity cDNA Reverse Transcription kit (Thermofisher Scientific, Cat# 4368814) as per manufacturer’s instructions. RT-qPCR was performed using Quantitect validated primers (Qiagen) or custom primers, with KAPA SYBR FAST Universal 2× qPCR Master Mix (Kapa Biosystems, (Cat# KK4602). RT-PCR was run on LightCycler 480 (Roche). The cycling condition were as follows: 10 min at 95 °C followed by 45 cycles, each consisting of 10 s at 95 °C and 30 s at 60 °C. Samples were run in triplicate. Threshold cycles (C_T_) were calculated using the LightCycler® 480 software. Relative quantification using the comparative C_T_ method was used to analyse to the data output. GAPDH were used as loading controls. Values were expressed as fold change over corresponding values for the control by the 2-ΔΔC_T_ method. The following primers were used, all from Qiagen Quantitect Primer Assay (Cat# 249900): Hs_ACTB_1_SG (Identifier: QT00095431), Hs_GAPDH_1_SG (Identifier: QT00079247), Hs_CDKN1A_SG (Identifier: QT00062090), Hs_PML_1_SG (Identifier: QT00090447), Hs_YPEL3_1_SG (Identifier: QT00078589), Hs_BHLHE41_1_SG (Identifier: QT00032697), Hs_CDKN1B_2_SG (Identifier: QT00998445), Hs_NR2F1_1_SG (Identifier: QT00089355), Hs_SOX2_1_SG (Identifier: QT00237601), Hs_SOX9_1_SG (Identifier: QT00001498), Hs_ORC1_1_SG (Identifier: QT00005341), Hs_NEK2_2_SG (Identifier: QT01668394), Hs_MCM5_1_SG (Identifier: QT00084000) and Hs_BUB1_1_SG (Identifier: QT00082929), Hs_ABCA1_SG (Identifier: QT00074606). PRDM9 custom designed primer (Forward: TGAAAGAATTGTCAAGAACAGCA; Reverse: CTCCTTCTTCCTGAGTTCCAGT) was manufactured by Integrated DNA Technologies.

### Retroviral packaging and CRISPR-Cas9 infection

SET1A, SET1B, PRDM9 and non-targeting control (NTC) sgRNAs were cloned by traditional restriction digestion into pLentiCRISPRv2 (Addgene plasmid 52961). HEK293T cells (7.2 x10^5^ per well, 6-well plate, one for each sgRNA) were seeded and once cells reached 70-90% confluency, cells were transfected using Lipofectamine 3000 (Thermofisher Scientific, Cat# L3000015) with packaging the plasmids pCAG-VSVG (Addgene plasmid 35616) and psPAX2 (Addgene plasmid 12260) and the respective sgRNA pLentiCRISPRv2 vector at a 1:3:3 ratio. 16 h after transfection the medium was replenished with fresh medium. At 48 h post transfection the lentivirus-containing supernatant was collected and filtered through a 0.45 µm ultra-low protein binding filter and concentrated using PEG according to manufacturer’s instructions. The concentrated lentivirus was stored at -80°C. RKI1 cells (7 x10^4^ cells/well) were incubated with polybrene-containing (8 μg/mL) media and lentiviral sgRNA particles for 16 hours, followed by resting the cells in fresh media for 24 hours. Cells were challenged in puromycin-containing media (1 μg/mL) for 72 hours, or until all un- transduced cells were eliminated. Puromycin-containing media was replaced with fresh media and knock-out/control cell lines were expanded. To validate gene knockout, genomic DNA was extracted from cell pellets with the ISOLATE II Genomic DNA Kit (Bioline, Cat# BIO-52067), subjected to PCR amplification and Sanger sequencing of each sgRNA target site and comparison to untransduced RKI1 cells through Synthego ICE (Supplementary Figure 8)^66^. sgRNA sequences and primers against gDNA are listed below.

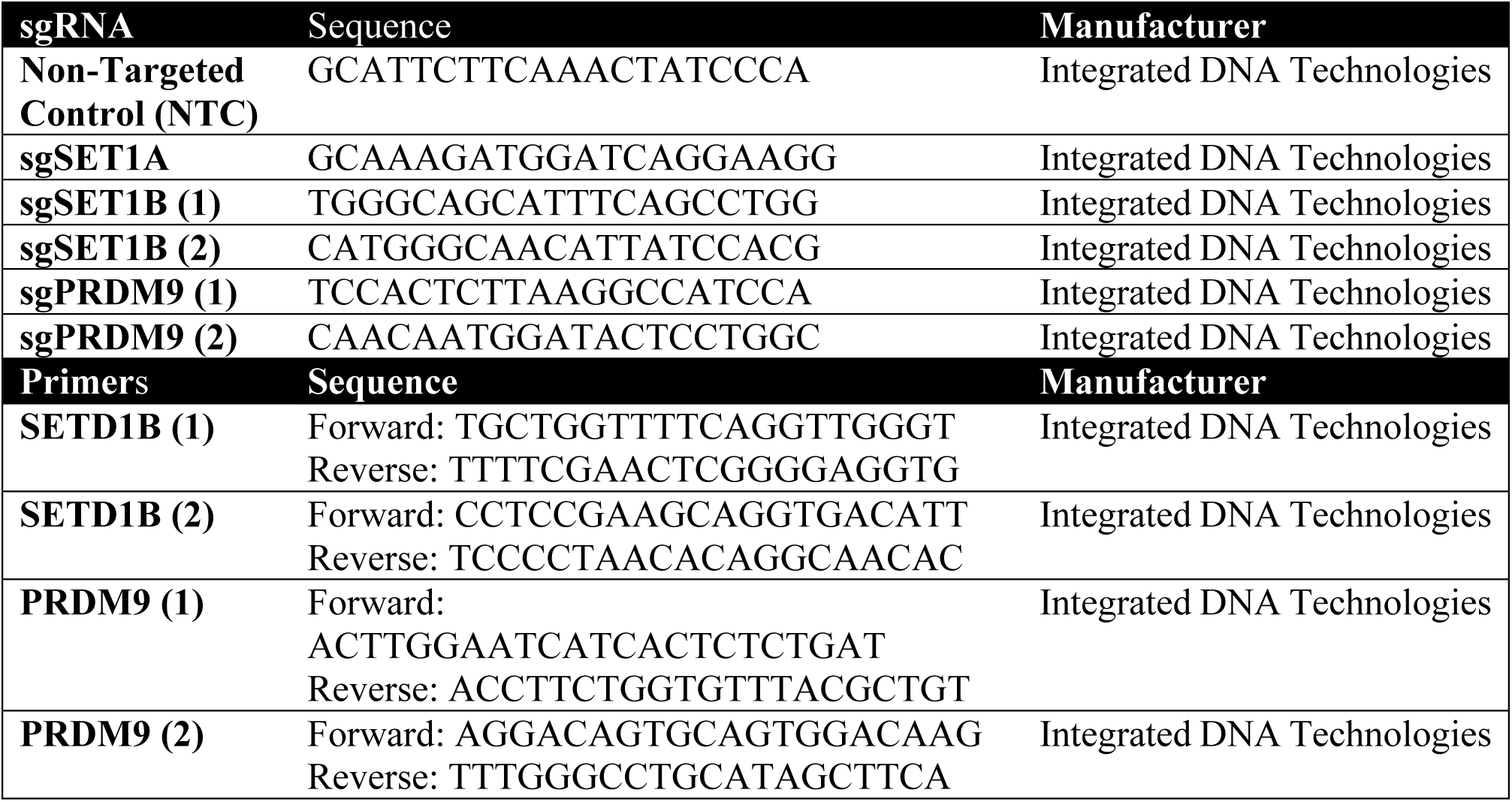

### Flow Cytometry for Reactive Oxygen Species and Lipid Peroxidation

Flow cytometric analyses of reactive oxygen species using CellROX^TM^ Green Reagent (ThermoFisher Scientific, Cat# C10444) and lipid peroxidation using Image-iT Lipid Peroxidation kit (Thermofisher Scientific, Cat# C10445) were carried out following manufacturer’s instructions. Briefly, following treatments, cells were stained for 30 min, harvested, washed in PBS and resuspended in media in flow cytometry tubes. For ROS quantification, staining intensity was assessed in the FITC channel (495/525) using a BD LSR Fortessa X-20. Lipid peroxidation was assessed in the FITC (495/525) channel for oxidised lipid species and Texas Red channels (586/603) for total lipid staining using a BD LSR Fortessa X-20.

### Tubulin polymerization assay

Tubulin polymerization assay was done using an assay kit (Cytoskeleton, Cat# BK006P) per manufacturer’s instructions. Briefly, porcine brain tubulin (10 mg/ml) was thawed in water before placing on ice and used within 2 hours. Tested compounds were incubated at 37°C with reaction mixture containing tubulin, in 55 μL reaction volume, and fluorescence over time was measured using TECAN Infite^®^ 1000 PRO plate reader (Männedorf, Switzerland).

### [^3^H]Colchicine binding assay

A radioligand competition assay was performed using the centrifugal gel filtration method. The reaction mixture (12 μL) contained 2 μM purified tubulin and 0.1 µCi (1 µl of 0.1mCi/ml, specific activity 80%) [^3^H]colchicine in PEM buffer (100 mM PIPES, 10 mM CaCl_2_, 1 mM Na-EGTA, and 1 mM MgCl_2_ [pH 6.9]). Competition binding reaction was started by the addition of 2 µL of tested compounds dissolved in dH_2_O (final concentration of 10 µM). Reaction mixtures were incubated at room temperature for 45 min, unlabelled radioactive material was filtered by using 75 μL spin column Zeba Micro Desalt Spin Column (Pierce Biotechnology) following manufacturer’s guidelines. Elute was added to the scintillation fluid to measure radioactivity of [^3^H]-colchicine bound tubulin using a scintillation counter.

### In vitro metabolic stability

All reactions were performed in 200 µL reaction volume in duplicate. 10 µM of tested compounds were incubated with human liver microsomes (0.4 mg/ml) in potassium phosphate buffer (0.1 M, pH 7.4) at 37°C with gentle shaking for 5 min. Assay was initiated by adding 12 µL of NADPH regenerating system (containing final concentrations of 1 mM NADP, 3 mM glucose-6-phosphate, 3.3 mM MgCl_2_ and 0.4u/ml glucose-6-phosphate dehydrogenase). Reaction was quenched by adding 130 µL of ice- cold methanol, vortexed vigorously and centrifuged at 15,000g at 4°C for 10 min. 5 µL of the supernatant was analysed using an Agilent 1260 LC system coupled to a QTRAP 6500 mass spectrometer. For LC, zorbax Extend-C18 (2.1 x 50 mm 1.8 um) column was used in reversed- phase mode at flow rate of 200 μL/min, with gradient elution starting with 10% of phase B (0.1% formic acid in water) and 90% of phase A (0.1% formic acid in acetonitrile). The amount of phase B was linearly increased from 10% to 90% in 5 min followed by 2 min at 90 % B then back to initial conditions at 8 min. The MS detector was operated with an ESI positive ionization mode. Source temperature and capillary voltage were set at 300°C and 4000 V, respectively. The Analyst software was used to control the instruments and data acquisition. Assaying of test compounds was carried out utilizing the mode of multiple reaction monitoring (MRM) using the following conditions for each compound. Ion transitions were 350.3 to 241 for CMPD1, 350.3 to 230 for analogue **17**, 335 to 106 for WJA88 and 335 to 185 for analogue **10**. Fragmentor voltage was set to 145 V with collision energy of 30 for all the compounds.

### Brain uptake and pharmacokinetics of WJA88

Brain uptake of WJA88 was assessed in male CD-1 mice (n = 3 per timepoint) following administration of a single bolus dose of 50 mg/kg by the intravenous route. Terminal blood and brain samples were collected from sub-groups of mice at 30 min and 60 min post-dose, the concentration of WJA88 in each sample was determined using LC-MS/MS. Pharmacokinetic parameters were assessed in male CD-1 mice (n = 3) following administration of a single 50 mg/kg dose intraperitoneally. Blood samples were collected at seven time-points up to 24 hrs, transferred into tubes containing K_2_-EDTA and WJA88 concentration in each sample was determined with LC-MS/MS. Pharmacokinetic parameters were calculated using Phoenix WinNonlin 6.3 software. All experiments and calculations were performed by WUXI, Study No. 410466-20191217-MPK.

### Data Analysis

Statistical analyses were performed using Prism 9.0 (GraphPad) or R Studio for sequencing data. The unpaired t-test was used when comparing between different treatments and one sample t-test was used when comparing between a sample and a control that the sample was normalised to. Statistical comparison of survival rates was carried out using the Mantel-Cox log-rank test. Correlation analyses were carried out using Pearson’s product moment correlation coefficient. The adjusted P-value was taken for determining significance of gene expression changes in RNA sequencing. For large data, without normal distribution, Wilcoxon rank-sum test was used. Throughout all figures: * p < 0.05; ** p < 0.01; *** p < 0.001 and **** p < 0.0001.

### Data and code availability

All sequencing datasets have been deposited into Gene Expression Omnibus (GEO) under the accession number GSE279066. The mass spectrometry proteomics data have been deposited to the ProteomeXchange Consortium via the PRIDE partner repository with the dataset identifier PXD050643. Custom RStudio scripts for RNAseq and ChIPseq data analysis will be shared upon request.

### Chemistry general experimental methods

Unless otherwise stated, reactions were conducted under positive pressure of a dry nitrogen or argon atmosphere. Temperatures of 0 °C and -10 °C were obtained using a water/ice bath or salt/ice bath respectively. Reaction mixture temperatures were reported according to the oil bath/cooling bath temperature unless otherwise stated. Anhydrous dichloromethane, triethylamine and di-isopropyl ethylamine was obtained by distillation over calcium hydride. Anhydrous DMF, methanol, THF, and acetonitrile were obtained from a PureSolv MD 7 solvent purific ation system (Innovative Technology, Inc.). Unless noted otherwise, commercially obtained reagents were used as purchased without further purification. Analytical thin-layer chromatography (TLC) was performed using Merck aluminium backed silica gel 60 F254 (0.2 mm) plates which were visualised with shortwave (254 nm) and/or longwave (365 nm) ultraviolet (UV) light, potassium permanganate, vanillin, *p*- anisaldehyde, ninhydrin or cerium molybdate (“Goofy’s Dip”) stains. Flash chromatography was performed using Grace Davisil silica gel, pore size 60 Å, 230–400 mesh particle size. Solvents for flash chromatography were distilled prior to use, or used as purchased for HPLC grade, with the eluent mixture reported as the volume/volume ratio (v/v).

Melting points were measured with open capillaries using a Stanford Research Systems (SRS) MPA160 melting point apparatus with a ramp rate of 0.5–2.0 °C/min and are uncorrected. Infrared absorption spectra were recorded on a Bruker ALPHA FT-IR spectrometer, and the data are reported as vibrational frequency (cm^−1^). Nuclear magnetic resonance spectra were recorded at 300 K unless stated otherwise, using either a Bruker AVANCE DRX200 (200 MHz), DRX300 (300 MHz), DRX400 (400.1 MHz), or AVANCE III 500 Ascend (500.1 MHz) spectrometer. The data is reported as the chemical shift (δ ppm) relative to the solvent residual peak, relative integral, multiplicity (s = singlet, d = doublet, t = triplet, q = quartet, p = pentet, m = multiplet, br = broad, dd = doublet of doublets), coupling constant (*J* Hz). Spectra for some compounds were observed as rotamers, in these cases the major peaks are reported. Low resolution mass spectra (LRMS) were recorded using electrospray ionisation (ESI) or atmospheric pressure chemical ionisation (APCI) recorded on a Finnigan LCQ ion trap spectrometer, or by electron impact gas chromatography mass spectrometry (GC/MS). High resolution mass spectra were run on a Bruker 7T Apex Qe Fourier Transform Ion Cyclotron resonance mass spectrometer equipped with an Apollo II ESI/APCI/MALDI Dual source by the Mass Spectrometry Facility of the School of Chemistry at The University of Sydney. Samples run by ESI were directly infused (150 µL/hr) using a Cole Palmer syringe pump. Samples run by APCI were injected (5 µL) into a flow of methanol (0.3 mL/min) by HPLC (Agilent 1100) coupled to the mass spectrometer. Elemental analysis was obtained from the Chemical Analysis Facility in the Department of Chemistry and Biomolecular Sciences, Macquarie University, Australia.

Analytical HPLC purity traces were taken on a Waters 2695 Separations module equipped with Waters 2996 Photodiode Array detector (set at 230, 254 and 271 nm). All samples were eluted through a Waters SunFire^™^ C18 5 µm column (2.1x150 mm) using a flow rate of 0.2 mL/min of Solvent A: MilliQ water (+0.1% trifluoroacetic acid or 0.1% formic acid) and Solvent B: acetonitrile (+0.1% trifluoroacetic acid or 0.1% formic acid). This method consisted of gradient elution (0-100% Solvent A:B over 30 min). Data acquisition and processing was performed with the Waters Empower 2 software. Reported data for all compounds are based on the 254 nm channel.

### General Synthetic Methods Suzuki Cross Coupling

Tetrakis(triphenylphosphine)palladium(0) (0.05 mol%) was added to a stirred suspension of boronic acid (1.2 eq.), Cs_2_CO_3_ (3 eq.) and 4-bromophenylbutanoic acid (1 eq.) in a degassed THF/water (9:1 v/v) mixture. The resultant mixture stirred at reflux for 8 h. Upon completion the reaction mixture was diluted with HCl (1 M, 1 x solvent volume) and extracted with EtOAc (3 x 50 mL). The combined organic phases subsequently washed with brine, before being dried (MgSO_4_), filtered and concentrated under reduced pressure. The resultant acid was purified by flash column chromatography (silica, 1:1 v/v EtOAc:Hex).

### PyBOP**^®^** Amidation

An ice-cold stirred solution of acid (1 eq)., aniline (1–1.2 eq.) and *^i^*Pr_2_Net (2 eq.) in DMF was treated with PyBOP^®^ (1 eq.), allowed to warm to room temperature and stirring continued for 12 h. The reaction mixture was diluted with CH_2_Cl_2_ (10 x solvent volume) and water (10 x solvent volume), the separated aqueous phase was subsequently extracted with CH_2_Cl_2_ (2 x with 10 x solvent volume) and combined organics washed with water (3 x with 10 x solvent volume) sat. aq. Solution of NaHCO_3_ (5 x solvent volume) and brine (100 mL) before being dried (MgSO_4_), filtered and purified *via* flash column chromatography (silica, 1:1 v/v EtOAc:Hex) to give the required amides.

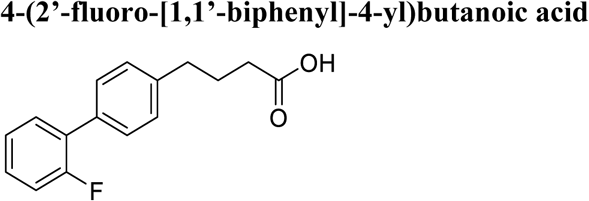

According to the general procedure for Suzuki Cross couplings, the title compound was synthesised from 2-fluorophenylboronic acid (1.70 g, 12.3 mmol, 1.2 eq.), and 4-bromophenylbutanoic acid (2.50 g, 10.2 mmol, 1.0 eq.) in dry degassed THF/water (100 mL, 9:1 v/v mixture). The crude product was purified by flash column chromatography (silica, 1:1 v/v EtOAc:Hex) to give the title compound as a white crystalline solid (1.6 g, 61%).

R_f_: 0.22 (SiO_2_, 1:1 EtOAc:Hex), ^1^H NMR (300 MHz, DMSO-*d*_6_) δ 12.06 (br s, 1H), 7.57 – 7.23 (m, 8H), 2.64 (t, *J* = 7.4 Hz, 2H), 2.25 (t, *J* = 7.4 Hz, 2H), 1.88 – 1.80 (m, 2H). ^13^C NMR (75 MHz, DMSO-*d*_6_) δ 174.2, 159.1 (d, ^1^*J*_CF_ = 245.8 Hz), 141.3, 132.6 (2C), 130.7 (d, ^4^*J*_CF_ = 3.7 Hz), 129.3 (d, ^3^*J*_CF_ = 8.5 Hz), 128.7 (d, ^4^*J*_CF_ = 3.0 Hz), 128.6 (2C), 128.2 (d, ^3^*J*_CF_ = 13.9 Hz), 124.9 (d, ^4^*J*_CF_ = 3.7 Hz), 116.0 (d, ^2^*J*_CF_ = 22.6 Hz), 34.1, 33.1, 26.2. ^19^F NMR (282 MHz, DMSO-*d*_6_) δ -118.4. IR (diamond cell, neat) ϖ_max_: 3026, 2902, 1696, 1480, 1435, 1250, 1202, 939, 799, 756, 563 cm^−1^. LRMS (ESI-) m/z: 257 [(M-H)^-^, 100%]

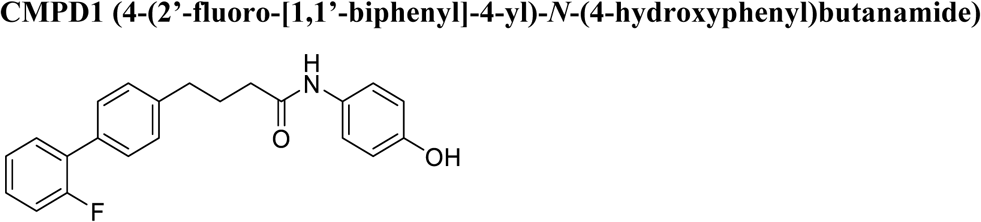

According to the general procedure for Suzuki Cross couplings, the title compound was synthesised from 2-fluorophenylboronic acid (84.0 g, 0.6 mmol, 1.0 eq.), and 4-(4-bromophenyl)-N-(4- hydroxyphenyl)butanamide (200 mg, 0.6 mmol, 1.0 eq.) in dry degassed THF/water (4 mL, 9:1 v/v mixture). The crude product was purified by flash column chromatography (silica, 1:1 v/v EtOAc:Hex) to give the title compound as a white solid (150 mg, 71%).

R_f_: 0.38 (SiO_2_, 1:1 EtOAc:Hex), ^1^H NMR (300 MHz, Chloroform-*d*) δ 7.49 (dd, *J* = 8.1, 1.7 Hz, 2H), 7.43 (td, J = 7.7, 1.9 Hz, 1H), 7.34 – 7.27 (m, 5H), 7.23 – 7.11 (m, 2H), 7.01 (s, 1H), 6.76 (d, *J* = 8.9 Hz, 2H), 5.25 (s, 1H), 2.77 (t, *J* = 7.4 Hz, 2H), 2.36 (t, *J* = 7.4 Hz, 2H), 2.12 (p, *J* = 7.4 Hz, 2H).^13^C NMR (75 MHz, Chloroform-*d*) δ 171.2, 159.9 (d, ^1^*J*_CF_ = 247.6 Hz), 152.9, 141.0, 133.8, 130.8 (d, ^4^*J*_CF_ = 3.7 Hz, 2C), 130.7, 129.2 (d, ^3^*J*_CF_ = 3.1 Hz), 128.9 (d, ^3^*J*_CF_ = 8.2 Hz), 128.8 (2C), 124.5 (d, ^4^*J*_CF_ =3.8 Hz), 122.5 (2C), 116.2 (d, ^2^*J*_CF_ = 22.9 Hz), 115.9 (2C), 36.7, 34.9, 26.9. ^19^F NMR (282 MHz, Chloroform-*d*) δ -118.1. IR (diamond cell, neat) ϖ_max_: 3317, 2965, 2816, 1654, 1514, 1350, 1230, 1100, 880 cm^−1^. LRMS (ESI+) m/z: 372 [(M+Na)^+^, 100%]. HRMS (ESI+) calcd for C_22_H_20_FNNaO_2_ (M+Na)^+^, 372.13703; Found 372.13687.

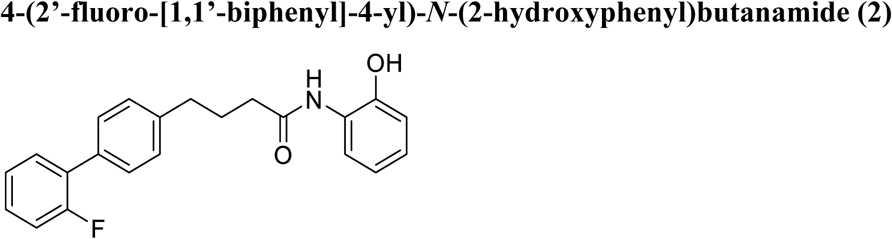

According to the general procedure for PyBOP^®^ amidation, the title compound was synthesised from 4-(2’-fluoro-[1,1’-biphenyl]-4-yl)butanoic acid (130 mg, 0.5 mmol, 1.0 eq.), and 2-hydroxyaniline (58.6 mg, 0.5 mmol, 1.0 eq.) in dry DMF (5 mL). The crude product was purified by flash column chromatography (silica, 1:1 v/v EtOAc:Hex) to give the title compound as a yellow oil (126 mg, 69%).

R_f_: 0.51 (SiO_2_, 1:1 EtOAc:Hex), m.p.: 53 – 55 °C, ^1^H NMR (500 MHz, DMSO-*d*_6_) δ 9.72 (s, 1H), 9.29 (s, 1H), 7.69 – 7.66 (m, 1H), 7.53 – 7.37 (m, 4H), 7.34 – 7.17 (m, 4H), 6.96 – 6.93 (m, 1H), 6.88 – 6.86 (m, 1H), 6.78 – 6.75 (m, 1H), 2.70 – 2.58 (m, 2H), 2.47 – 2.38 (m, 2H), 1.98 – 1.84 (m, 2H). ^13^C NMR (126 MHz, DMSO-*d*_6_) δ 171.6, 159.1 (d, ^1^*J*_CF_ = 245.7 Hz), 148.0, 141.5, 132.6, 131.1, 130.6 (d, ^4^*J*_CF_ = 2.7 Hz, 2C), 129.2 (d, ^3^*J*_CF_ = 8.2 Hz), 128.7 (d, ^4^*J*_CF_ = 2.9 Hz), 128.6 (2C), 128.2 (d, ^3^*J*_CF_ = 13.2 Hz), 126.4, 124.9 (d, ^4^*J*_CF_ = 3.5 Hz), 124.7, 122.5, 119.0, 116.1, 116.0 (d, ^4^*J*_CF_ = 2.7 Hz), 35.4, 34.3, 26.9. ^19^F NMR (471 MHz, DMSO-*d*_6_) δ -118.4. IR (diamond cell, neat) ϖ_max_: 3175, 1633, 1594, 1524, 1483, 1453, 1362, 1283, 1103, 821, 744, 458 cm^−1^. LRMS (ESI+) m/z: 372 [(M+Na)^+^, 100%].

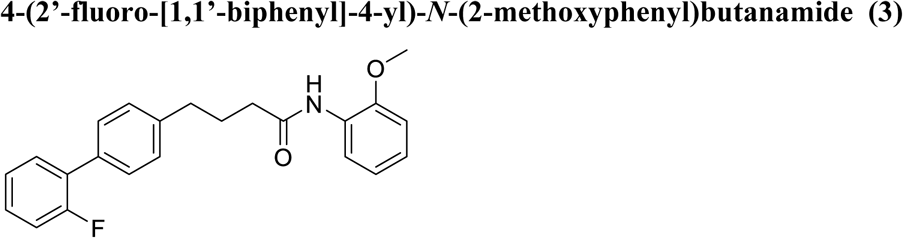

According to the general procedure for PyBOP^®^ amidation, the title compound was synthesised from 4-(2’-fluoro-[1,1’-biphenyl]-4-yl)butanoic acid (130 mg, 0.5 mmol, 1.0 eq), and 2-methoxyaniline (61.6 mg, 0.5 mmol, 1.0 eq) in dry DMF (5 mL). The crude product was purified by flash column chromatography (silica, 1:1 v/v EtOAc:Hex) to give the title compound as a yellow oil (126 mg, 69%).

R_f_: 0.41 (SiO_2_, 1:1 EtOAc:Hex), ^1^H NMR (500 MHz, DMSO-*d*_6_) δ 9.07 (s, 1H), 7.94 (t, *J* = 9.1 Hz, 1H), 7.57 – 7.34 (m, 4H), 7.34 – 7.13 (m, 4H), 7.11 – 6.96 (m, 2H), 6.95 – 6.84 (m, 1H), 3.82 (s, 3H), 2.75 – 2.55 (m, 2H), 2.48 – 2.33 (m, 2H), 2.05 – 1.81 (m, 2H). ^13^C NMR (126 MHz, DMSO-*d*_6_) δ 171.1, 159.1 (d, ^1^*J*_CF_ = 245.6 Hz), 149.7, 141.5, 132.6, 130.6 (2C), 129.2 (d, ^3^*J*_CF_ = 8.3 Hz), 128.7 (d, ^4^*J*_CF_ = 2.8 Hz), 128.6 (2C), 128.2 (d, ^3^*J*_CF_ = 13.2 Hz), 127.4, 124.9 (d, ^4^*J*_CF_ = 3.5 Hz), 124.2, 122.2, 120.1, 116.0 (d, ^2^*J*_CF_ = 22.7 Hz), 111.1, 55.6, 35.6, 34.3, 26.8. ^19^F NMR (471 MHz, DMSO-*d*_6_) δ - 118.4. IR (diamond cell, neat) ϖ_max_: 3417, 3311, 2934, 1676, 1599, 1518, 1482, 1457, 1432, 1288, 1249, 1216, 1115, 1026, 745 cm^−1^. LRMS (ESI+) m/z: 386 [(M+Na)^+^, 100%].

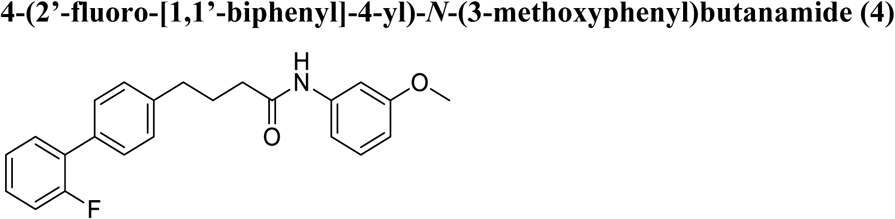

According to the general procedure for PyBOP^®^ amidation, the title compound was synthesised from 4-(2’-fluoro-[1,1’-biphenyl]-4-yl)butanoic acid (130 mg, 0.5 mmol, 1.0 eq.), and 3-methoxyaniline (61.6 mg, 0.5 mmol, 1.0 eq.) in dry DMF (5 mL). The crude product was purified by flash column chromatography (silica, 1:1 v/v EtOAc:Hex) to give the title compound as a colourless oil (151 mg, 83%).

R_f_: 0.41 (SiO_2_, 1:1 EtOAc:Hex), ^1^H NMR (500 MHz, DMSO-*d*_6_) δ 9.88 (s, 1H), 7.56 – 7.24 (m, 8H), 7.24 – 7.07 (m, 3H), 6.72 – 6.49 (m, 1H), 3.71 (s, 3H), 2.73 – 2.56 (m, 2H), 2.43 – 2.24 (m, 2H), 2.00 – 1.83 (m, 2H). ^13^C NMR (126 MHz, DMSO-*d*_6_) δ 171.0, 159.1 (d, ^1^*J*_CF_ = 245.6 Hz), 141.4, 140.5, 132.6, 131.1, 130.6 (d, ^4^*J*_CF_ = 2.7 Hz, 2C), 129.4, 129.2 (d, ^3^*J*_CF_ = 8.2 Hz), 128.7 (d, ^4^*J*_CF_ = 2.9 Hz), 128.6 (2C), 128.2 (d, ^3^*J*_CF_ = 13.1 Hz), 124.9 (d, ^4^*J*_CF_ = 3.4 Hz), 116.0 (d, ^2^*J*_CF_ = 22.5 Hz), 111.4, 108.4, 104.9, 54.9, 35.8, 34.3, 26.5. ^19^F NMR (471 MHz, DMSO-*d*_6_) δ -118.4. IR (diamond cell, neat) ϖ_max_: 1659, 1597, 1543, 1483, 1451, 1416, 1284, 1209, 1155, 1042, 755, 687, 565 cm^−1^. LRMS (ESI+) m/z: 386 [(M+Na)^+^, 100%].

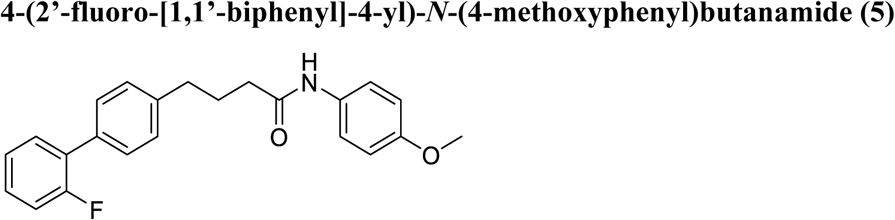

According to the general procedure for PyBOP^®^ amidation, the title compound was synthesised from 4-(2’-fluoro-[1,1’-biphenyl]-4-yl)butanoic acid (130 mg, 0.5 mmol, 1.0 eq.), and 4-methoxyaniline (61.6 mg, 0.5 mmol, 1.0 eq.) in dry DMF (5 mL). The crude product was purified by flash column chromatography (silica, 1:1 v/v EtOAc:Hex) to give the title compound as a pink powder (133 mg, 73%).

R_f_: 0.40 (SiO_2_, 1:1 EtOAc:Hex), m.p.: 103 – 106 °C, ^1^H NMR (300 MHz, DMSO-*d*_6_) δ ^1^H NMR (500 MHz, DMSO-*d*_6_) δ 9.75 – 9.72 (m, 1H), 7.53 – 7.45 (m, 5H), 7.40 – 7.16 (m, 5H), 6.86 (d, *J* = 9.1 Hz, 2H), 3.71 (s, 3H), 2.69 – 2.58 (m, 2H), 2.34 – 2.26 (m, 2H), 1.97 – 1.84 (m, 2H). ^13^C NMR (126 MHz, DMSO-*d*_6_) δ 170.4, 159.1 (d, ^1^*J*_CF_ = 245.6 Hz), 155.0, 141.5, 132.6 (d, ^3^*J*_CF_ = 9.6 Hz), 131.1, 130.6 (2C), 129.2 (d, ^3^*J*_CF_ = 8.4 Hz), 128.7 (d, ^4^*J*_CF_ = 2.9 Hz), 128.6 (2C), 124.9 (d, ^4^*J*_CF_ = 3.5 Hz), 120.6 (2C), 116.1 (d, ^2^*J*_CF_ = 22.7 Hz), 113.8 (2C), 55.1, 35.7, 34.4, 26.7. ^19^F NMR (471 MHz, DMSO-*d*_6_) δ -118.4. IR (diamond cell, neat) ϖ_max_: 3283, 2914, 1646, 1510, 1483, 1409, 1245, 1106, 1030, 827, 802, 753, 601, 568, 521 cm^−1^. LRMS (ESI+) m/z: 386 [(M+Na)^+^, 100%].

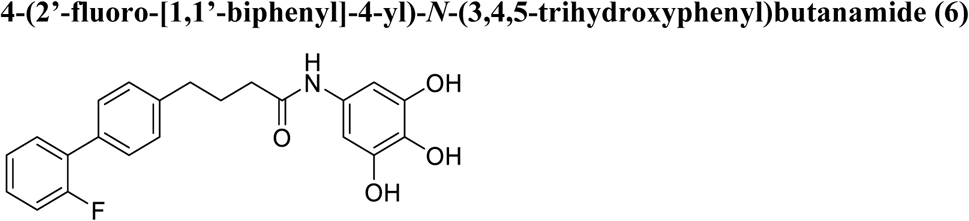

BBr_3_ (1.75 mL of a 1 M soln. in CH_2_Cl_2_, 1.75 mmol, 5.0 eq.) was added dropwise to an ice cold, stirring solution of 4-(2’-fluoro-[1,1’-biphenyl]-4-yl)-*N*-(3,4,5-trimethoxyphenyl)butanamide (150 mg, 0.35 mmol, 1.0 eq.) in dry CH_2_Cl_2_ (5 mL) over 15 minutes. The resultant solution was warmed to room temperature and stirring continued for 8 h. The reaction was then diluted with CH_2_Cl_2_ (20 mL), washed with water (25 mL), NaHCO_3_ (25 mL of a sat. aq. Soln.) before being dried (MgSO_4_), filtered and concentrated under reduced pressure. The crude mass was purified by flash column chromatography (silica, 1:1 v/v EtOAc:Hex) to give the title compound as a white crystalline solid (116 mg, 87%).

R_f_: 0.09 (SiO_2_, 1:1 EtOAc:Hex), m.p.: 168 – 170 °C, ^1^H NMR (500 MHz, DMSO-*d*_6_) δ 9.41 (s, 1H), 8.76 (s, 2H), 7.73 (s, 1H), 7.53 – 7.37 (m, 4H), 7.32 – 7.27 (m, 4H), 6.63 (s, 2H), 2.66 – 2.55 (m, 2H), 2.28 – 2.20 (m, 2H), 1.93 – 1.82 (m, 2H). ^13^C NMR (126 MHz, DMSO-*d*_6_) δ 170.0, 159.1 (d, ^1^*J*_CF_ = 245.6 Hz), 145.8 (2C), 141.5, 132.6, 131.1, 130.9, 130.7 (d, ^4^*J*_CF_ = 3.5 Hz, 2C), 129.3 (d, ^3^*J*_CF_ = 8.3 Hz), 128.7 (d, ^4^*J*_CF_ = 2.8 Hz), 128.6 (2C), 128.2 (d, ^3^*J*_CF_ = 13.0 Hz), 124.9 (d, ^4^*J*_CF_ = 3.7 Hz), 116.1 (d, ^2^*J*_CF_ = 22.6 Hz), 99.0 (2C), 35.8, 34.4, 26.8. ^19^F NMR (471 MHz, DMSO-*d*_6_) δ -118.4. IR (diamond cell, neat) ϖ_max_: 3520, 3370, 3119, 1628, 1541, 1483, 1448, 1374, 1336, 1290, 1190, 1048, 852, 822, 758, 567 cm^−1^. LRMS (ESI+) m/z: 404 [(M+Na)^+^, 100%].

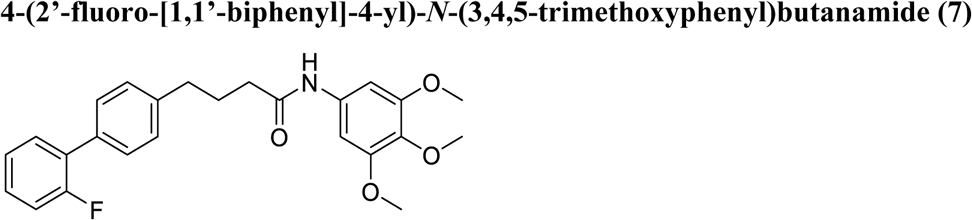

According to the general procedure for PyBOP^®^ amidation, the title compound was synthesised from 4-(2’-fluoro-[1,1’-biphenyl]-4-yl)butanoic acid (300 mg, 1.16 mmol, 1.0 eq.), and 3,4,5- methoxyaniline (216 mg, 1.16 mmol, 1.0 eq.) in dry DMF (10 mL). The crude product was purified by flash column chromatography (silica, 1:1 v/v EtOAc:Hex) to give the title compound as a white crystalline solid (275 mg, 56%).

R_f_: 0.37 (SiO_2_, 1:1 EtOAc:Hex), m.p.: 130 – 132 °C, ^1^H NMR (500 MHz, DMSO-*d*_6_) δ 9.84 (s, 1H), 7.51 – 7.36 (m, 4H), 7.33 – 7.26 (m, 4H), 7.03 (s, 2H), 3.74 (s, 6H), 3.62 (s, 3H), 2.68 (t, *J* = 7.6 Hz, 2H), 2.35 (t, *J* = 7.4 Hz, 2H), 1.98 – 1.92 (m, 2H). ^13^C NMR (126 MHz, DMSO-*d*_6_) δ 170.8, 159.1 (d, ^1^*J*_CF_ = 245.7 Hz), 152.7 (2C), 141.4, 135.5, 133.2, 132.6, 130.6 (d, ^4^*J*_CF_ = 3.3 Hz, 2C), 129.2 (d, ^3^*J*_CF_ = 8.4 Hz), 128.7 (d, ^4^*J*_CF_ = 2.9 Hz), 128.6 (2C), 128.2 (d, ^3^*J*_CF_ = 13.2 Hz), 124.9 (d, ^4^*J*_CF_ = 3.5 Hz), 116.0 (d, ^2^*J*_CF_ = 22.6 Hz), 96.8 (2C), 60.0, 55.6 (2C), 35.9, 34.3, 26.5. ^19^F NMR (471 MHz, DMSO-*d*_6_) δ -118.4. IR (diamond cell, neat) ϖ_max_: 3331, 2937, 1686, 1610, 1544, 1509, 1482, 1446, 1222, 1130, 985, 843, 819, 766, 637, 456 cm^−1^. LRMS (ESI+) m/z: 446 [(M+Na)^+^, 100%].

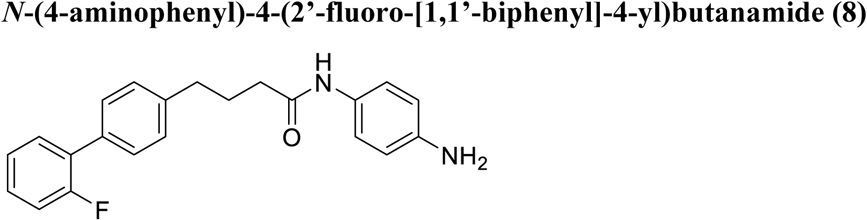

A stirring suspension of 4-(2’-fluoro-[1,1’-biphenyl]-4-yl)-*N*-(4-nitrophenyl)butanamide (250 mg, 0.66 mmol, 1.0 eq.) and palladium on carbon (10% w/w, 50 mg) in EtOAc (10 mL) was placed under an atmosphere of hydrogen (1 atm.) and stirring continued for 18 h. The reaction volume was filtered through a Celite & basic alumina plug. The residue was washed with EtOAc (3 x 20 mL) and combined filtrates concentrated under reduced pressure. The resultant product was purified by flash column chromatography (silica, 1:1 v/v EtOAc:Hex) to give the title compound as a white crystalline solid (141 mg, 61%).

R_f_: 0.26 (SiO_2_, 1:1 EtOAc:Hex), m.p.: 100 – 102 °C, ^1^H NMR (500 MHz, DMSO-*d*_6_) δ 9.45 (s, 1H), 7.53 – 7.37 (m, 4H), 7.32 (d, *J* = 8.1 Hz, 2H), 7.28 (t, *J* = 7.1 Hz, 1H), 7.23 (d, *J* = 8.6 Hz, 2H), 6.50 (d, *J* = 8.6 Hz, 2H), 4.81 (br s, 2H), 2.66 (t, *J* = 7.6 Hz, 2H), 2.28 (t, *J* = 7.4 Hz, 2H), 1.92 (p, *J* = 7.6 Hz, 2H) . ^13^C NMR (126 MHz, DMSO-*d*_6_) δ 169.8, 159.1 (d, ^1^*J*_CF_ = 245.5 Hz), 144.5, 141.5, 132.6, 130.6 (d, ^4^*J*_CF_ = 3.4 Hz, 2C), 129.3 (d, ^3^*J*_CF_ = 8.2 Hz), 129.1, 128.7 (d, ^4^*J*_CF_ = 2.9 Hz), 128.6 (2C), 128.2 (d, ^3^*J*_CF_ = 13.1 Hz), 124.9 (d, ^4^*J*_CF_ = 3.5 Hz), 120.9 (2C), 116.0 (d, ^2^*J*_CF_ = 22.6 Hz), 113.8 (2C), 35.6, 34.4, 26.8. ^19^F NMR (471 MHz, DMSO-*d*_6_) δ -118.4. IR (diamond cell, neat) ϖ_max_: 3457, 3374, 3276, 1641, 1536, 1514, 1482, 1424, 1275, 1252, 1204, 943, 823, 801, 755, 569, 510, 471 cm^−1^. LRMS (ESI+) m/z: 371 [(M+H)^+^, 100%].

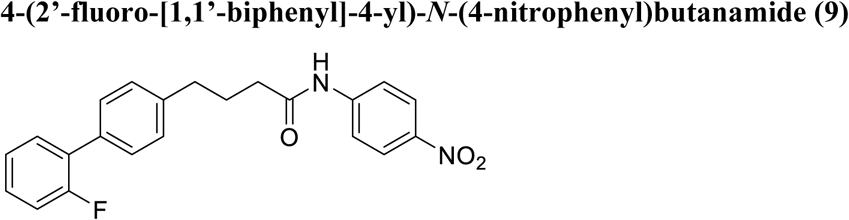

According to the general procedure for PyBOP^®^ amidation, the title compound was synthesised from 4-(2’-fluoro-[1,1’-biphenyl]-4-yl)butanoic acid (420 mg, 1.6 mmol, 1.03 eq), and 4-nitroaniline (213 mg, 1.55 mmol, 1 eq) in dry DMF (10 mL). The crude product was purified by flash column chromatography (silica, 1:1 v/v EtOAc:Hex) to give the title compound as a yellow powder (473 mg, 78%).

R_f_: 0.33 (SiO_2_, 1:1 EtOAc:Hex), m.p.: 118 – 120 °C, ^1^H NMR (400 MHz, Chloroform-*d*) δ 8.17 – 8.04 (m, 2H), 7.87 (s, 1H), 7.70 – 7.67 (m, 2H), 7.48 – 7.46 (m, 2H), 7.39 (td, *J* = 7.7, 1.9 Hz, 1H), 7.32 – 7.27 (m, 1H), 7.26 – 7.24 (m, 2H), 7.21 – 7.11 (m, 2H), 2.76 (t, *J* = 7.3 Hz, 2H), 2.42 (t, *J* = 7.4 Hz, 2H), 2.15 – 2.08 (m, 2H). ^13^C NMR (101 MHz, Chloroform-*d*) δ 171.7, 159.9 (d, ^1^*J*_CF_ = 247.1 Hz), 144.1, 143.4, 140.7, 133.8, 130.7 (d, ^4^*J*_CF_ = 3.5 Hz, 2C), 129.2 (d, ^4^*J*_CF_ = 2.9 Hz), 129.0 (d, ^3^*J*_CF_ = 8.3 Hz), 128.7 (2C), 125.2 (2C), 124.5 (d, ^4^*J*_CF_ = 3.6 Hz), 119.1, 119.0 (2C), 116.2 (d, ^2^*J*_CF_ = 22.8 Hz), 36.8, 34.8, 26.5. ^19^F NMR (376 MHz, Chloroform-*d*) δ -118.2. IR (diamond cell, neat) ϖ_max_: 1661, 1595, 1502, 1450, 1406, 1342, 1255, 1208, 1107, 858, 748, 564, 496 cm^−1^. LRMS (ESI-) m/z: 377 [(M-H)^-^, 100%].

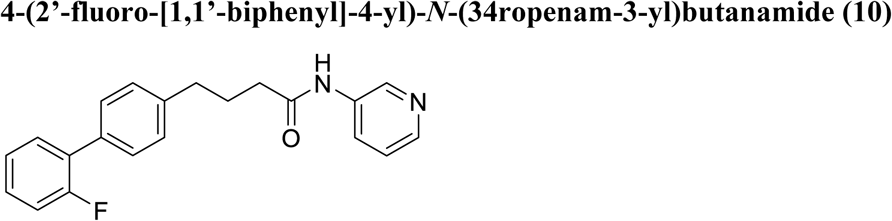

An ice cold magnetically stirred solution of 4-(2’-fluoro-[1,1’-biphenyl]-4-yl)butanoic acid (200 mg, 0.77 mmol, 1.0 eq), 3-aminopyridine (80 mg, 0.85 mmol, 1.1 eq) and *^i^*Pr_2_Net (268 μL, 1.54 mmol, 2.0 eq) in DMF (5 mL) was treated with PyBOP^®^ (400 mg, 0.77 mmol, 1.0 eq), allowed to warm to room temperature and stirring continued for 12 h. The reaction mass was diluted with CH_2_Cl_2_ (50 mL) and water (50 mL), the separated organic phase was subsequently washed with NaHCO_3_ (25 mL of a sat. aq. Solution) and brine (100 mL) before being dried (MgSO_4_), filtered and purified *via* flash column chromatography (silica, 1:1 v/v EtOAc:Hex) to give the title compound as a white solid (169 mg, 64%).

R_f_: 0.14 (SiO_2_, 1:1 EtOAc:Hex), m.p.: 78 – 80 °C, ^1^H NMR (400 MHz, DMSO-*d*_6_) δ 10.11 (s, 1H), 8.73 (d, *J* = 2.6 Hz, 1H), 8.23 (dd, *J* = 4.7, 1.5 Hz, 1H), 8.03 (ddd, *J* = 8.3, 2.5, 1.6 Hz, 1H), 7.52 – 7.46 (m, 3H), 7.42 – 7.37 (m, 1H), 7.34 – 7.26 (m, 5H), 2.69 (t, *J* = 7.5 Hz, 2H), 2.39 (t, *J* = 7.5 Hz, 2H), 1.95 (p, *J* = 7.5 Hz, 2H). ^13^C NMR (101 MHz, DMSO-*d*_6_) δ 171.6, 159.1 (d, ^1^*J*_CF_ = 245.6 Hz), 144.0, 141.4, 140.8, 135.9, 132.7, 130.7 (d, ^4^*J*_CF_ = 3.5 Hz, 2C), 129.3 (d, ^3^*J*_CF_ = 8.4 Hz), 128.8 (d, ^4^*J*_CF_ = 2.9 Hz), 128.7 (2C), 128.2 (d, ^3^*J*_CF_ = 13.2 Hz), 126.0, 124.9 (d, ^4^*J*_CF_ = 3.6 Hz), 123.6, 116.1 (d, ^2^*J*_CF_ = 22.6 Hz), 35.6, 34.3, 26.4. ^19^F NMR (471 MHz, DMSO) δ -118.4. IR (diamond cell, neat) ϖ_max_: 3333, 2936, 1644, 1605, 1483, 1431, 1298, 1237, 1214, 1134, 1023, 866, 799, 760, 601, 567, 472 cm^−1^. LRMS (ESI+) m/z: 335 [(M+H)^+^, 25%], 357 [(M+Na)^+^, 100%].

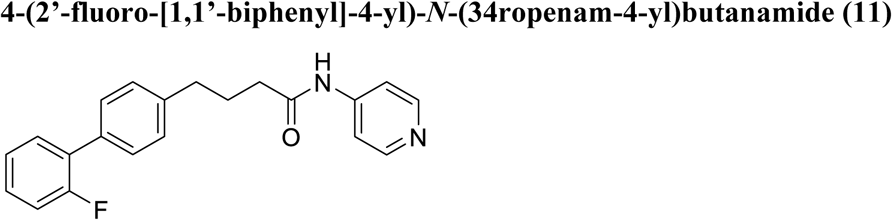

An ice cold magnetically stirred solution of 4-(2’-fluoro-[1,1’-biphenyl]-4-yl)butanoic acid (200 mg, 0.77 mmol, 1.0 eq.), 4-aminopyridine (80 mg, 0.85 mmol, 1.1 eq.) and *^i^*Pr_2_Net (268 μL, 1.54 mmol, 2.0 eq.) in DMF (5 mL) was treated with PyBOP^®^ (400 mg, 0.77 mmol, 1.0 eq.), allowed to warm to room temperature and stirring continued for 12 h. The reaction mass was diluted with CH_2_Cl_2_ (50 mL) and water (50 mL), the separated organic phase was subsequently washed with NaHCO_3_ (25 mL of a sat. aq. Solution) and brine (100 mL) before being dried (MgSO_4_), filtered and purified *via* flash column chromatography (silica, 1:1 v/v EtOAc:Hex) to give the title compound as a white solid (228 mg, 86%).

R_f_: 0.11 (SiO_2_, 1:1 EtOAc:Hex), m.p.: 108 – 110 °C, ^1^H NMR (400 MHz, DMSO-*d*_6_) δ 11.92 (s, 1H), 8.67 (d, *J* = 7.2 Hz, 2H), 8.16 (d, *J* = 7.3 Hz, 2H), 7.50 – 7.42 (m, 3H), 7.41 – 7.35 (m, 1H), 7.33 – 7.13 (m, 4H), 2.73 – 2.64 (m, 2H), 2.58 (t, *J* = 7.3 Hz, 2H), 2.02 – 1.92 (m, 2H). ^13^C NMR (101 MHz, DMSO-*d*_6_) δ 173.8, 159.1 (d, ^1^*J*_CF_ = 245.6 Hz), 153.0, 142.0 (2C), 141.2, 132.7, 131.1, 130.6 (d, ^4^*J*_CF_ = 3.4 Hz, 2C), 129.3 (d, ^3^*J*_CF_ = 8.4 Hz), 128.7 (d, ^4^*J*_CF_ = 2.9 Hz), 128.6 (2C), 128.1 (d, ^3^*J*_CF_ = 13.2 Hz), 124.9 (d, ^4^*J*_CF_ = 3.6 Hz), 116.1 (d, ^2^*J*_CF_ = 22.5 Hz), 114.2 (2C), 36.2, 34.1, 25.9. ^19^F NMR (376 MHz, DMSO-*d*_6_) δ -118.4. IR (diamond cell, neat) ϖ_max_: 2926, 1716, 1561, 1500, 1483, 1313, 1135, 823, 754, 514 cm^−1^. LRMS (ESI+) m/z: 335 [(M+H)^+^, 100%], 357 [(M+Na)^+^, 40%].

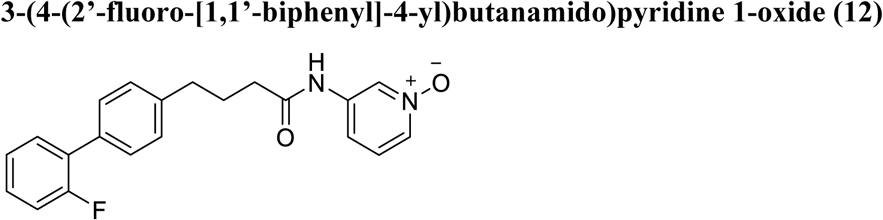

An ice cold magnetically stirred solution of 4-(2’-fluoro-[1,1’-biphenyl]-4-yl)-*N*-(35ropenam-3- yl)butanamide (100 mg, 0.30 mmol, 1 eq.) in dry CH_2_Cl_2_ (10 mL) was treated with mCPBA (3 eq.), allowed to warm to room temperature and stirring continued for 12 h. The reaction mass was diluted with CH_2_Cl_2_ (50 mL) and water (50 mL), the separated organic phase was subsequently washed with NaHCO_3_ (25 mL of a sat. aq. Solution) and brine (100 mL) before being dried (MgSO_4_), filtered and purified *via* flash column chromatography (silica, 1:1 v/v EtOAc:Hex) to give the title compound as a white solid (80 mg, 76%).

R_f_: 0.10 (SiO_2_, 1:1 EtOAc:Hex), m.p.: 58 – 60 °C, ^1^H NMR (500 MHz, DMSO-*d*_6_) δ 10.29 (s, 1H), 8.71 (s, 1H), 7.94 – 7.93 (m, 1H), 7.51 – 7.46 (m, 3H), 7.41 – 7.36 (m, 2H), 7.36 – 7.26 (m, 5H), 2.67 (t, *J* = 7.5 Hz, 2H), 2.38 (t, *J* = 7.4 Hz, 2H), 1.94 (p, *J* = 7.6 Hz, 2H). ^13^C NMR (126 MHz, DMSO-*d*_6_) δ 171.8, 159.1 (d, ^1^*J*_CF_ = 245.7 Hz), 141.3, 138.4, 133.5, 132.7, 130.6 (d, ^4^*J*_CF_ = 3.4 Hz, 2C), 129.9, 129.3 (d, ^3^*J*_CF_ = 8.4 Hz), 128.8 (d, ^4^*J*_CF_ = 2.9 Hz), 128.6 (2C), 128.2 (d, ^3^*J*_CF_ = 13.1 Hz), 126.1, 124.9 (d, ^4^*J*_CF_ = 3.4 Hz), 116.1 (d, ^2^*J*_CF_ = 22.6 Hz), 115.7, 35.6, 34.2, 26.2. ^19^F NMR (471 MHz, DMSO-*d*_6_) δ -118.4. IR (diamond cell, neat) ϖ_max_: 2928, 1693, 1575, 1547, 1482, 1415, 1284, 1207, 1146, 980, 789, 751, 672, 590, 550 cm^−1^. LRMS (ESI+) m/z: 373 [(M+Na)^+^, 100%].

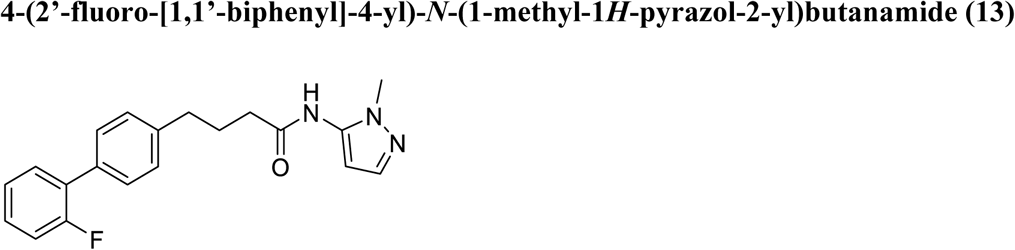

According to the general procedure for PyBOP^®^ amidation, the title compound was synthesised from 4-(2’-fluoro-[1,1’-biphenyl]-4-yl)butanoic acid (150 mg, 0.58 mmol, 1.0 eq.), and aminopyrazole (62 mg, 0.64 mmol, 1.1 eq.) in dry DMF (10 mL). The crude product was purified by flash column chromatography (silica, 1:1 v/v EtOAc:Hex) to give the title compound as a gummy solid (110 mg, 56%).

R_f_: 0.11 (SiO_2_, 1:1 EtOAc:Hex), ^1^H NMR (500 MHz, DMSO-*d*_6_) δ 9.88 (s, 1H), 7.53 – 7.48 (m, 3H), 7.42 – 7.37 (m, 1H), 7.34 – 7.27 (m, 5H), 6.18 (d, *J* = 1.9 Hz, 1H), 3.65 (s, 3H), 2.69 (t, *J* = 7.6 Hz, 2H), 2.40 (t, *J* = 7.5 Hz, 2H), 1.95 (p, *J* = 7.6 Hz, 2H). ^13^C NMR (126 MHz, DMSO-*d*_6_) δ 170.9, 159.1 (d, ^1^*J*_CF_ = 245.6 Hz), 141.3, 137.3, 136.5, 132.7, 130.6 (d, ^4^*J*_CF_ = 3.4 Hz, 2C), 129.3 (d, ^3^*J*_CF_ = 8.4 Hz), 128.7 (d, ^4^*J*_CF_ = 2.9 Hz), 128.6 (2C), 128.2 (d, ^3^*J*_CF_ = 13.0 Hz), 124.9 (d, ^4^*J*_CF_ = 3.4 Hz), 116.1 (d, ^2^*J*_CF_ = 22.5 Hz), 98.7, 35.5, 34.7, 34.2, 26.4. ^19^F NMR (471 MHz, DMSO-*d*_6_) δ -118.4. IR (diamond cell, neat) ϖ_max_: 3268, 1663, 1543, 1482, 1249, 1194, 1106, 968, 928, 822, 800, 758, 691, 565, 532, 461cm^−1^. LRMS (ESI+) m/z: 360 [(M+Na)^+^, 100%].

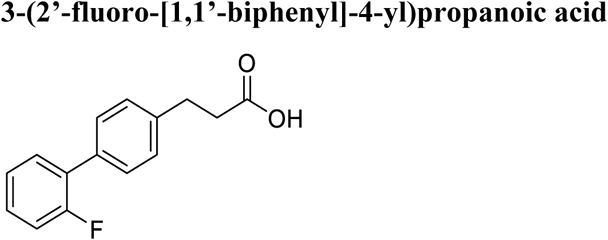

Tetrakis(triphenylphosphine)palladium(0) (554.7 mg, 0.48 mmol, 0.05 mol%) was added to a stirred suspension of 2-fluorophenylboronic acid (1.61 g, 11.5 mmol, 1.2 eq.), Cs_2_CO_3_ (11.2 g, 34.5 mmol,

3.6 eq.) and 4-bromophenylpropionic acid (2.2 g, 9.6 mmol, 1.0 eq.) in dry degassed THF/water (100 mL, 9:1 v/v mixture) and the resultant mixture stirred at reflux for 8 h. The resultant solution was diluted with HCl (100 mL, 1 M) and extracted with EtOAc (3 x 50 mL) the combined organic phase was subsequently washed with brine, before being dried (MgSO_4_), filtered and concentrated under reduced pressure. The resultant acid was purified by flash column chromatography (silica, 1:1 v/v EtOAc:Hex) to give the title compound as a white crystalline solid (1.3 g, 54%).

^1^H NMR (300 MHz, DMSO-*d*_6_) δ 12.16 (br s, 1H), 7.58 – 7.16 (m, 8H), 2.88 (t, *J* = 7.8 Hz, 2H), 2.67 – 2.54 (m, 2H). ^13^C NMR (75 MHz, DMSO-*d*_6_) δ 173.7, 159.1 (d, ^1^*J*_CF_ = 245.7 Hz), 140.6, 132.8, 130.6 (d, ^4^*J*_CF_ = 6.0 Hz, 2C), 129.3 (d, ^3^*J*_CF_ = 7.9 Hz), 128.7 (2C), 128.5, 128.1 (d, ^3^*J*_CF_ = 12.3 Hz), 124.9 (d, ^4^*J*_CF_ = 3.6 Hz), 116.0 (d, ^2^*J*_CF_ = 22.6 Hz), 35.0, 30.0. ^19^F NMR (282 MHz, DMSO-*d*_6_) δ - 118.4. IR (diamond cell, neat) ϖ_max_: 3187, 1696, 1483, 1410, 1216, 1009, 940, 814, 755, 666, 566, cm^−1^. LRMS (ESI-) m/z: 243 [(M-H)^-^, 100%]

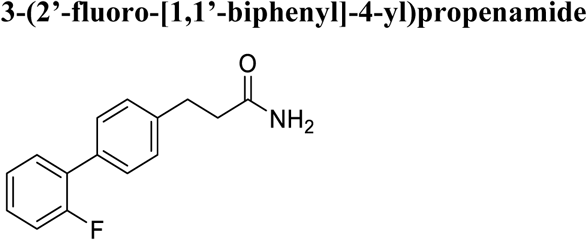

3-(2’-fluoro-[1,1’-biphenyl]-4-yl)propanoic acid (1.0 g, 4.1 mmol, 1.0 eq.) and 1,1’- carbonyldiimidazole (854 mg, 5.2 mmol, 1.26 eq.) were stirred for 1 h at room temperature in THF (4 mL) under a N_2_ atmosphere. The reaction was cooled on ice then aqueous ammonia (28%, 2.25 mL) was added. The reaction was stirred for 4 h, allowing the solution to warm to room temperature. The solvent was removed by rotary evaporation and the residue dissolved in dichloromethane (15 mL) and washed with aqueous sodium hydroxide (1 M, 5 mL), then aqueous hydrochloric acid (1 M, 5 mL) and then water (5 mL). The organic layer was dried (MgSO4), filtered and evaporated to dryness to yield the title compound as a white powder (607 mg, 61%).

^1^H NMR (300 MHz, DMSO-*d*_6_) δ 7.57 – 7.17 (m, 8H), 6.79 (s, 2H), 2.86 (t, *J* = 7.9 Hz, 2H), 2.41 (t, *J* = 7.9 Hz, 2H). ^13^C NMR (75 MHz, DMSO-*d*_6_) δ 173.3, 157.4 (d, ^1^*J*_CF_ = 246.3 Hz), 141.2, 132.6, 130.5 (d, ^4^*J*_CF_ = 7.9 Hz, 2C), 129.2 (d, ^3^*J*_CF_ = 8.3 Hz), 128.6 (2C), 128.4, 128.1 (d, ^3^*J*_CF_ = 12.1 Hz), 124.8, 116.0 (d, ^2^*J*_CF_ = 22.7 Hz), 36.4, 30.5. ^19^F NMR (282 MHz, DMSO) δ -118.4. IR (diamond cell, neat) ϖ_max_: 3400, 3180, 1650, 1482, 1412, 1009, 806, 754, 624 cm^−1^. LRMS (ESI+) m/z: 266 [(M+Na)^+^, 100%].

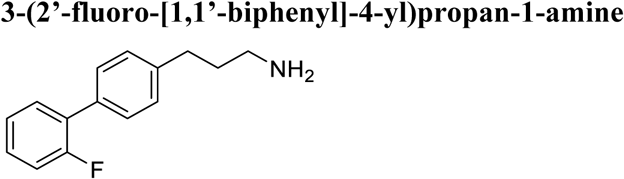

A solution of amide (500 mg, 2.1 mmol, 1.0 eq) in THF (8 mL) was treated with LiAlH_4_ (312 mg, 8.2 mmol, 3.9 eq) at 0 °C and stirred under a N_2_ atmosphere whilst warming to room temperature. After 2 h, the reaction was heated at reflux for 16 h and then cooled on ice. Chilled water (300 µL) was added dropwise, with vigorous stirring, and then followed by aqueous sodium hydroxide (15% w/v, 300 µL) and additional water (1 mL). The solution was left stirring at room temperature until effervescence had ceased and the grey powder had turned white (30 min). The solution was dried (MgSO_4_) and then filtered. The precipitate was washed with additional dichloromethane (2 x 10 mL). The filtrate in each case was combined, and solvent removed under reduced pressure. The crude oil thus obtained was purified by flash column chromatography (silica, 0.5:9.5 v/v MeOH (saturated with NH_3_):CH_2_Cl_2_) to give the title compound as a colourless wax (375 mg, 78%).

^1^H NMR (400 MHz, DMSO-*d*_6_) δ 7.60 – 7.12 (m, 8H), 4.17 (br s, 2H), 3.05 – 2.88 (m, 2H), 2.75 – 2.52 (m, 2H), 1.77 – 1.64 (m, 2H). ^19^F NMR (282 MHz, DMSO-*d*_6_) δ -118.4. IR (diamond cell, neat) ϖ_max_: 3334, 2923, 1611, 1481, 1314, 814, 751, 551cm^−1^. LRMS (ESI+) m/z: 230 [(M+Na)^+^, 100%].

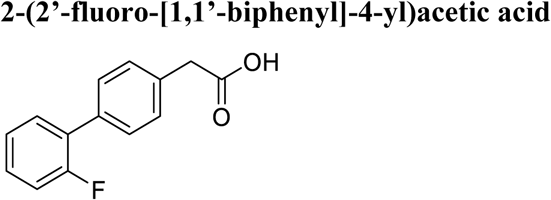

Tetrakis(triphenylphosphine)palladium(0) (554.7 mg, 0.48 mmol, 0.05 mol%) was added to a stirred suspension of 2-fluorophenylboronic acid (1.61 g, 11.5 mmol, 1.2 eq.), Cs_2_CO_3_ (11.2 g, 34.5 mmol, 3.6 eq.) and 4-bromophenylacetic acid (2.1 g, 9.6 mmol, 1.0 eq.) in dry degassed THF/water (100 mL, 9:1 v/v mixture) and the resultant mixture stirred at reflux for 8 h. The resultant solution was diluted with HCl (100 mL, 1 M) and extracted with EtOAc (3 x 50 mL) the combined organic phase was subsequently washed with brine, before being dried (MgSO_4_), filtered and concentrated under reduced pressure. The resultant acid was purified by flash column chromatography (silica, 1:1 v/v EtOAc:Hex) to give the title compound as a white crystalline solid (2.1 g, 95%).

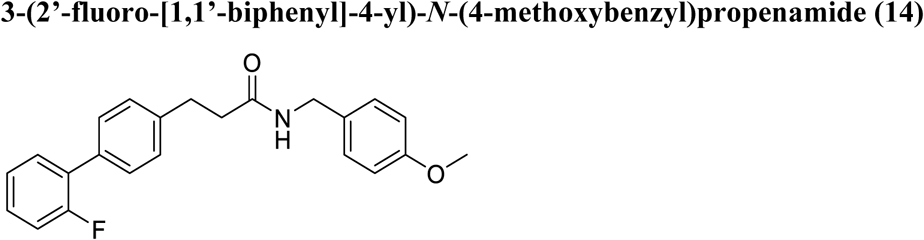

3-(2’-fluoro-[1,1’-biphenyl]-4-yl)propanoic acid (100 mg, 0.41 mmol, 1.0 eq.) and 1,1’- carbonyldiimidazole (85.4 mg, 0.52 mmol, 1.27 eq.) were stirred for 1 h at room temperature in THF (1 mL) under a N_2_ atmosphere. The reaction was cooled on ice then 4-aminomethyl phenol (67 mg, 0.49 mmol, 1.2 eq.) was added. The reaction was stirred for 4 h, allowing the solution to warm to room temperature. The solvent was removed by rotary evaporation and the residue dissolved in dichloromethane (15 mL) and washed with aqueous sodium hydroxide (1 M, 5 mL), then aqueous hydrochloric acid (1 M, 5 mL) and then water (5 mL). The organic layer was dried ( MgSO_4_), filtered and evaporated to dryness to yield the title compound as a white powder (111 mg, 74%).

R_f_: 0.37 (SiO_2_, 1:1 EtOAc:Hex), m.p.: 111 – 113 °C, ^1^H NMR (500 MHz, DMSO-*d*_6_) δ 8.27 (t, *J* = 5.9 Hz, 1H), 7.51 – 7.48 (m, 1H), 7.45 – 7.44 (m, 2H), 7.41 – 7.37 (m, 1H), 7.32 – 7.26 (m, 4H), 7.07 (d, *J* = 8.5 Hz, 2H), 6.82 (d, *J* = 8.7 Hz, 2H), 4.19 (d, *J* = 5.8 Hz, 2H), 3.68 (s, 3H), 2.90 (t, *J* = 7.6 Hz, 2H), 2.48 (t, *J* = 7.6 Hz, 2H). ^13^C NMR (126 MHz, DMSO-*d*_6_) δ 171.0, 159.1 (d, ^1^*J*_CF_ = 245.7 Hz), 158.1, 141.0, 132.7, 131.4, 130.6 (d, ^4^*J*_CF_ = 3.5 Hz, 2C), 129.3 (d, ^3^*J*_CF_ = 8.3 Hz), 128.6 (d, ^4^*J*_CF_ = 3.0 Hz), 128.6 (2C), 128.4 (2C), 128.1 (d, ^3^*J*_CF_ = 13.0 Hz), 124.9 (d, ^4^*J*_CF_ = 3.5 Hz), 116.1 (d, ^2^*J*_CF_ = 22.6 Hz), 113.6 (2C), 55.0, 41.4, 36.8, 30.8. ^19^F NMR (471 MHz, DMSO-*d*_6_) δ -118.4. IR (diamond cell, neat) ϖ_max_: 3297, 1632, 1511, 1483, 1244, 1217, 1175, 1106, 1037, 823, 754, 565, 515 cm^−1^. LRMS (ESI+) m/z: 386 [(M+Na)^+^, 100%].

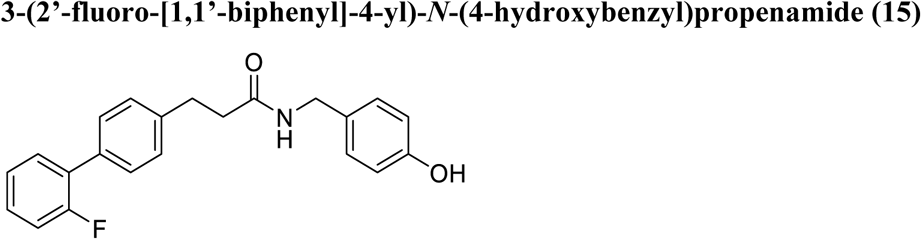

3-(2’-fluoro-[1,1’-biphenyl]-4-yl)propanoic acid (100 mg, 0.41 mmol, 1.0 eq.) and 1,1’- carbonyldiimidazole (85.4 mg, 0.52 mmol, 1.27 eq.) were stirred for 1 h at room temperature in THF (1 mL) under a N_2_ atmosphere. The reaction was cooled on ice then 4-aminomethyl phenol (61 mg, 0.49 mmol, 1.2 eq.) was added. The reaction was stirred for 4 h, allowing the solution to warm to room temperature. The solvent was removed by rotary evaporation and the residue dissolved in dichloromethane (15 mL) and washed with aqueous sodium hydroxide (1 M, 5 mL), then aqueous hydrochloric acid (1 M, 5 mL) and then water (5 mL). The organic layer was dried (MgSO4), filtered and evaporated to dryness to yield the title compound as a white powder (126 mg, 88%).

R_f_: 0.33 (SiO_2_, 1:1 EtOAc:Hex), m.p.: 153 – 155 °C ^1^H NMR (500 MHz, DMSO-*d*_6_) δ 9.25 (s, 1H), 8.23 (t, *J* = 5.8 Hz, 1H), 7.50 (td, *J* = 7.9, 1.7 Hz, 1H), 7.45 (dd, *J* = 8.2, 1.8 Hz, 2H), 7.43 – 7.36 (m, 1H), 7.33 – 7.28 (m, 4H), 6.99 (d, *J* = 8.4 Hz, 2H), 6.68 (d, *J* = 8.4 Hz, 2H), 4.15 (d, *J* = 5.8 Hz, 2H), 2.90 (t, *J* = 7.6 Hz, 2H), 2.47 (t, *J* = 7.6 Hz, 2H). ^13^C NMR (126 MHz, DMSO-*d*_6_) δ 171.0, 159.1 (d, ^1^*J*_CF_ = 245.6 Hz), 156.2, 141.1, 132.7, 130.6 (d, ^4^*J*_CF_ = 3.3 Hz, 2C), 129.6, 129.3 (d, ^3^*J*_CF_ = 8.4 Hz), 128.6 (d, *J* = 2.9 Hz), 128.5 (d, ^4^*J*_CF_ = 2.3 Hz), 128.2 (d, ^3^*J*_CF_ = 13.0 Hz), 124.9 (d, ^4^*J*_CF_ = 3.5 Hz), 116.1 (d, ^2^*J*_CF_ = 22.5 Hz), 114.9 (2C), 41.6, 36.8, 30.8. ^19^F NMR (471 MHz, DMSO-*d*_6_) δ -118.4. IR (diamond cell, neat) ϖ_max_: 3320, 1613, 1515, 1481, 1434, 1205, 1105, 841, 807, 761, 567, 498, 475 cm^−1^. LRMS (ESI+) m/z: 372 [(M+Na)^+^, 100%].

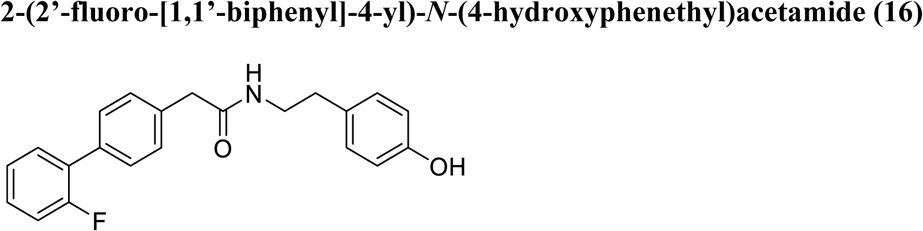

According to the general procedure for PyBOP^®^ amidation, the title compound was synthesised from 3-(2’-fluoro-[1,1’-biphenyl]-4-yl)acetic acid (200 mg, 0.86 mmol, 1.0 eq.) and 4-aminoethyl phenol (143 mg, 1.0 mmol, 1.16 eq.). The crude product was purified by flash column chromatography (silica, 1:1 v/v EtOAc:Hex) to give the title compound as a white crystalline solid (211 mg, 70%).

R_f_: 0.30 (SiO_2_, 1:1 EtOAc:Hex), m.p.: 83 – 87 °C, ^1^H NMR (500 MHz, DMSO-*d*_6_) δ 9.16 (s, 1H), 8.31 (s, 1H), 8.11 (t, *J* = 5.6 Hz, 1H), 7.51 (t, *J* = 7.9 Hz, 1H), 7.47 (d, *J* = 8.1 Hz, 1H), 7.42 – 7.38 (m, 1H), 7.32 (d, *J* = 8.1 Hz, 2H), 7.30 – 7.25 (m, 2H), 6.97 (d, *J* = 8.3 Hz, 2H), 6.68 (d, *J* = 8.4 Hz, 2H), 3.45 (s, 2H), 3.26 – 3.21 (m, 2H), 2.61 (t, *J* = 7.3 Hz, 2H). ^13^C NMR (126 MHz, DMSO-*d*_6_) δ 169.8, 159.1 (d, ^1^*J*_CF_ = 245.8 Hz), 155.6, 136.2, 133.1, 131.1 (d, ^2^*J*_CF_ = 30.0 Hz), 130.7 (d, ^4^*J*_CF_ = 3.4 Hz, 2C), 129.5 (2C), 129.3 (d, ^3^*J*_CF_ = 8.3 Hz), 129.2 (2C), 128.6 (d, ^4^*J*_CF_ = 2.8 Hz), 128.1 (d, ^3^*J*_CF_ = 13.2 Hz), 124.9 (d, ^4^*J*_CF_ = 3.6 Hz), 116.1 (d, ^2^*J*_CF_ = 22.5 Hz), 115.1 (2C), 79.2, 42.1, 40.7, 34.3. ^19^F NMR (471 MHz, DMSO-*d*_6_) δ -118.4. IR (diamond cell, neat) ϖ_max_: 3267, 2929, 1609, 1561, 1513, 1483, 1360, 1210, 813, 753, 580, 462 cm^−1^. LRMS (ESI+) m/z: 372 [(M+Na)^+^, 100%].

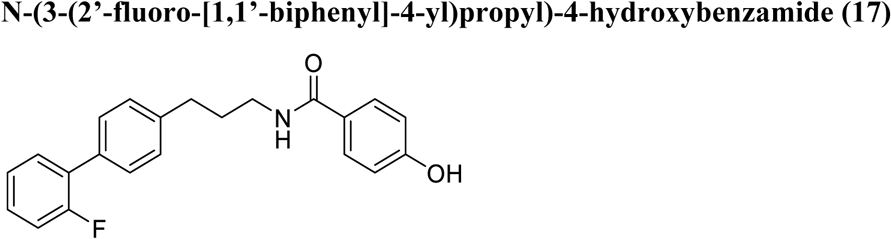

An ice cold magnetically stirred solution of 3-(2’-fluoro-[1,1’-biphenyl]-4-yl)propan-1-amine (200 mg, 0.87 mmol, 1.0 eq.), 4-hydroxybenzoic acid (141 mg, 1.04 mmol, 1.2 eq.) and *^i^*Pr_2_Net (303 μL, 1.74 mmol, 2.0 eq.) in DMF (5 mL) was treated with PyBOP^®^ (452 mg, 0.87 mmol, 1.0 eq.), allowed to warm to room temperature and stirring continued for 12 h. The reaction mass was diluted with CH_2_Cl_2_ (50 mL) and water (50 mL), the separated organic phase was subsequently washed with NaHCO_3_ (25 mL of a sat. aq. Solution) and brine (100 mL) before being dried (MgSO_4_), filtered and purified *via* flash column chromatography (silica, EtOAc).

^1^H NMR (400 MHz, DMSO-*d*_6_) δ 9.03 (dd, *J* = 2.3, 0.9 Hz, 1H), 8.71 – 8.67 (m, 2H), 8.21 – 8.17 (m, 2H), 7.64 – 7.14 (m, 10H), 3.35 – 3.29 (m, 2H), 2.72 – 2.62 (m, 2H), 1.93 – 1.83 (m, 2H). ^13^C NMR (101 MHz, DMSO-*d*_6_) δ 165.3, 159.6 (d, ^1^*J*_CF_ = 245.5 Hz), 152.2, 148.8, 141.9, 135.4, 133.1, 131.1 (d, ^4^*J*_CF_ = 3.5 Hz), 130.6, 129.7 (d, ^3^*J*_CF_ = 8.3 Hz), 129.4 (d, ^3^*J*_CF_ = 7.3 Hz), 129.2 (d, ^4^*J*_CF_ = 2.9 Hz), 129.1, 128.8 (d, ^3^*J*_CF_ = 4.6 Hz), 127.0 (d, ^2^*J*_CF_ = 11.6 Hz), 126.2, 125.3 (d, ^4^*J*_CF_ = 3.4 Hz), 123.9, 116.5 (d, ^2^*J*_CF_ = 22.7 Hz), 32.8 (2 overlapping signals), 31.1. ^19^F NMR (376 MHz, DMSO-*d*_6_) δ -118.4. IR (diamond cell, neat) ϖ_max_: 3302, 3027, 2948, 2465, 1626, 1588, 1544, 1481, 1448, 1431, 1406, 1362, 1317, 1211, 820, 757, 742, 708, 697, 622, 535 cm^−1^. LRMS (ESI-) m/z 348 [(M-H)^-^, 100%].

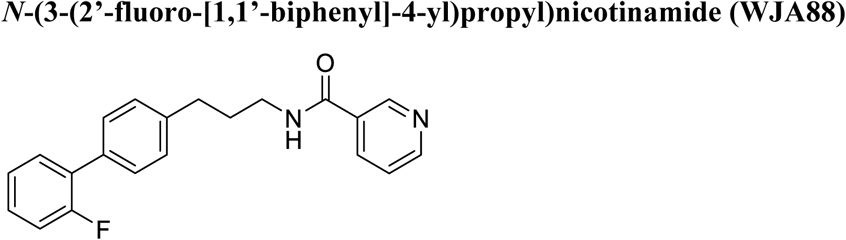

An ice cold magnetically stirred solution of 3-(2’-fluoro-[1,1’-biphenyl]-4-yl)propan-1-amine (200 mg, 0.87 mmol, 1.0 eq.), nicotinic acid (129 mg, 1.04 mmol, 1.2 eq.) and *^i^*Pr_2_Net (303 μL, 1.74 mmol, 2.0 eq.) in DMF (5 mL) was treated with PyBOP^®^ (452 mg, 0.87 mmol, 1.0 eq.), allowed to warm to room temperature and stirring continued for 12 h. The reaction mass was diluted with CH_2_Cl_2_ (50 mL) and water (50 mL), the separated organic phase was subsequently washed with NaHCO_3_ (25 mL of a sat. aq. Solution) and brine (100 mL) before being dried (MgSO_4_), filtered and purified *via* flash column chromatography (silica, EtOAc).

R_f_: 0.35 (SiO_2_, EtOAc), m.p.: 96 – 98 °C, ^1^H NMR (400 MHz, DMSO-*d*_6_) δ 9.01 (d, *J* = 1.5 Hz, 1H), 8.70 – 8.67 (m, 1H), 8.18 (dt, *J* = 7.9, 2.0 Hz, 1H), 7.53 – 7.46 (m, 4H), 7.42 – 7.26 (m, 5H), 3.36 – 3.29 (m, 2H), 2.70 (t, *J* = 7.6 Hz, 2H), 1.89 (p, *J* = 7.6 Hz, 2H). ^13^C NMR (101 MHz, DMSO-*d*_6_) δ 164.8, 159.1 (d, ^1^*J*_CF_ = 245.6 Hz), 151.7, 148.3, 141.4, 134.9, 132.6, 130.6 (d, ^4^*J*_CF_ = 3.4 Hz, 2C), 130.1, 129.3 (d, ^3^*J*_CF_ = 8.4 Hz), 128.71 (d, ^4^*J*_CF_ = 2.9 Hz), 128.68 (2C), 128.2 (d, ^3^*J*_CF_ = 13.2 Hz), 124.9 (d, ^4^*J*_CF_ = 3.6 Hz), 123.4, 116.0 (d, ^2^*J*_CF_ = 22.6 Hz), 32.3, 30.6. ^19^F NMR (282 MHz, DMSO-*d*_6_) δ -118.4. IR (diamond cell, neat) ϖ_max_: 3302, 3027, 2948, 2465, 1626, 1588, 1544, 1481, 1448, 1431, 1406, 1362, 1317, 1211, 820, 757, 742, 708, 697, 622, 535 cm^−1^. LRMS (ESI+) m/z 357 [(M+Na)^+^, 100%]. HRMS (ESI+) calcd for C_21_H_19_FN_2_NaO [M+Na]+, 357.13736; Found 357.13704.

## SUPPLEMENTAL INFORMATION

Supplementary data are provided as Supplementary Figure 1-8 and Supplementary Table 1-10.

## ACKNOWLEDGEMENTS

We thank Paul Brennan (University of Oxford, UK) for providing KDM5 and KDM6 inhibitors KDOAM-25, KDOPZ-36A, KDOBA-67a and KDOBA-97a. We thank Andrew Jenner (Bioanalytical Mass Spectrometry Facility at the Mark Wainwright Analytical Centre, University of New South Wales, Sydney) for technical and scientific support during the GC-MS analysis of cholesterol. We acknowledge Duan Ni (Charles Perkins Centre, University of Sydney) for intellectual input on patients’ data analyses. We thank Jessica Ho (Duke-NUS Medical School, Singapore) for guidance with the design of PRDM9 primers. We acknowledge the Sydney Mass Spectrometry Core Research Facility at the University of Sydney and thank the technical staff for the maintenance of the instruments and support of the Australian National Fabrication Facility (ANFF) – Materials Node. This study was supported by NHMRC grant APP2003150 to L.M., J.R.B and T.G.J; NHMRC grant APP1153961 to L.M., M.K. T.G.J.; Tour de Cure grant (RSP-370-FY2023) to L.M., and the Arto Hardy Family and Tour de Cure (FY2023-2025) grants to J.M.C. and E.T.C. E.K., B.C., R.H.A., J.R.S. and S.L.H were funded by the Australian Government Research Training Program (RTP) Scholarship.

## AUTHOR CONTRIBUTIONS

G.L.J.: conceptualization, methodology, validation, formal analysis, investigation, data curation, writing – review and editing, visualisation. E.G.K., B.C., J.R.S., W.D.P., R.H.A., A.R., M.H., D.C.I., T.C., T.Y.D., J.K., M.K., R.P.: methodology, investigation and validation. J.K.K.L., H.K., P.Y.: software, formal analysis, visualisation. W.T.J., A.P.M., J.R.B.: chemistry – conceptualisation, methodology, investigation, formal analysis. S.L.H., E.T.C., J.M.C.: methodology, investigation, formal analysis, and funding acquisition. L.L.: methodology and resources. G.G.N.: resources and supervision. B.W.D.: resources. E.G.: methodology, writing – review & editing. T.G.J.: methodology, supervision, funding acquisition. M.K.: conceptualisation, supervision, funding acquisition. A.S.D.: methodology, formal analysis, data curation, writing – review & editing, supervision. L.M.: conceptualisation, methodology, validation, data curation, writing – original draft, visualisation, supervision, project administration and funding acquisition.

---

**Supplementary Figure 1.**
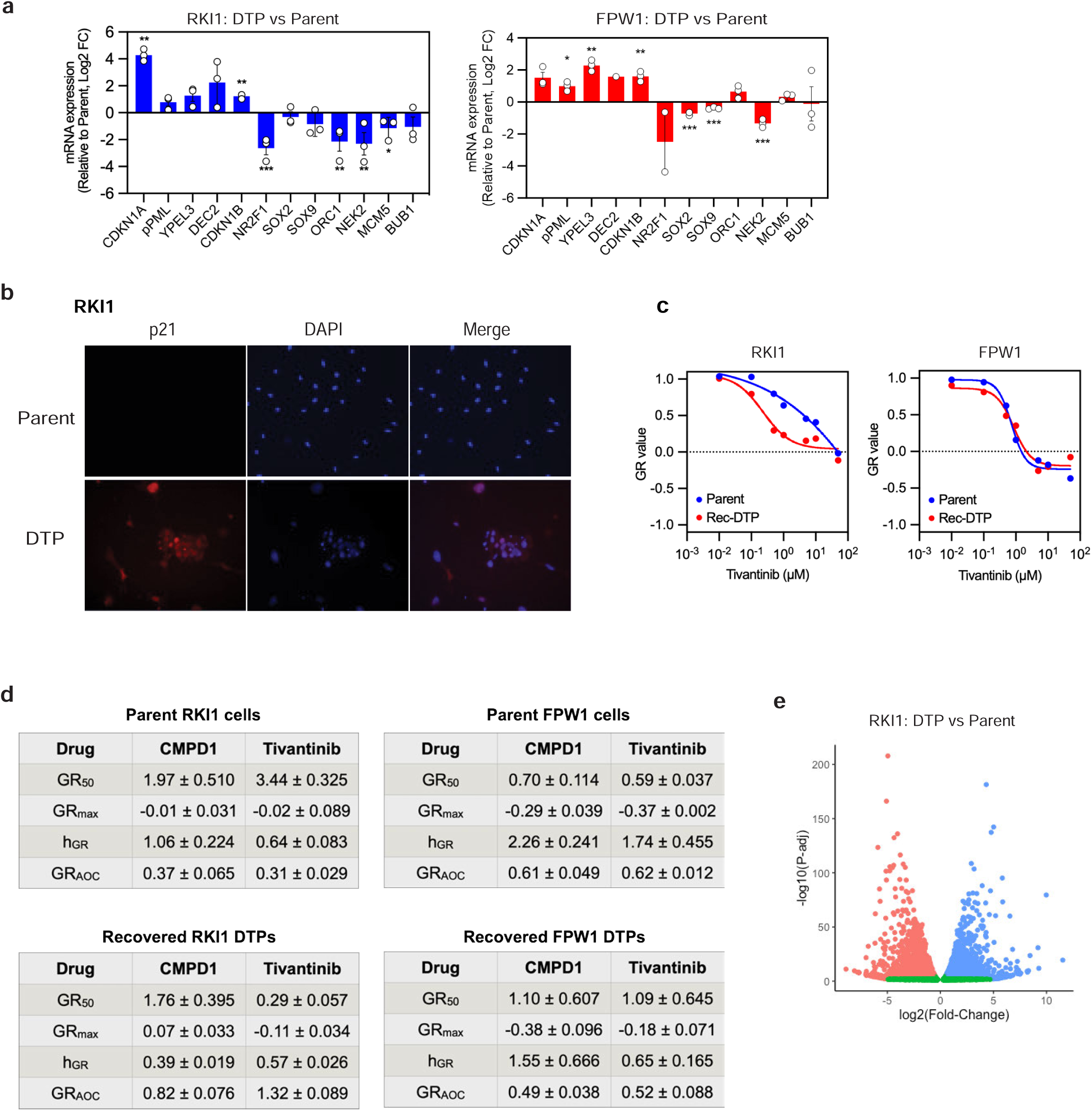
a) RT-qPCR analysis of indicated mRNAs in CMPD1 (25 μM, 14 days) derived drug-tolerant persister (DTP) cells compared to parent cells. Data are mean ± SD (n = 3; one sample t-test). b) Immunofluorescence imaging of p21 expression in parent and CMPD1 (25 μM, 14 days) derived drug-tolerant persister (DTP) cells. Images taken at 10X magnification. c) Tivantinib dose response curves (using growth rate (GR) values) in parent and recovered DTP cells. Data are mean of n = 3, corresponding GR metrics are in Supplementary Figure 1d. Rec-DTP cells were generated with tivantinib (25 µM, 14 days), followed by drug holiday. d) GR metrics for CMPD1 and tivantinib in parent and recovered DTP cells, values were calculated from dose-response curves in Figure 1c and Supplementary Figure 1c. e) Volcano plot of differentially expressed genes (DEGs, via RNA-seq of n = 3) in CMPD1 (25 μM, 14 days) derived drug-tolerant persister (DTP) cells compared to parent RKI1 cells. The x-axis represents the log2-fold change in normalised gene expression, the y-axis shows the -log10 of the adjusted P-value. Down-regulated DEGs are in red, upregulated DEGs in blue, non-significant genes in green.

**Supplementary Figure 2.**
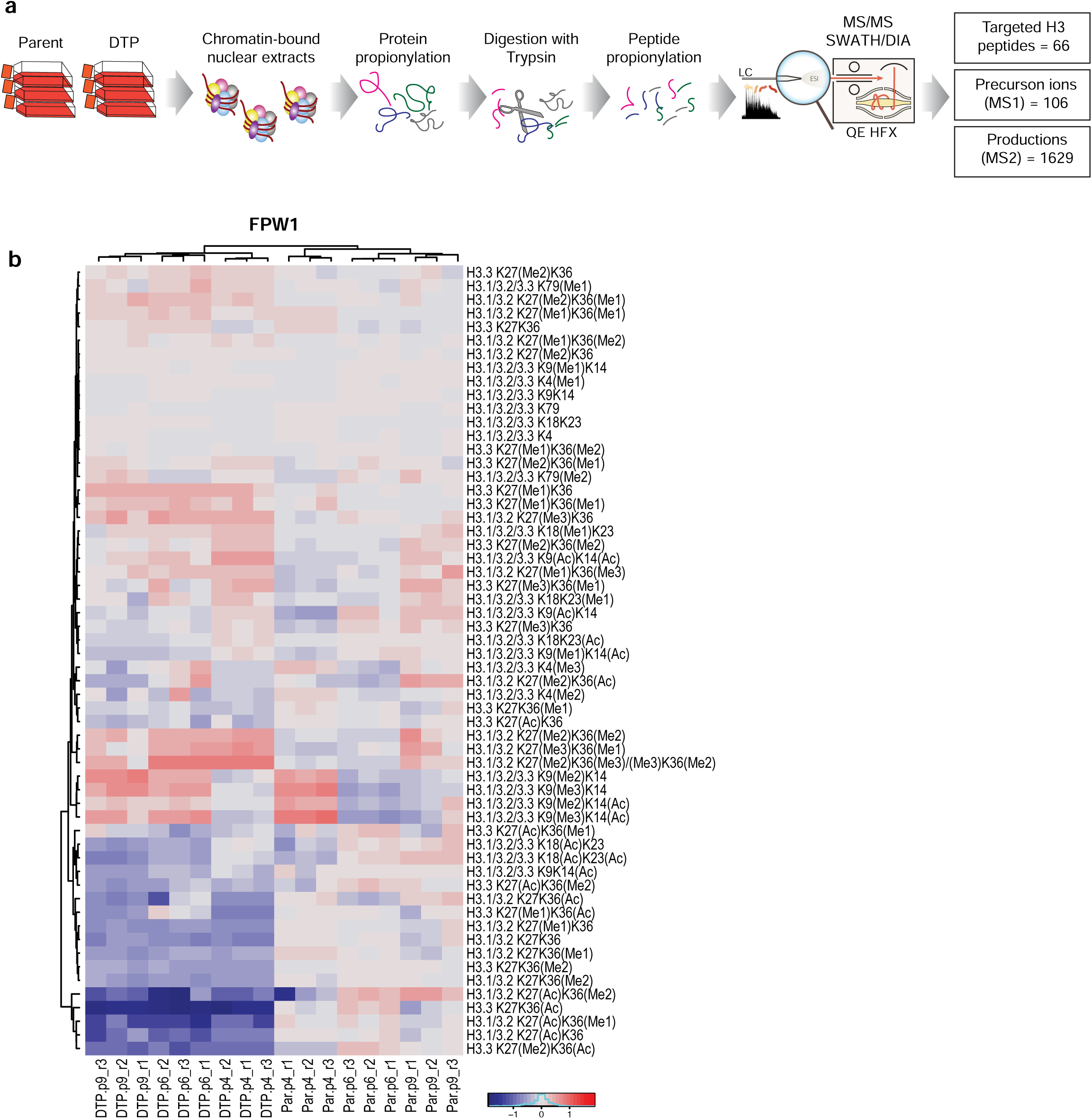
a) Schematic of mass spectrometry workflow for histone proteomics in Figure 2. b) Heatmap of peak areas for 66 unmodified, methylated and acetylated H3 peptides in FPW1 parent (Day 0) and CMPD1 (25 µM, 14 days) derived drug-tolerant persister (DTP) cells. Data are parent- DTP pairs of 3 independent experiments performed in triplicate.

**Supplementary Figure 3.**
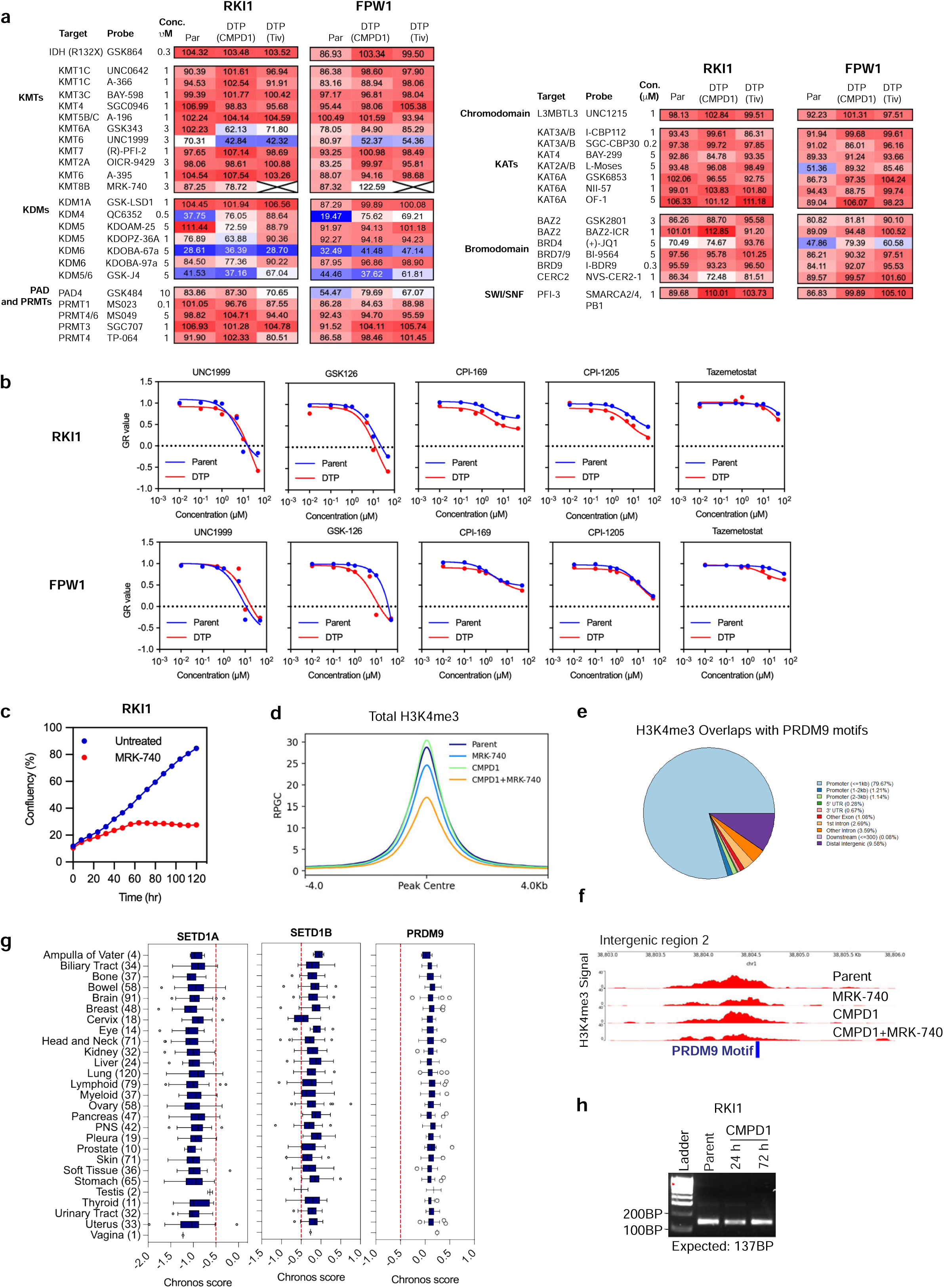
a) Drug-tolerant persister (DTP) cells were derived via treatment with CMPD1 or tivantinib (25 µM) for 14 days. Treatment-naïve parent (Par) and DTP cells were treated with epigenetic probes for 5 days and cell viability measured with CellTitre Blue. Heatmap displays mean values of cell viability at Day 5 relative to untreated cells (%). The SEM between biological repeats did not exceed 20% of the mean. b) Drug-tolerant persister (DTP) cells were derived from treatment with CMPD1 (25 µM) for 14 days. Parent and DTP cells were treated with KMT6 (EZH2) inhibitors for 5 days, cell viability measured with CellTitre Blue, growth-rate (GR) values were calculated using the online *GRcalculator* tool. Data are mean (n β 2). c) Incucyte SX5 Live-Cell imaging of RKI1 cells treated with MRK-740 (3 µM). Each datapoint is the average of confluency of nine images per well. d) Total H3K4me3 ChIPseq signal intensity (RPGC) in RKI1 cells treated with CMPD1 (10 µM) ± MRK-740 (3µM) for 3 days. e) Region annotations of H3K4me3 peaks overlapping with a PRDM9 motif, related to Figure 3f. f) Individual genome track of H3K4me3 intensity (RPGC) at an intergenic region aligning with PRDM9 motif (blue bar) in RKI1 cells treated with CMPD1 (10 µM) ± MRK-740 (3 µM) for 3 days. g) Chronos scores for SETD1A, SETD1B, and PRDM9 knockout in 1095 cancer cell lines categorised into tissue types with the number of cell lines in parentheses. Data sourced using the “23Q2 + Score” dataset from the DepMap portal. h) Agarose gel images of amplicon product for custom-designed PRDM9 primer.

**Supplementary Figure 4.**
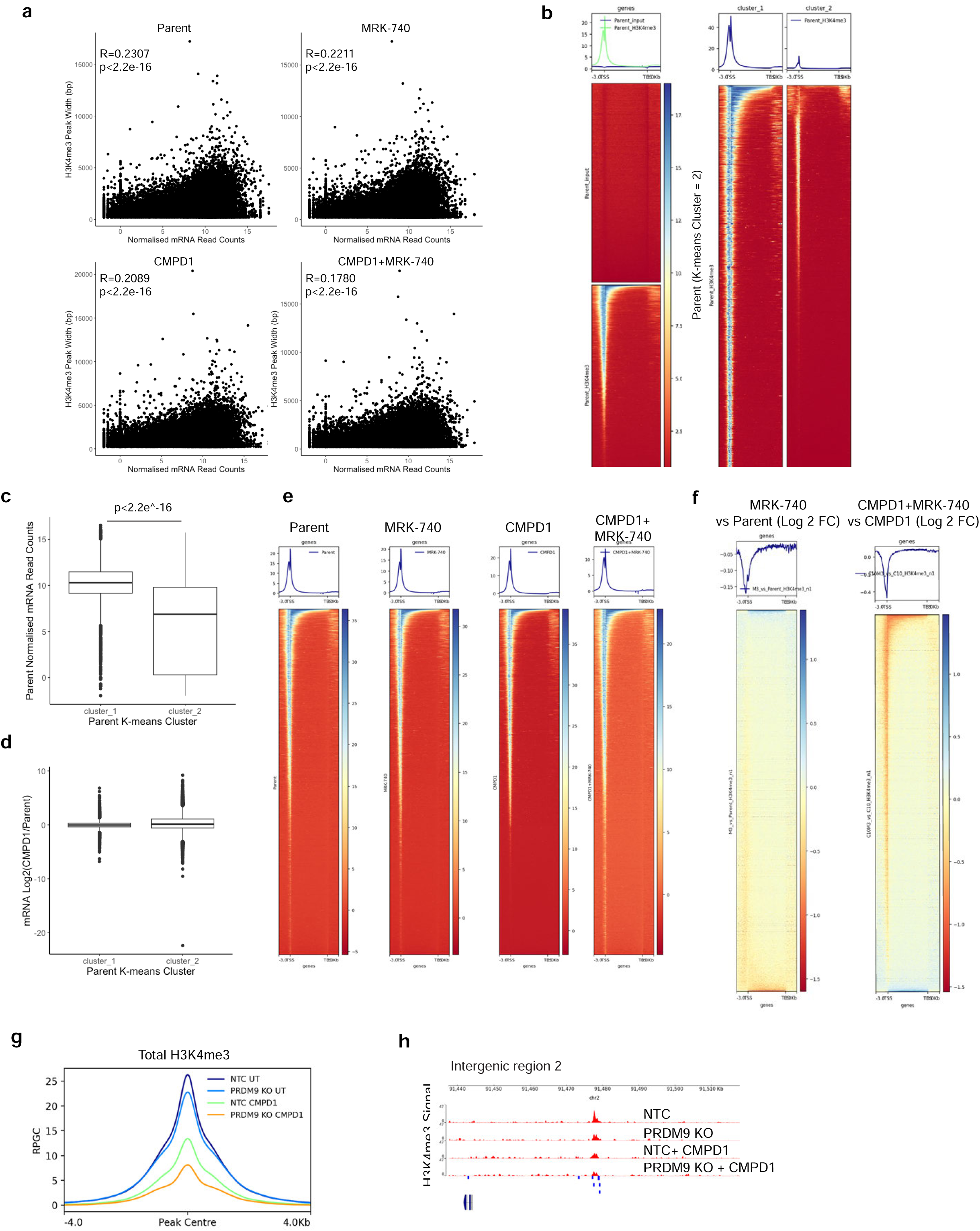
a) Scatter plots showing correlation between H3K4me3 peak width and normalised transcript counts (rlog). Pearson’s product moment correlation coefficient was used to determine R value and statistical significance. b) Genome wide H3K4me3 enrichment heatmaps for parent RKI1 cells divided into two K-means clusters. c) mRNA levels for genes in Clusters 1 and Cluster 2 from Supplementary Fig 2b. Wilcoxon rank- sum test was used to calculate P-value. d) mRNA changes (Log2 FC) for Cluster 1 and Cluster 2 genes in CMPD1 (10 µM, 3 days) treated vs parent RKI1 cells. e -f) Genome wide heatmaps for H3K4me3 signal (± 3kb from transcription start/end site) in RKI1 cells treated with CMPD1 (10 µM) ± MRK-740 (3µM) for 3 days. g) Total H3K4me3 ChIPseq signal density (RPGC) in CMPD1 (10 µM, 3 days) treated RKI1 cells transduced with NTC sgRNA (NTC) or PRDM9 sgRNA (PRDM9 KO). h) Individual genome track of H3K4me3 intensity (RPGC) at an intergenic region aligning with PRDM9 motif (blue bar) in CMPD1 (10 µM, 3 days) treated RKI1 cells transduced with NTC sgRNA (NTC) or PRDM9 sgRNA (PRDM9 KO).

**Supplementary Figure 5.**
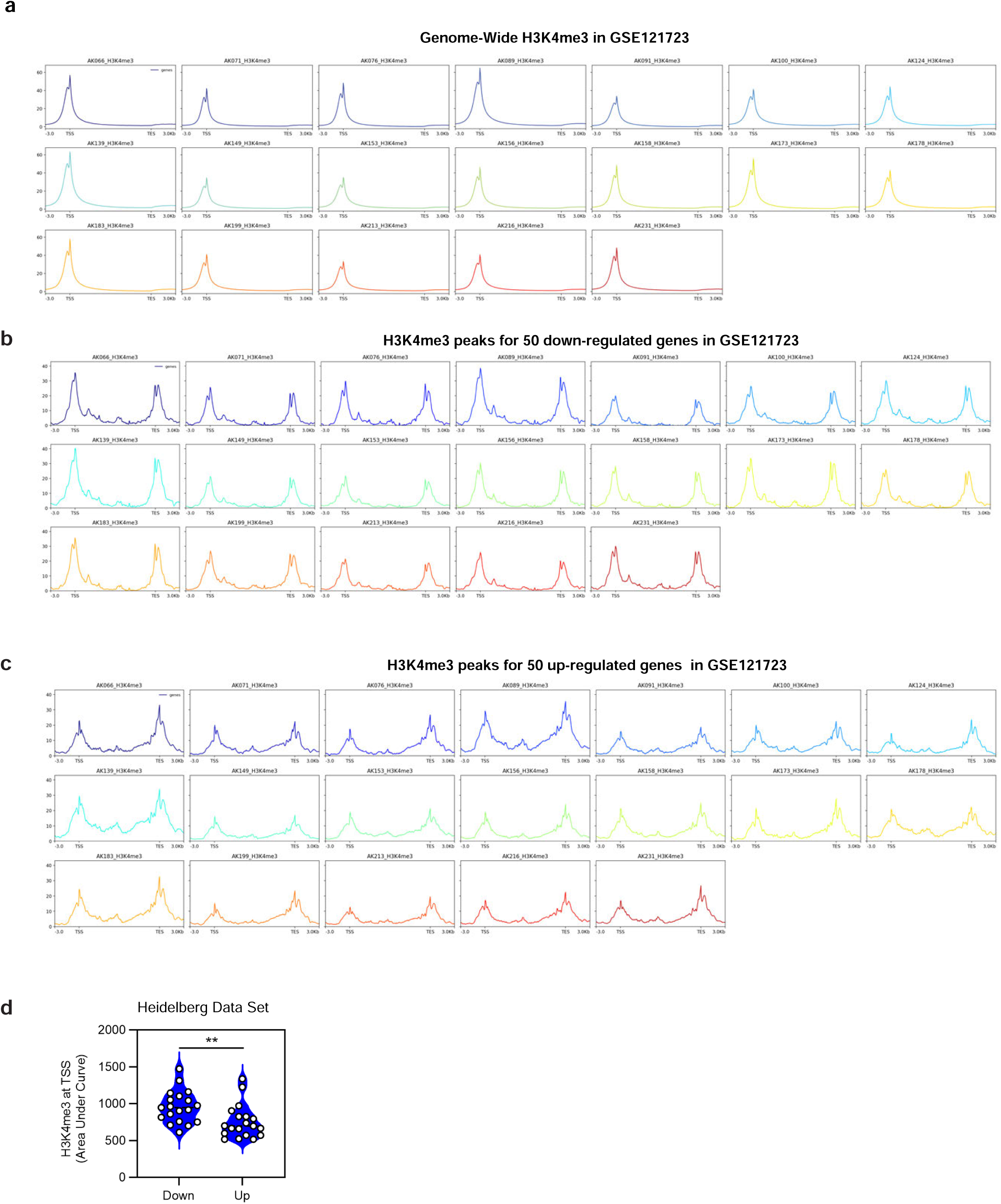
a) Genome wide H3K4me3 intensity (RPKM; ± 3kb from transcription start/end site) in 19 glioblastoma specimens (GSE121723). b) H3K4me3 signal intensity (RPKM; ± 3kb from transcription start/end site) for down-regulated genes (listed in Figure 4a) in in 19 glioblastoma specimens (GSE121723). c) H3K4me3 signal intensity (RPKM; ± 3kb from transcription start/end site) for up-regulated genes in 19 glioblastoma specimens (GSE121723). d) Quantification of H3K4me3 peaks at the transcription start sites (TSS, ± 3kb from transcription start site) of down-regulated and up-regulated genes (identified in RKI1 cells: CMPD1+MRK-740 vs CMPD1) in 19 glioblastoma specimens (GSE121723).

**Supplementary Figure 6.**
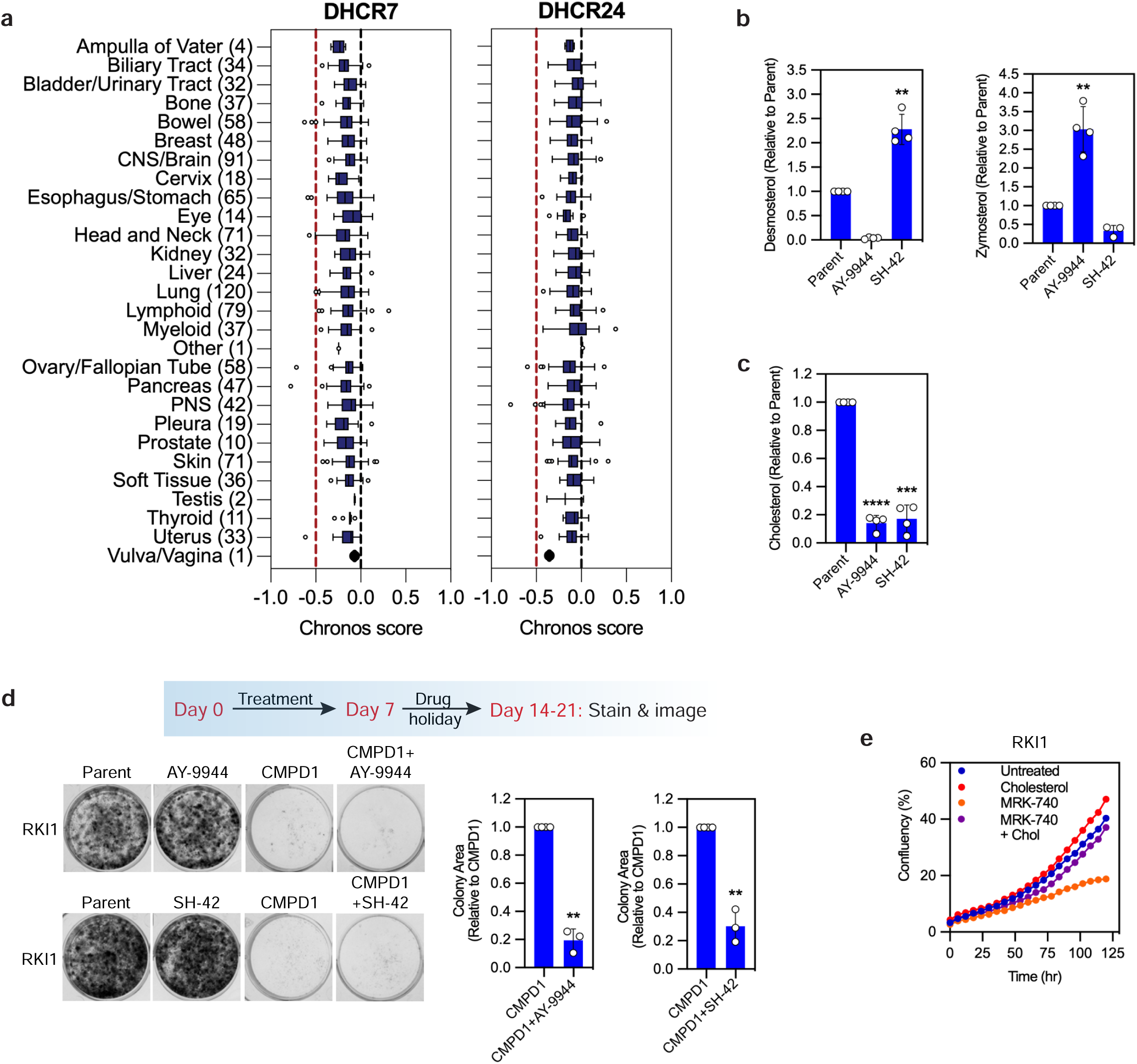
a) Chronos scores for DHCR7 and DHCR24 knockout in 1,095 cancer cell lines categorised into tissue types with the number of cell lines in parentheses. Data sourced using the “23Q2 + Score” dataset from the DepMap portal. b - c) GC-MS quantification of desmosterol, zymosterol and free cholesterol in RKI1 cells treated with AY-9944 (1 μM) or SH-42 (1 µM) for 3 days. Data are mean ± SD (n = 4; one sample t-test). d) Representative images and quantification of colony formation assays with RKI1 cells treated with CMPD1 (25 µM) ± AY-9944 or SH-42 (1 μM) for 7 days, followed by recovery in drug-free media until visible colonies formed in CMPD1-only treatments. Data are mean ± SD (n = 3; one sample t- test). e) Incucyte SX5 Live-Cell imaging of RKI1 cells treated with MRK-740 (3 µM) ± cholesterol complexed to methyl-beta-cyclodextrin (100 μg/mL, cholesterol component is 5% w/w). Each datapoint is the average of confluency in sixteen images per well.

**Supplementary Figure 7.**
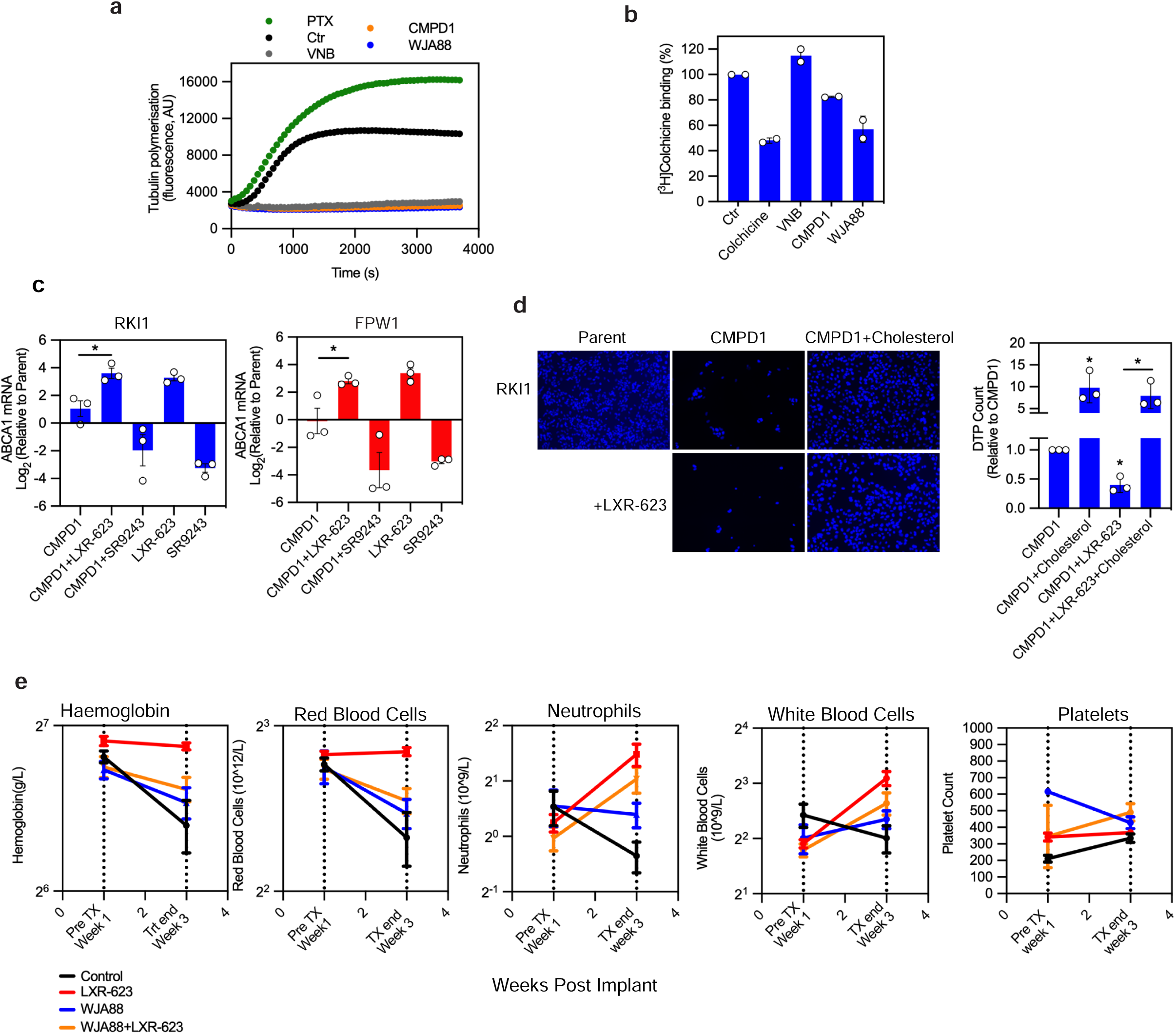
a) Kinetics of tubulin polymerisation in the presence of paclitaxel (PTX, 3 µM), vinblastine (VNB, 3 μM), CMPD1 (50 μM) and WJA88 (50 μM). Data are mean (n = 3 in duplicate). b) Tubulin binding assay measuring displacement of [^3^H]colchicine by colchicine (10 μM), vinblastine (VNB, 10 μM), CMPD1 (10 μM) and WJA88 (10 μM). Data are mean (n = 2). c) RT-qPCR of ABCA1 mRNA expression in RKI1 and FPW1 cells treated with CMPD1 (25 µM) ± LXR-623 (1 µM) or SR9243 (1 µM) for 72 hours. Data are mean ± SD (n = 3; unpaired t-test). d) DAPI-stained images of RKI1 cells treated with CMPD1 (25 µM) ± LXR-623 (1 µM) ± cholesterol complexed to methyl-beta-cyclodextrin (100 μg/mL, cholesterol component is 5% w/w) for 14 days. Quantification of drug-tolerant persisters (DTP) shown as bar graph. Data are mean ± SD (n = 3, one sample t-test compared to CMPD1-only, unpaired t-test comparing between samples). e) Toxicity was monitored immediately before and after the treatment. Blood counts (n=3 per group) from whole blood with ethylenediamine tetra-acetic acid were measured using a Forcyte Hematology Analyzer (Oxford Scientific). Shown are haemoglobin, red blood cells, neutrophils, white blood cells and platelets.

**Supplementary Figure 8.**
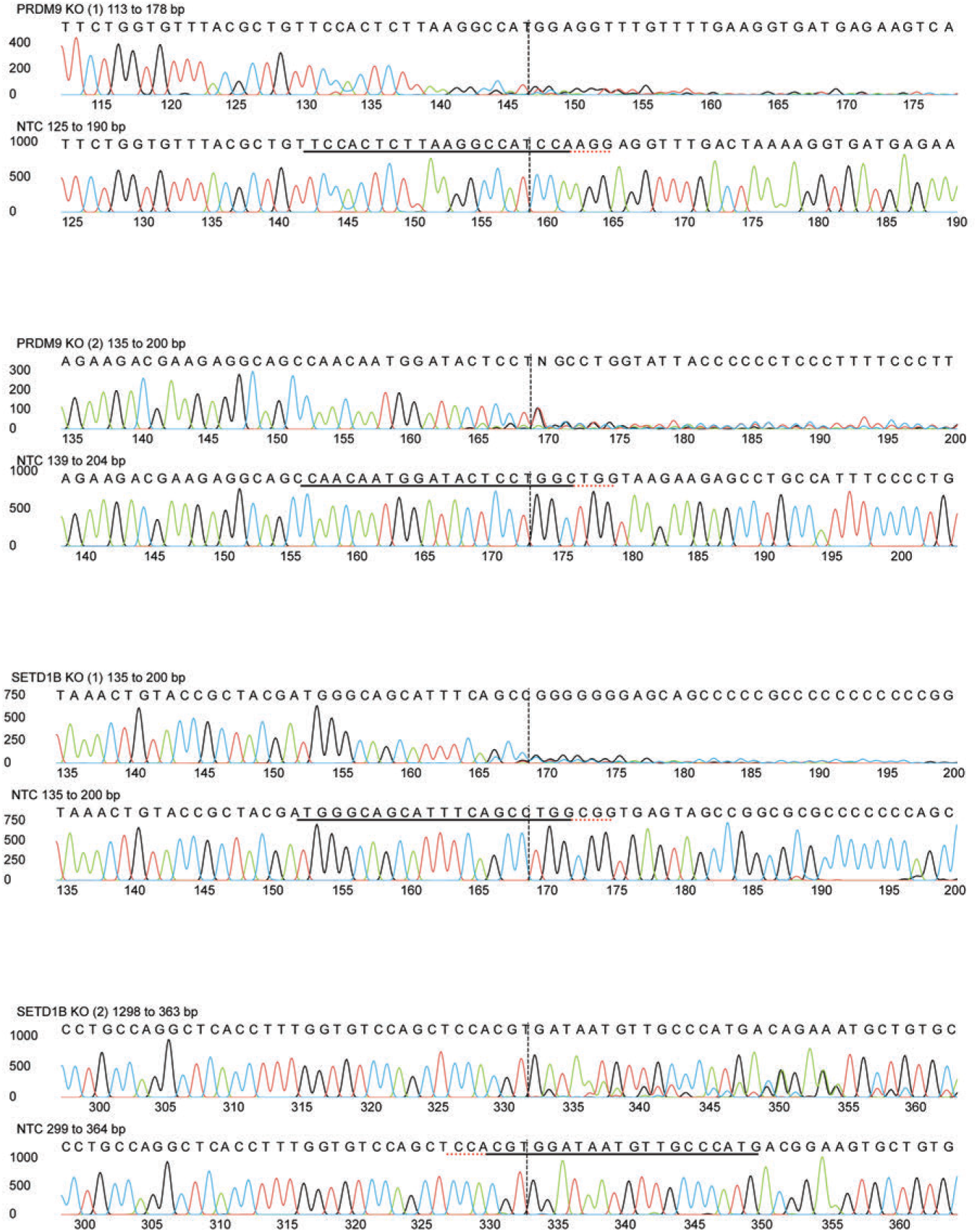

**Supplementary Table 1– 7.** will be provided in Browser Extensible Data format as Supplementary Information.

**Supplementary Table 8.**
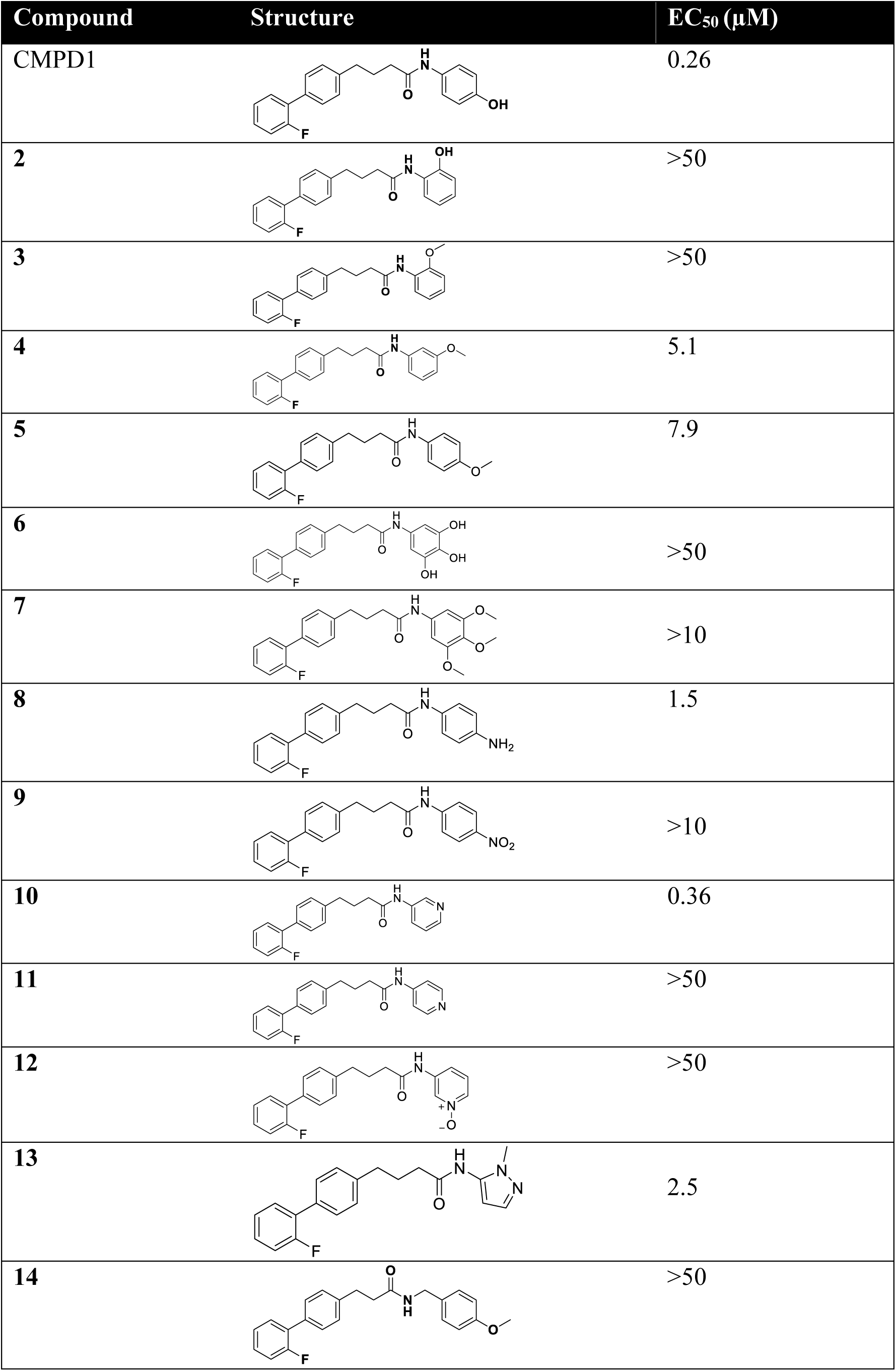
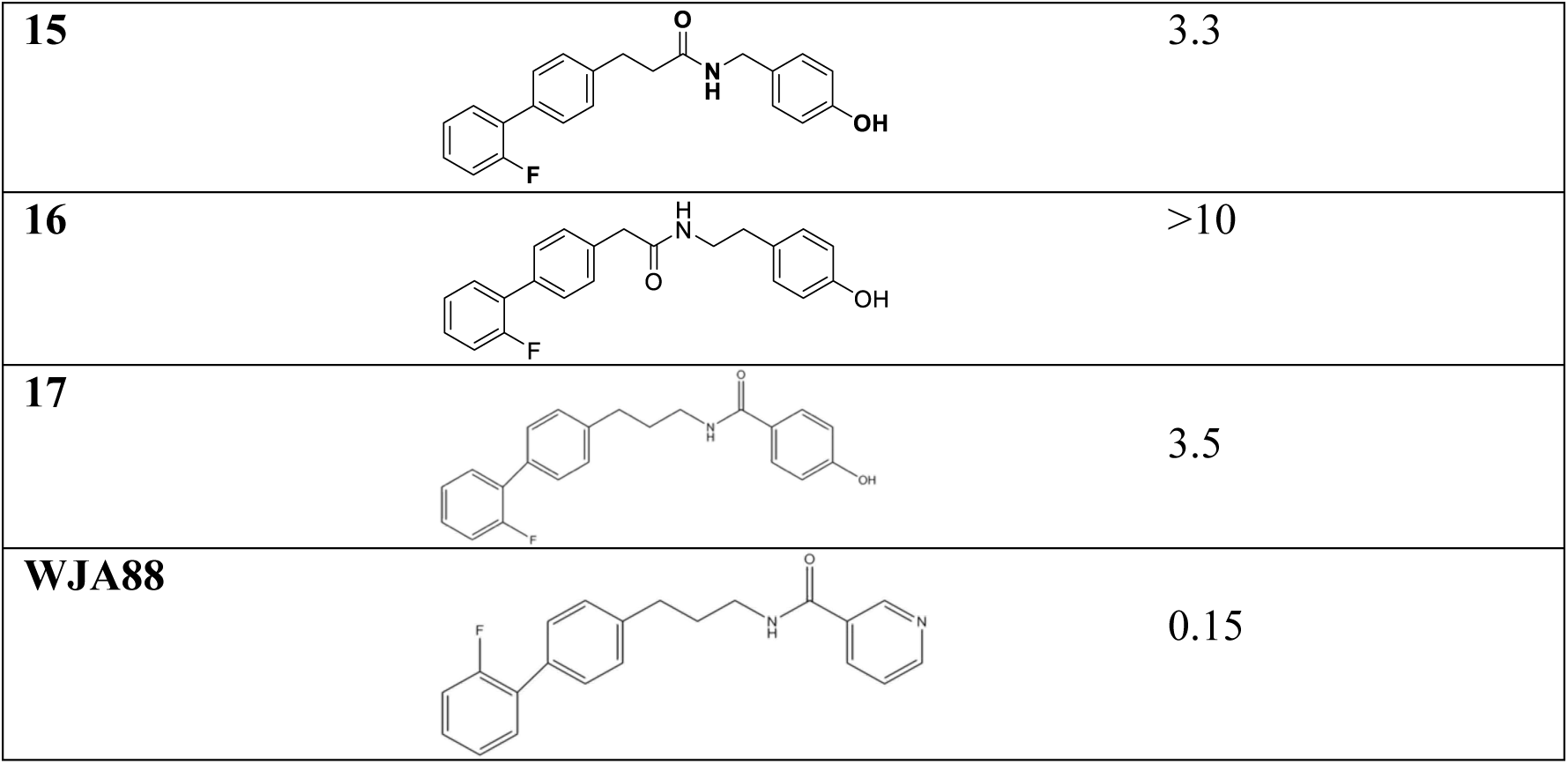
Cellular efficacy (EC_50_; n = 2 - 7) of CMPD1, analogues **1** - **17** and WJA88 in A172 cells.

| Compound | Structure | EC<sub>50</sub> (μM) |
| --- | --- | --- |
| CMPD1 |  | 0.26 |
| 2 |  | >50 |
| 3 |  | >50 |
| 4 |  | 5.1 |
| 5 |  | 7.9 |
| 6 |  | >50 |
| 7 |  | >10 |
| 8 |  | 1.5 |
| 9 |  | >10 |
| 10 |  | 0.36 |
| 11 |  | >50 |
| 12 |  | >50 |
| 13 |  | 2.5 |
| 14 |  | >50 |
| **15** |  | 3.3 |
| **16** |  | >10 |
| **17** |  | 3.5 |
| **WJA88** |  | 0.15 |

**Supplementary Table 9.** Cellular efficacy (EC_50_; n = 2) of CMPD1, analogue **10** and WJA88 in glioblastoma stem cell lines.

| Cell line | CMPD1<br>EC<sub>50</sub> (μM) | Analogue 10<br>EC<sub>50</sub> (μM) | WJA88<br>EC<sub>50</sub> (μM) |
| --- | --- | --- | --- |
| **MMK1** | 0.1 | 0.7 | 0.29 |
| **HW1** | 0.60 | 0.80 | 0.76 |
| **SB2** | 0.29 | 0.21 | 0.25 |
| **PB1** | 0.51 | 0.56 | 0.39 |
| **RN1** | 1.56 | 1.49 | 0.6 |
| **WK1** | 0.60 | 0.83 | 0.66 |
| **RKI1** | 0.9 | 0.73 | 0.6 |

**Supplementary Table 10.** Pharmacokinetic parameters of WJA88 after single 50 mg/kg dose administered intraperitoneally to male CD-1 mice (n = 3, data are mean).

| **WJA88** |  |
| --- | --- |
| **C<sub>max</sub> (ng/mL)** | 7907 |
| **T<sub>max</sub> (h)** | 0.25 |
| **T<sub>1/2</sub> (h)** | 1.19 |
| **T<sub>last</sub> (h)** | 8.00 |
| **AUC<sub>0-last</sub> (ng.h/mL)** | 4262 |
| **AUC<sub>0-inf</sub> (ng.h/mL)** | 4271 |
| **MRT<sub>0-last</sub> (h)** | 0.883 |
| **MRT<sub>0-inf</sub> (h)** | 0.905 |
| **AUC<sub>Extra</sub> (%)** | 0.263 |
| **AUMC<sub>Extra</sub> (%)** | 3.00 |

