## Supplementary Table 11 for "Histone methyltransferase PRDM9 promotes survival of drug-tolerant persister cells in glioblastoma"

| **Compound Name** | **Company** | **Catalogue Number** |
| --- | --- | --- |
| CMPD1 | Santa Cruz | Cat# Sc-203138, CAS: 41179-33-3 |
| Colchicine | Tocris | Cat# 1364, CAS: 64-86-8 |
| Nocodazole | Tocris | Cat# 1228, CAS: 31430-18-9 |
| Tivantinib | Selleckchem | Cat# S2753, CAS: 905854-02-6 |
| Paclitaxel | Tocris | Cat# 1097, CAS: 33069-62-4 |
| Vinblastine | Tocris | Cat# 1256, CAS:143-67-9 |
| Ixabepilone | AdooQ Bioscience | Cat# A11449; CAS: 219989-84-1 |
| Verapamil | Sigma-Aldrich | Cat# V4629; CAS: 152-11-4 |
| MK571 | Sigma-Aldrich | Cat# M7571; CAS: 115103-85-0 |
| CP-100356 | Sigma-Aldrich | Cat# PZ0171; CAS: 142715-48-8 |
| Elacridar | Tocris | Cat# 4646; CAS: 143851-98-3 |
| Zosuquidar | Tocris | Cat# 5456; CAS: 167465-36-3 |
| SCG Probe Set (inhibitors of epigenetic readers, writers and erasers) | Cayman Chemicals | Cat# 17748 |
| CPI-169 | Cayman Chemicals | Cat# 18299; CAS: 1450655-76-1 |
| CPI-1205 | AdooQ Bioscience | Cat# A16357; CAS: 1621862-70-1 |
| EPZ6438 (Tazemetostat) | Cayman Chemicals | Cat# 16174; CAS: 1403254-99-8 |
| GSK126 | Cayman Chemicals | Cat# 15415; CAS: 1346574-57-9 |
| CTSL1inhibitor I | MerckMillipore | Cat# 219421; CAS: 108005-94-3 |
| MRK-740 | MedChemExpress | Cat # HY-114209 |
| Ro-3306 | Cayman Chemicals | Cat# 15149 |
